# SeqDistK: a Novel Tool for Alignment-free Phylogenetic Analysis

**DOI:** 10.1101/2021.08.16.456500

**Authors:** Xuemei Liu, Wen Li, Guanda Huang, Tianlai Huang, Qingang Xiong, Wen Chen, Li C. Xia

## Abstract

Algorithms for constructing phylogenetic trees are fundamental to study the evolution of viruses, bacteria, and other microbes. Established multiple alignment-based algorithms are inefficient for large scale metagenomic sequence data because of their high requirement of inter-sequence correlation and high computational complexity. In this paper, we present SeqDistK, a novel tool for alignment-free phylogenetic analysis. SeqDistK computes the dissimilarity matrix for phylogenetic analysis, incorporating seven k-mer based dissimilarity measures, namely d2, d2S, d2star, Euclidean, Manhattan, CVTree, and Chebyshev. Based on these dissimilarities, SeqDistK constructs phylogenetic tree using the Unweighted Pair Group Method with Arithmetic Mean algorithm. Using a golden standard dataset of 16S rRNA and its associated phylogenetic tree, we compared SeqDistK to Muscle – a multi sequence aligner. We found SeqDistK was not only 38 times faster than Muscle in computational efficiency but also more accurate. SeqDistK achieved the smallest symmetric difference between the inferred and ground truth trees with a range between 13 to 18, while that of Muscle was 62. When measures d2, d2star, d2S, Euclidean, and k-mer size k=5 were used, SeqDistK consistently inferred phylogenetic tree almost identical to the ground truth tree. We also performed clustering of 16S rRNA sequences using SeqDistK and found the clustering was highly consistent with known biological taxonomy. Among all the measures, d2S (k=5, M=2) showed the best accuracy as it correctly clustered and classified all sample sequences. In summary, SeqDistK is a novel, fast and accurate alignment-free tool for large-scale phylogenetic analysis. SeqDistK software is freely available at https://github.com/htczero/SeqDistK.

## 1 Introduction

Phylogenetic analysis is the cornerstone of evolutionary biology and taxonomy. In molecular phylogenetic analysis, phylogenetic trees are constructed from comparing a group of homologous DNA or protein sequences **[1-2]**. Canonically, there are four steps in the analysis: (1) obtaining homologous sequence data, (2) determining the evolutionary distance, (3) performing multi-sequence alignment, and (4) building the phylogenetic tree. Global multiple alignment is the long-time standard for computing phylogenetic distances between sequences. Many phylogenetic tools were developed based on multi-sequence alignment since the 1970s and were applied in many studies, for examples see refs **[3-5]**.

The arrival of low-cost high-quality next generation sequencing (NGS) has led to a significant increase of the size of whole genome and metagenome sequencing data. Because of the high computation burden associated with multiple alignment, the subjectivity of its scoring function, and the high requirement on sequence relatedness, established phylogenetic algorithms can no longer meet the new computational challenges arising from those massive NGS datasets. It thus encouraged the development of alignment-free tools for comparative biological sequence analysis. Different to its alignment based counterpart, the alignment-free phylogenetic process is like follows: (1) transforming each sequence into a multiset of its building subsequence (e.g., base, amino acid, or **k-mer**); (2) calculating the dissimilarity between these subsequence multisets using dissimilarity measures; (3) building the phylogenetic tree with a tree algorithm and the computed dissimilarity measures as evolutionary distance.

A subsequence of size k is called k-mer. Researchers have extensively studied the statistical property of k-mers and the dissimilarity measures derived from them. These measures were proposed to study the evolutionary relationship between genomes, to assemble the fragments in metagenome samples, and to compare microbial communities **[6-8]**. For instance, *Pride et al* used tetra nucleotide frequency to infer microbial genome distances **[9]**. *Miller et al* clustered expressed human gene sequence using k-mer frequency consensus **[10]**. Those studies demonstrated that k-mer derived statistics is conserved within one organism’s genome but different between organisms. This organismal conservation of k-mer makes it an efficient dissimilarity measure for phylogenetic analysis.

K-mer measures are also preferable for large-scale phylogenetic analysis because of high computational efficiency. For instances, *Huan et al*. used k-mer based approach to perform assembly and phylogenetic analysis and they demonstrated that the runtime for computing the k-mer measures is significantly lower that of deriving Maximum Likelihood and Bayesian based distances with multiple alignment **[11]**. *Chan et al* also demonstrated that the alignment-free dissimilarity computation was 140 times faster than that of alignment-based **[12]**. Moreover, parallel computing methods were available to further accelerate the alignment-free dissimilarity measure calculations **[13]**.

Despite of these desirable qualities of k-mer based statistics and dissimilarity measures, their adoption in phylogenetic analysis is still in early stage. Among the existing works, *Qi et al*, then *Xu et al*, developed the CV-Tree, a web service that infers the microbial tree-of-life based on coding sequences **[14-15]**. CV-Tree employed a k-mer based evolutionary distance developed by Hao et al. **[16]** (abbreviated as *Hao*), which is a cosine distance of k-mer vector with (k-1)-mer background subtracted. There are also k-mer natural vector based approaches - a statistical variant involving position-aware multisets of k-mers. E.g., *Wen et al* first applied k-mer natural vector to construct phylogenetic trees using whole or partial genomes, while *Huang et al* further proposed an ensemble statistic **[17-18]**.

It is evident that there is still a lack of consensus on which dissimilarity measure to use in current alignment-free phylogenetics. There is also the lack of an integrative analysis tool incorporating key k-mer dissimilarity measures. This study addresses these deficiencies. By far, the most studied normalized k-mer measure is d2, which is the count of all exact k-mer matches between two sequences, summing over all k-mers by a given k **[19]**. Other normalized measures derived from d2, such as d2S and d2star, also had good results in clustering genome sequences hierarchically **[20]**. For instances, *Ahlgren et al* **[21]** made a systematic assessment of k-mer measures of viral and host genomes such as d2star and d2S, as well as *Hao*, Euclidean (Eu), Manhattan (Ma) and Chebyshev (Ch) distances, and found they were highly predictive of virus’ infection potential to hosts. In another study, *Chan et al* while inferring the phylogenetic tree from 4156 nucleotide sequence, found that the topology structure obtained using d2S was highly consistent with that obtained by multiple alignment **[12]**.

In view of the previous excellent performance from these k-mer dissimilarity measures and the need for an integrative and intuitive tool, we developed SeqDistK, a novel tool for alignment-free phylogenetic analysis. It can perform efficient calculation for seven key k-mer measures: Ch, Ma, Eu, d2, d2S, d2star, and Hao. Using SeqDistK, we extensively compared the performance of these measures with that of Muscle - a multiple alignment phylogenetic tool, using a standard 16S RNA dataset with known groud truth. We validated that the symmetric differences between the inferred trees and the ground truth were between 13 to 18 in most cases and much smaller than Muscle derived trees. We identified the d2S measure with k=5 and M=2 (using 5-mer and 2^nd^ order Markovian background subtraction) as the best measure for phylogenetic inference achieving the smallest symmetric difference. We have made the software of SeqDistK completely open source at https://github.com/htczero/SeqDistK.

## 2 Methods

### 2.1 The Workflow of SeqDistK

As shown in **Fig. 1A**, SeqDistK constructs a phylogenetic tree from input sequences in four steps: (1) First, SeqDistK counts k-mer frequency in each input sequence. Efficient counting of k-mer frequency is the premise for all k-mer based dissimilarity measures. Recent years have seen rapid development in methods to index and count k-mer frequency. Given the fact that k-mer based statistics were mostly useful for phylogenetic analysis when k is relatively small (<16), we implemented in SeqDistK a mature and simple algorithm to count k-mers frequency (**Fig.1B**). We mathematically transformed k-mer to an index can randomly address and operate an array-based memory storage efficiently. (2) Secondly, we record the k-mer frequency into a 4^*k*^ vector for each input sequence. If N input sequences were to be compared, their merged vectors are stored in a 4^*k*^ *N* matrix. (3) Thirdly, a user specifies the desired dissimilarity measure(s), which SeqDistK will use to calculate the distance matrix, which is a N-by-N matrix. (4) Finally, with this distance/dissimilarity matrix, SeqDistK employs the Unweighted Pair Group Method with Arithmetic Mean (UPGMA) to construct the phylogenetic tree. At the last step, SeqDistK can also perform clustering analysis of the sequences based on the distance matrix.

**Fig. 1.**
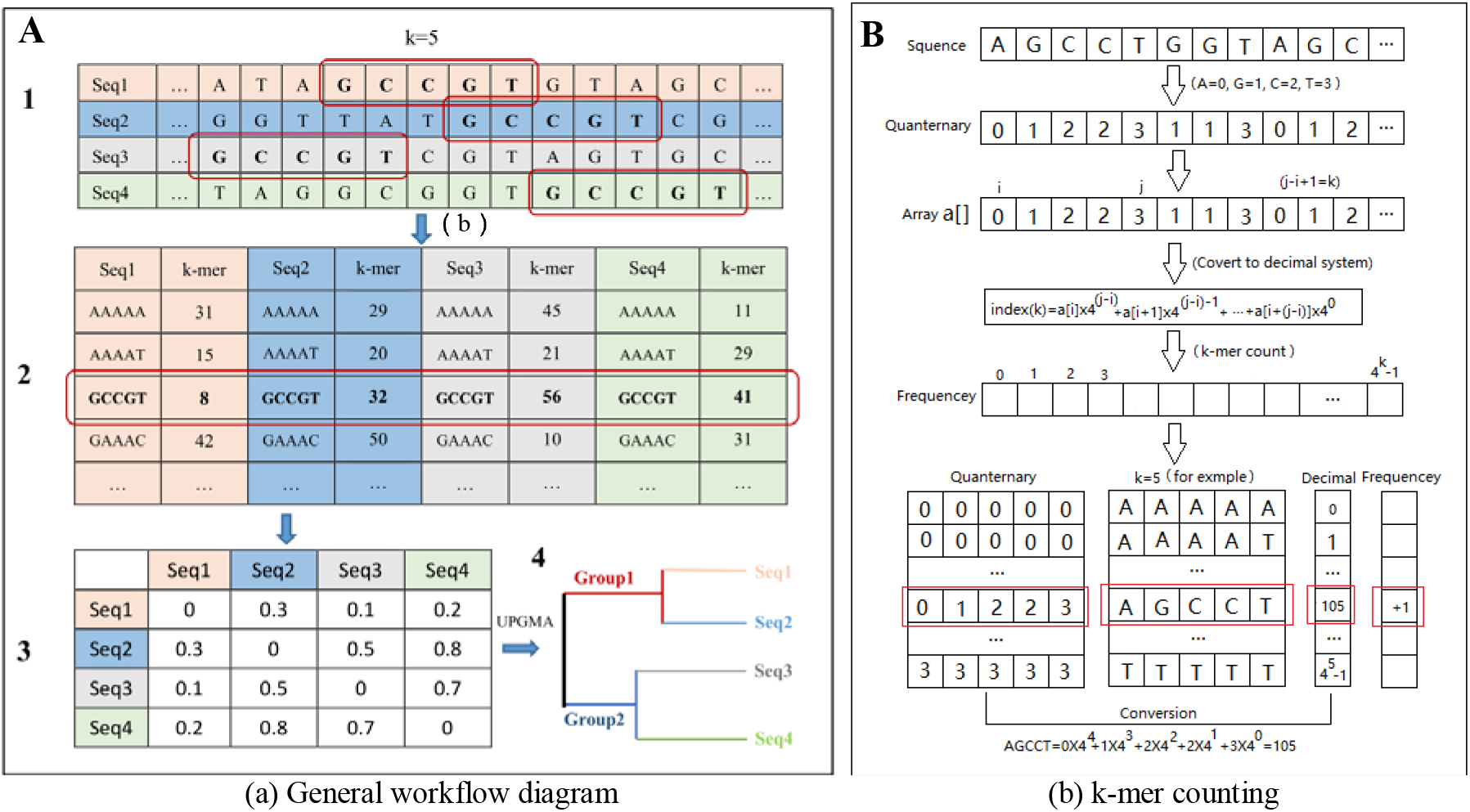
The Workflow of SeqDistK. In (**A**), we illustrated the steps of SeqDistK constructing a phylogenetic tree: (1) it counts k-mer occurrence (k=5 was shown, specifiable) in each input sequence (4 input sequences were shown); (2) it gathers k-mer occurrence vectors from all input sequences; (3) it computes the distance matrix of the input sequences using specifiable dissimilarity measures; (4) it draws the phylogenetic tree using the Unweighted Pair Group Method with Arithmetic Mean (UPGMA) algorithm via Phylip. In (**B**), we illustrated the algorithm and associated data structure for k-mer searching, counting and storage as implemented in SeqDistK. In principle, we mathematically transformed k-mer to an index that can randomly address and operate an array-based memory storage efficiently.

In the example illustrated in **Fig. 1A**, we analyzed four input sequences with 5-mer statistics and constructed their phylogenetic tree. When k=5, there are 4^5^ combinations of k-mer. Taking GCCGT as our example, in the first step, we counted the frequency of GCCGT. In the next step, the frequency vectors of GCCGT and all other 5-mers from each sequence were combined to form a matrix. In the third step, we computed the dissimilarity matrix by pairwise comparison of all k-mer vectors using the dissimilarity measures. And in the last step, we used UPGMA to draw the inferred phylogenetic tree using the open source software Phylip (http://evolution.genetics.washington.edu/phylip.html) **[22]**.

We implemented SeqDistK workflow into an open source software package, which features intuitive user interface, fast calculation, small memory and storage requirements. It accepts FASTA inputs and outputs a dissimilarity matrix file, which is compatible for Phylip to visualize the phylogenetic tree. SeqDistK allows real time specification of k-mer size (k) and, if needed, the order of Markov background (M). SeqDistK supports one-to-many and many-to-many comparisons. It incorporates most frequently used dissimilarity measures, such as Eu, Ma, Ch, Hao, d2, d2S and d2star. The time complexity of SeqDistK is O(*LN*^2^) and is independent of k, where N is the number of input sequences and L is their average length. The space complexity of SeqDistK is O(4^*k*^).

SeqDistK was implemented with several advanced programming technique: (1) It makes full use of multi-thread programming to improve CPU use through multi-core optimization. It is highly responsive even on personal computers. (2) It has a graphical interface that is simple, intuitive, and easy to interact with. (3) It is fully compatible with all current versions of a major operation system – MS Windows. We are migrating SeqDistK to Linux and MAC platforms.

### 2.2 k-mer based dissimilarity measures

Most alignment-free dissimilarity measures are statistical transformations of the k-mer frequency found in biological sequences. For all the dissimilarity measures, k is the key differentiating parameter. Previous results showed that, for k within the optimal range, many dissimilarity measures can perform well, even if there is rearrangement or missing bases within the input sequences. In most applications, the optimal k does not exceed 32. We implemented seven dissimilarity measures in SeqDistK, namely, Ch, Eu, Ma, d2, d2S, d2star **[20]** and Hao **[14-15]**. For d2S and d2star, their Markov orders were in the range M=0, 1, 2.

We introduce the seven dissimilarity measures implemented in SeqDistK as follows. For two sequences A’ = A _1_ A _2_ …A _n_ and B’ = B _1_ B _2_ …B _m_, with the length n and m respectively, where the letters of the sequences are drawn from a finite alphabet Λ∈ {A, C, G, T}, we define X_w_ and Y_w_, the occurrences of word w of length k in sequence A’ and B’ respectively, such that w∈Λ^k^. Let 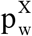 and 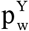 be the expected background probability of w in a model. For instance, a widely used measure d2 (Eq. 1) is simply the count of exact k-mer matches between two sequences, summing all k-mer at a given k **[25]**.

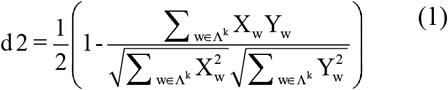

*Sun et al*. also defined dissimilarity measures d2S (Eq. 2) and d2star (Eq. 3), which is between 0 and 1 **[20]**:

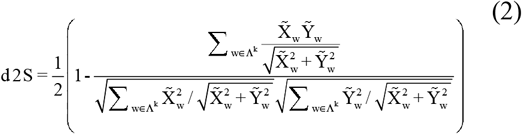

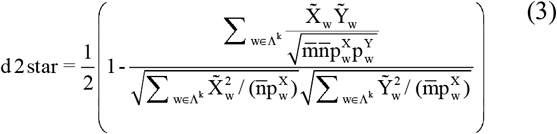

where, 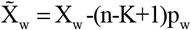 and 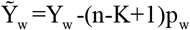 are the deviations of the observed occurrences from expected occurrences based on background models (Markov or i.i.d).

*Hao et al* **[14–15]** considered the relative difference vector of the number of occurrences of every k-mer w with its expected count under the (k-2)-th order of Markov model. They defined 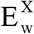 and 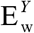 as the expected occurrences of *w*. Hao’s dissimilarity measure is:

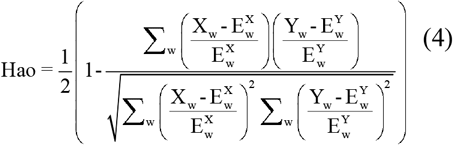

Dissimilarity measures such as d2S, d2star and Hao all can use Markov model to estimate background occurrence. The order of the Markov model is also a key parameter. There are also classical dissimilarity measures such as Manhattan (Eq. 5), Euclidean (Eq. 6), and Chebyshev (Eq. 7). d2, Eu, Ma and Ch measures do not consider a Markov background.

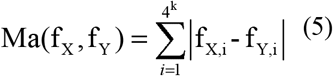

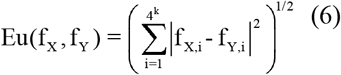

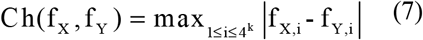

where

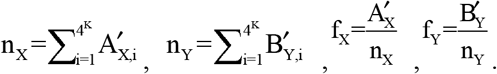

### 2.3 Benchmark dataset

While alignment-free phylogenetic analysis is unrestricted to specific genes, we used a golden standard 16S rRNA dataset for our benchmark for its wide acceptance. 16S rRNA is part of the 30S subgenre in the ribosome of prokaryotes, which is highly conserved in bacteria and archaea **[26]**. 16S rRNA gene is the most ancient gene among prokaryotes and contains both conserved and variable regions. Woese pioneered the idea of using 16S rRNA as biological phylogenetic indicators **[27]**. It is now a common practice to infer microbial taxonomy using multiple alignment or manual comparison of 16S rRNA sequences.

We downloaded a golden standard dataset of 16S rRNA sequences and the associated and expert curated phylogenetic tree from the *All-species Living Tree Project* (LTP) **[28-31]**. By the date, LTP has more than 6,700 entire 16S rRNA sequences, each of which represents a strain of bacteria. All sequenced strains of archaea and bacterial species were classified and preserved in LTP. We randomly selected 10 taxonomic groups, and from which we randomly selected 57 16S rRNA sequences. The ground truth tree is shown in **Fig. 2-a**, where branches are color-coded according to their taxonomy group (phylum, domains, classes, orders and genera). In the figure, Sequences C (blue), J (black), F (yellow), H (sky-blue) are Archaea and sequences A (light-gray), E (red), I (dark-purple), B (dark-blue), D (purple), and G (green) are Bacteria.

**Fig. 2.**
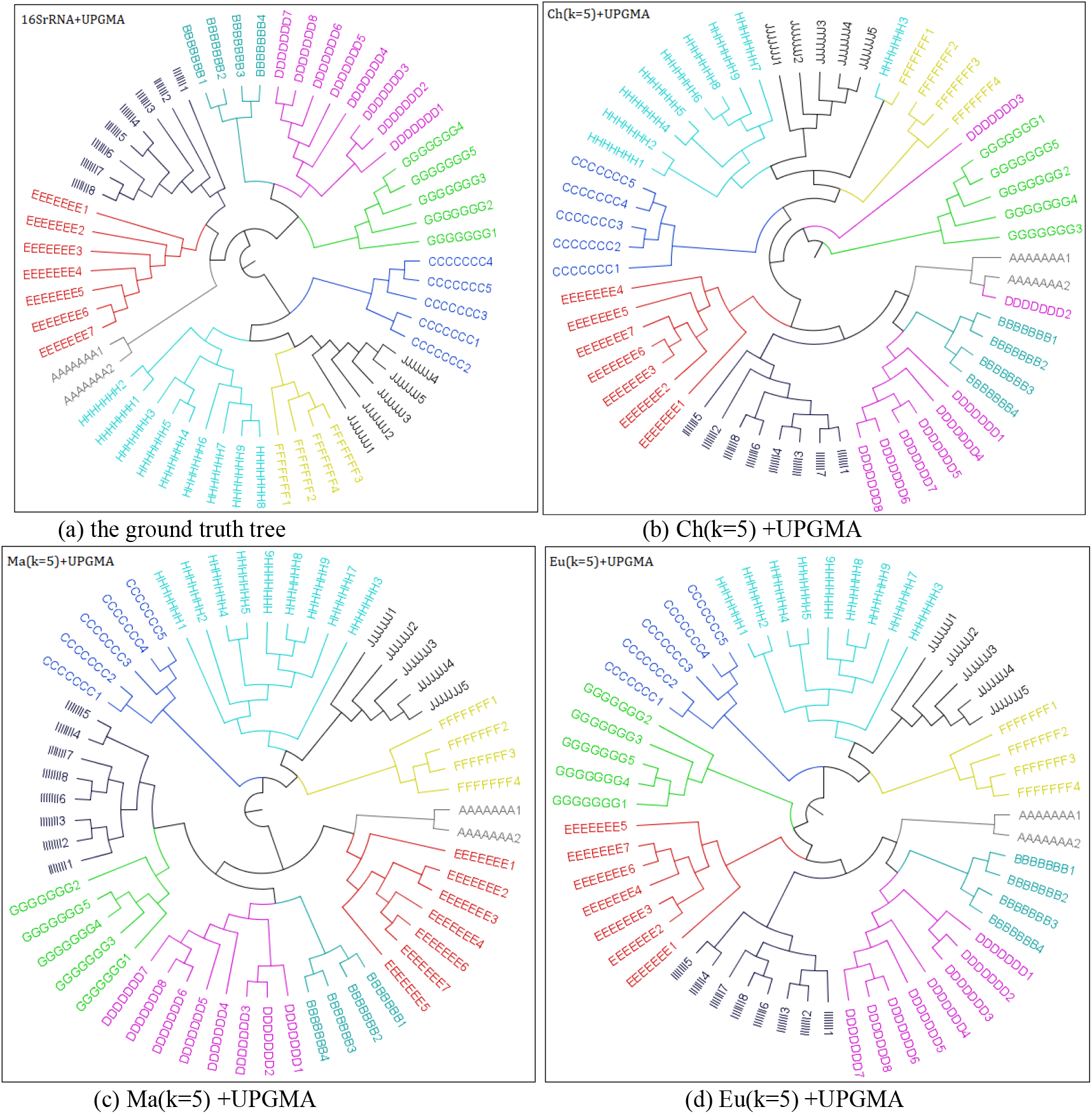
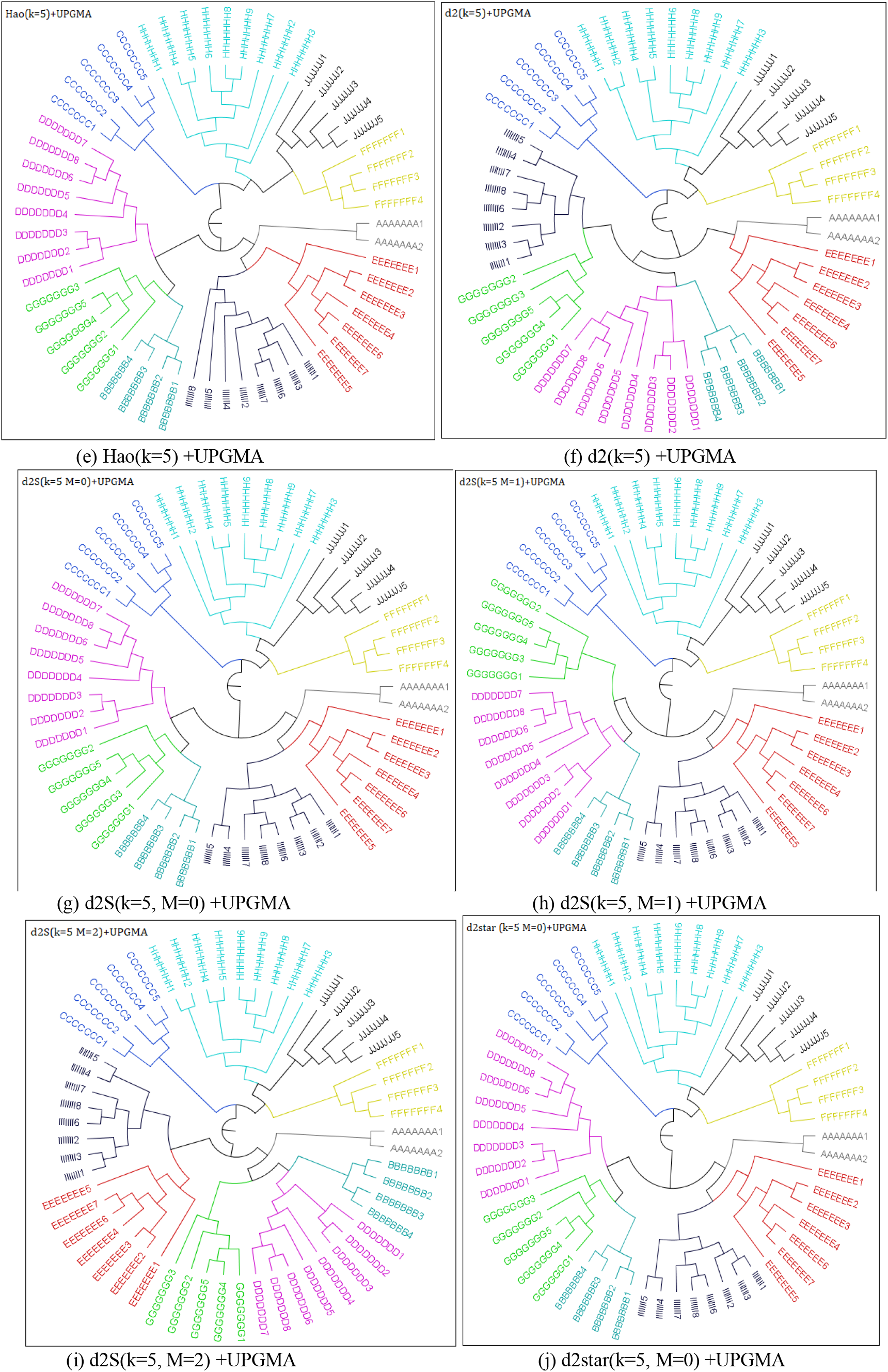
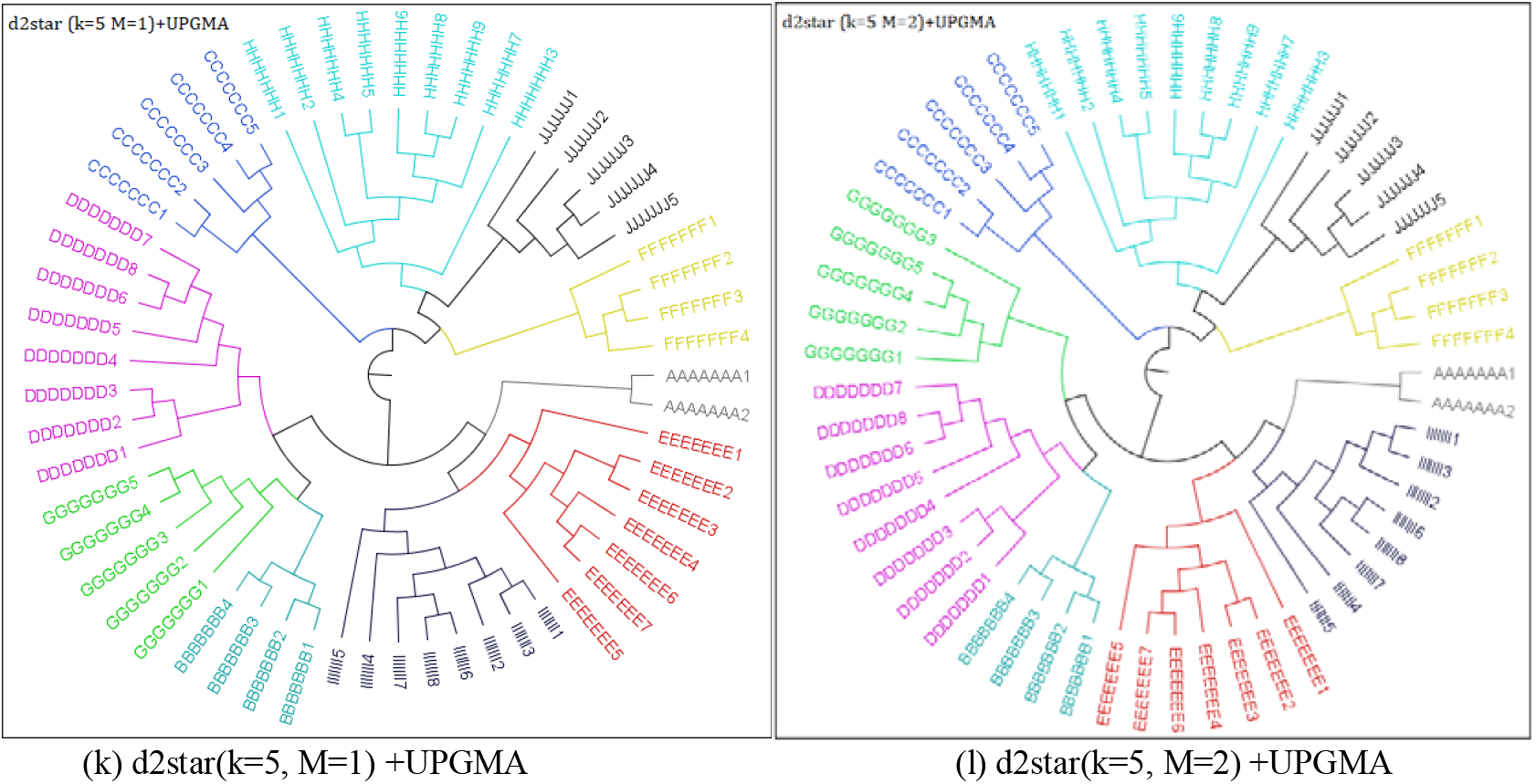
Phylogenetic trees from 16S rRNA sequences using 7 dissimilarity measures

### 2.4 Tree Distance

We used symmetric difference to compare phylogenetic trees. The symmetric difference is mathematically defined as the number of elements of two sets which are in either of the sets but not in their intersection. The symmetric difference of sets *A* and *B* was first proposed by Robinson et al as an evaluation standard to compare phylogenetic trees **[32]**. Compared with the parsimony score, another phylogenetic tree distance measure, symmetry difference takes into account the sequential order of hierarchical clustering and provides a systematic comparison. In addition, symmetry difference does not require branch length information, which is desirable because the ground truth tree is metric free. We computed the symmetric difference with Phylip’s TreeDist function.

## 3 Results

### 3.1 Phylogenetic trees benchmark

We applied SeqDistK to calculated the dissimilarity matrix of the 57 sequences selected from the LTP and inferred phylogenetic trees from these 16S rRNA sequences as illustrated in **Figs. 2** (b) to (k). As we can see, these inferred trees generally showed good accordance to the ground truth tree in **Fig. 2** (a). Indeed, for d2S and d2star, for Markov order ranging from 0 to 2, and k ranging from 4 to 10, all samples were correctly separated into their taxonomic group, achieving 100% classification accuracy. We thus chose k=5 for our down the line analysis. The phylogenetic trees obtained with k={4, 6, 7, 8, 9, 10} were also provided in the **Supplementary Figures**. As one can see from **Figs. 2** (b) to (k), compared with the ground truth tree (Fig.2 (a)), all other k-mer based dissimilarity measures, except for Ch, were also able to separate all the 57 sequences correctly into 10 their taxonomic groups.

### 3.2 Symmetric distance benchmark

With symmetric difference, we evaluated the phylogenetic tree inferred based on dissimilarity measures with a range of k and M. The symmetric difference between the inferred phylogenetic trees and the ground truth tree was shown in **Table 1**. All dissimilarity measures, except for Ch, had small symmetric difference at k=5. The symmetric difference by Ch was significantly larger, which means Ch inferred tree is different from the ground truth tree. This observation quantitatively substantiated our intuitive claim in the previous section. One potential reason for Ch’s low performance is, its metric is determined only using the maximum of all k-mer frequency difference, which is far from being a sufficient statistic data for all k-mers.

**Table 1.** Symmetric difference between inferred phylogenetic trees and the ground truth

| k | Ch | Eu | Ma | Hao | d2 | d2S<br>M=0 | d2S<br>M=1 | d2S<br>M=2 | d2star<br>M=0 | d2star<br>M=1 | d2star<br>M=2 |
| --- | --- | --- | --- | --- | --- | --- | --- | --- | --- | --- | --- |
| 4 | 17 | 17 | 18 | 17 | 17 | 17 | 17 | 13 | 17 | 18 | 14 |
| 5 | 22 | 14 | 16 | 17 | 14 | 16 | 15 | 13 | 16 | 17 | 17 |
| 6 | 22 | 16 | 16 | 15 | 16 | 16 | 16 | 16 | 17 | 17 | 17 |
| 7 | 21 | 16 | 16 | 18 | 16 | 16 | 16 | 16 | 17 | 17 | 17 |
| 8 | 25 | 16 | 16 | 16 | 16 | 16 | 16 | 16 | 17 | 16 | 17 |
| 9 | 28 | 16 | 16 | 16 | 16 | 16 | 16 | 16 | 17 | 16 | 17 |
| 10 | 86 | 16 | 16 | 16 | 16 | 16 | 16 | 16 | 16 | 16 | 17 |

SeqDistK also has better accuracy than Muscle, a popular multi-alignment based phylogenetic tool. For this data, the symmetry differences between SeqDistK inferred trees (except Ch) and ground truth is in the range of 13 to 18 (**Table 1**). In contrast, the symmetry difference between the Muscle multi-alignment derived phylogenetic tree using the same UPGMA procedure and the ground truth tree is significantly larger, at 62.

We then did an in-depth comparison of the 6 well-performing dissimilarity measures using radar charts (**Fig. 3**). All symmetric difference values were offset by 12 for better visualization. The offseted symmetric difference between the ground truth tree and inferred phylogenetic trees was plotted by k. Symmetric difference was then intuitively observed as the distance between the curve’s folding point and the center. The larger this distance gets, the larger the symmetric difference and vice versa.

**Fig. 3.**
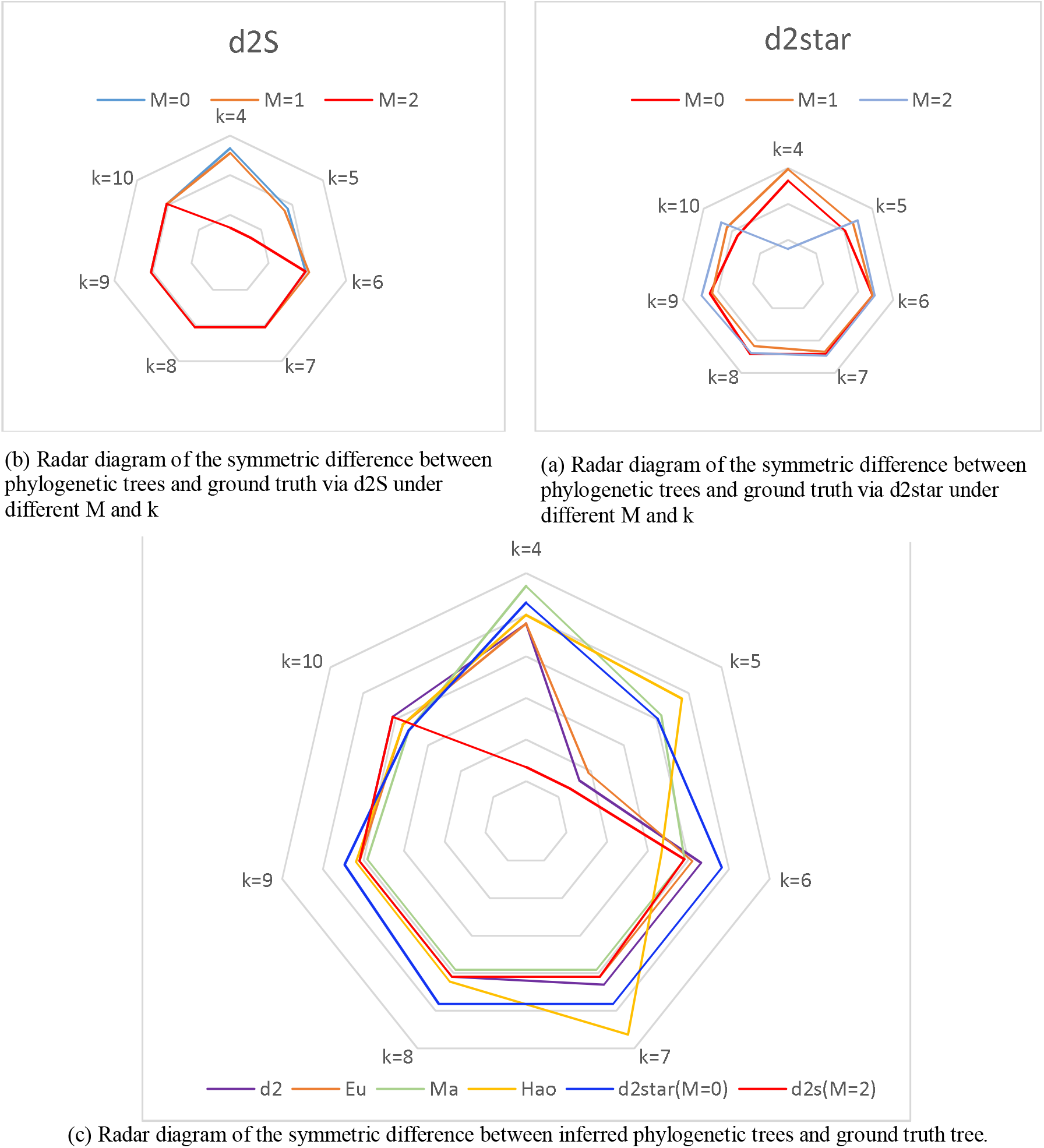
Radar diagram of the symmetric difference between phylogenetic trees and ground truth tree

As we can see from **Fig. 3** (a) and (b), d2S had its best performance at k=5, M=2. d2star had its best at k=4, M=2, while M=0 consistently gave d2star optimal results for other k values. We thus decided to use k=5, M=0 for d2star in downstream comparisons. It was also indicated by the figure that, if k=5 is selected, d2S was still sensitive to the Markov order parameter, while d2star is not. In **Fig.3** (c) we compared dissimilarity measures Eu, Ma, Hao, d2, d2star and d2S. We identified d2S as the overall best dissimilarity measure, though most other measures had good performance too (k=4 or k=5). Eu and d2 were the second tier best when k=5. In summary, we observed that the most k-mer measures with k>5 are consistently working well. Their performance becomes sensitive to k when k≤5. Summarizing from that, we have identified k=5 as the parameter for most effective alignment-free phylogenetics.

### 3.3 clustering application

We randomly selected and downloaded 63 sequences from the Silva database **[31]** for validating SeqDistK’s accuracy to phylogenetically classify these sequences. These 63 16S rRNA sequences were from 6 families. Based on our previous results, we set k=5 and computed the distance matrices for these sequences using d2, d2S (M=2), d2star (M=0) and Eu, respectively. We then input the dissimilarity matrices to the clustering analysis by multi-dimensional scaling (MDS). With MDS, we reduced all dissimilarity matrices into two-dimensional plots (**Fig. 4**). We used both shape and coloring to indicate families. The sequence set composition was presented in **Table 2** and further described in the **Supplementary Table 2**.

**Table 2.** Composition of the 16S rRNA sequence set

| Domain | Phylum | Class | Order | Family | Count | Marker |
| --- | --- | --- | --- | --- | --- | --- |
| Archaea | Euryarchaeota | Methanococci | Methanococcales | Methanocaldococcaceae | 12 | * |
|  | Crenarchaeota | Thermoprotei | Desulfurococcales | Desulfurococcaceae | 10 | * |
|  |  |  |  | Ignicoccaceae | 16 | * |
| Bacteria | Firmicutes | Clostridia | Thermoanaerobacterales | Thermodesulfobiaceae | 4 | ● |
|  | Thermotogae | Thermotogae | Kosmotogales | Kosmotogaceae | 10 | ● |
|  |  |  | Petrotogales | Petrotogaceae | 11 | ● |

**Fig. 4.**
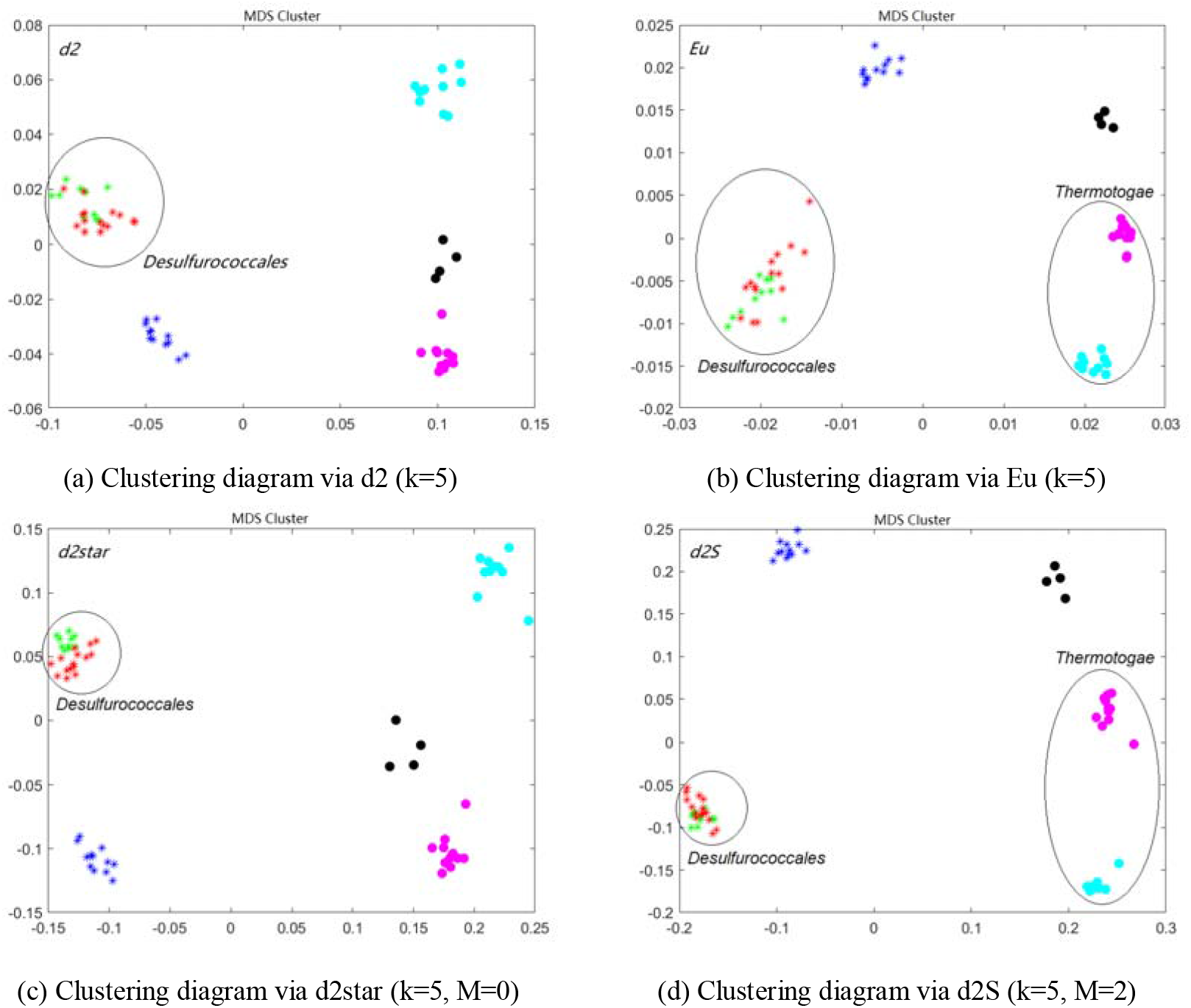
Clustering of 16S rRNA sequences

One expects an effective clustering algorithm would correctly group the sequences by their family identities. From **Fig. 4**, we did see that all the four dissimilarity measures correctly clustered the sequences. They all correctly separated bacteria families (right groups) from archaea families (left groups). Within archaea, the red and green colored sequences were closely clustered because those are all from the same order (*Desulfurococcales*), while the blue sequences were from a different order. Within bacteria, it can be also observed that cyan and purple colored sequences were more closely clustered because they are two orders of the same class (*Thermotogae*), while the black colored sequences were separated further because it belongs to a different phylum.

### 3.4 computation efficiency benchmark

**Table 2.** Runtime of SeqDistK by input sequence size

| 16s RNA<br>Sequences (N) | 50 | 200 | 500 | 1000 | 2500 | 5000 | 10000 |
| --- | --- | --- | --- | --- | --- | --- | --- |
| Time (Seconds) | 0.21 | 1.78 | 10.17 | 40.35 | 230.58 | 934.46 | 3854.99 |

To compare the computational efficiency between alignment-free and alignment based phylogenetics and between these k-mer statistics, we used SeqDistK and Muscle to compute the distance matrices of a series of randomly selected 16s RNA sequences. In Table 2, we first demonstrated the ability of SeqDistK to scale with large-scale data. As the number of sequences increased from 50 to 10,000, the runtime of SeqDistK increase linearly from 53.97s to 3854.99s. In Table 3, we demonstrated the superior scalability of SeqDistK as compared to Muscle (MEGA) [23-24]. As the number of sequences increased from 57 to 1824, the runtime of SeqDistK was capped at 175.76s while that of Muscle is 6714.58s, which means SeqDistK has a 3800% reduction of runtime. For all six measures chosen: Eu, Ma, d2, d2star, d2S, and Hao, the speed was comparable to each other and all much faster than that of Muscle.

**Table 3.**
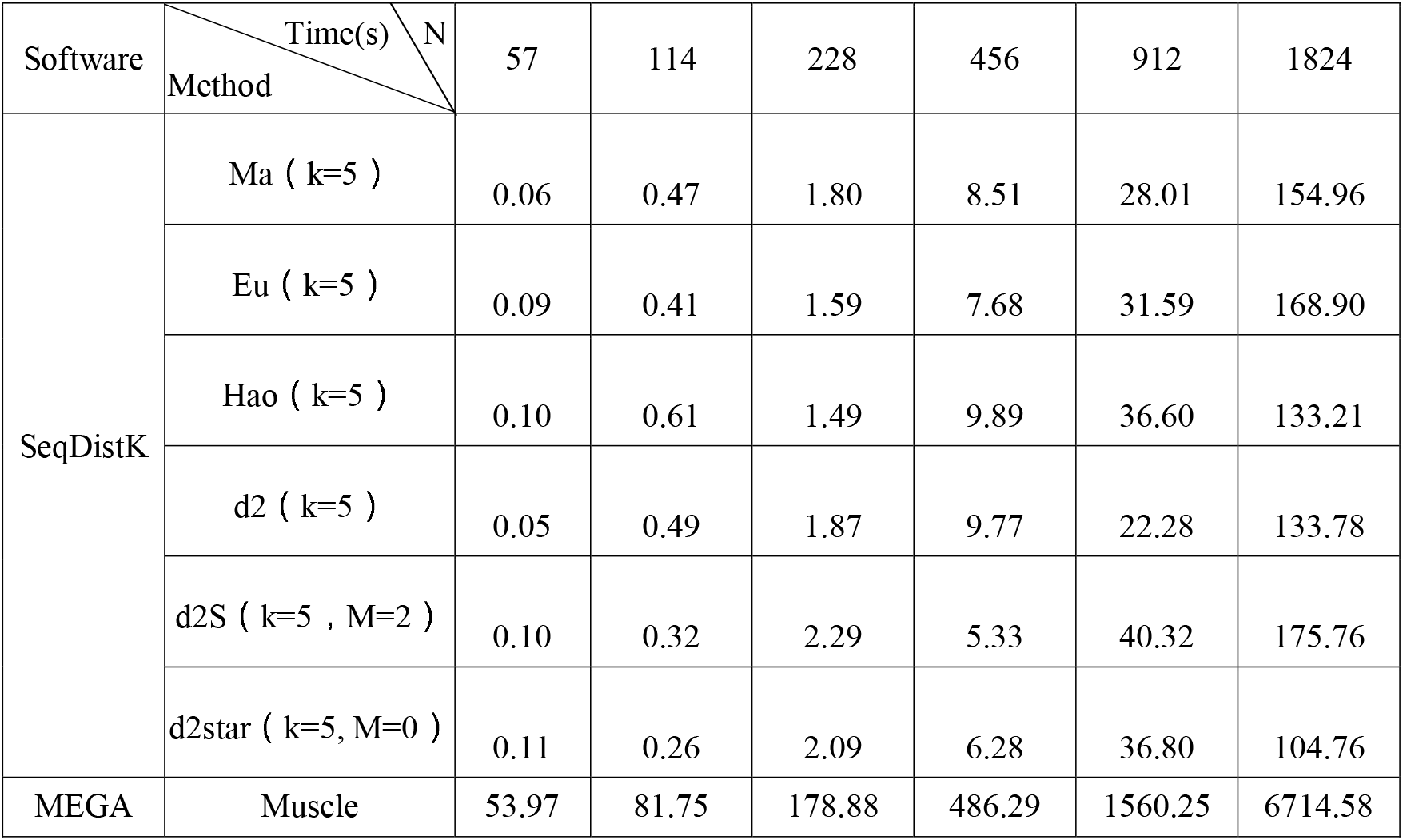
Efficiency comparison of alignment-based to and alignment-free algorithms

| Software | Time(s) \ N | 57 | 114 | 228 | 456 | 912 | 1824 |
| --- | --- | --- | --- | --- | --- | --- | --- |
|  | Method |  |  |  |  |  |  |
| SeqDistK | Ma ( k=5 ) | 0.06 | 0.47 | 1.80 | 8.51 | 28.01 | 154.96 |
|  | Eu ( k=5 ) | 0.09 | 0.41 | 1.59 | 7.68 | 31.59 | 168.90 |
|  | Hao ( k=5 ) | 0.10 | 0.61 | 1.49 | 9.89 | 36.60 | 133.21 |
|  | d2 ( k=5 ) | 0.05 | 0.49 | 1.87 | 9.77 | 22.28 | 133.78 |
|  | d2S ( k=5 , M=2 ) | 0.10 | 0.32 | 2.29 | 5.33 | 40.32 | 175.76 |
|  | d2star ( k=5, M=0 ) | 0.11 | 0.26 | 2.09 | 6.28 | 36.80 | 104.76 |
| MEGA | Muscle | 53.97 | 81.75 | 178.88 | 486.29 | 1560.25 | 6714.58 |

In addition, SeqDistK can set the CPU usage by itself, and in the software, if different sequences are used to place sequences that need to be calculated by different projects, different calculation projects can be run according to the folder, and multiple projects can be completed at one time, and the running speed is fast. The interface is simple and the directory settings are flexible. All the runtime comparisons were computed with (k=5) and using a personal computer, with Intel Core i7-4790K CPU @ 4.00GHz, 32G memory, Windows10.

## 4 Discussion

Phylogenetic tree is an important tool for comparative analysis of genomic data in the context of evolution. Based on molecular phylogenetic, this field has elucidated the evolution of genes and proteins since the early 1960s. With the advent and spread of next generation sequencing, very large datasets are available for phylogenetic inference. Phylogenetics at this scale require large data storage, high computing power, and a large amount of memory. Phylogenetics also need to adapt to different sequence patterns (e.g., protein coding and non-coding regions, sequence region may have different origins and evolution of history, etc). Therefore, it is critical to develop new phylogenetic method with computational scalability.

Alignment-free dissimilarity measures provide a good search space for efficient phylogenetics. We can easily extract, index, store and retrieve k-mers from biological sequences. We can capture homologous signals by summarizing on k-mer counts or frequencies and transform them into distance matrix. Unlike multi-sequence alignment, k-mer distance trees can be calculated efficiently agnostic of sequence regions. We showed in this paper, the dissimilarity measures are reliable basis for large-scale phylogenetics. Our software package SeqDistK demonstrated fast calculation and high precision, which is suitable for phylogenetic research of complex large-scale metagenomics datasets.

## Acknowledgements

L.C.X. wants to thank Dr. Nancy Zhang at University of Pennsylvania and Dr. Hanlee Ji at Stanford University for their support and helpful discussions. L.C.X. is supported by the United States National Institute of Health (HG006137-07), the American Cancer Society (132922-PF-18-184-01-TBG), the Innovation in Cancer Informatics fund, and funds from Intermountain Healthcare.

## Appendix A. Supplementary materials

### (1) Working interface

**Fig. 1.**
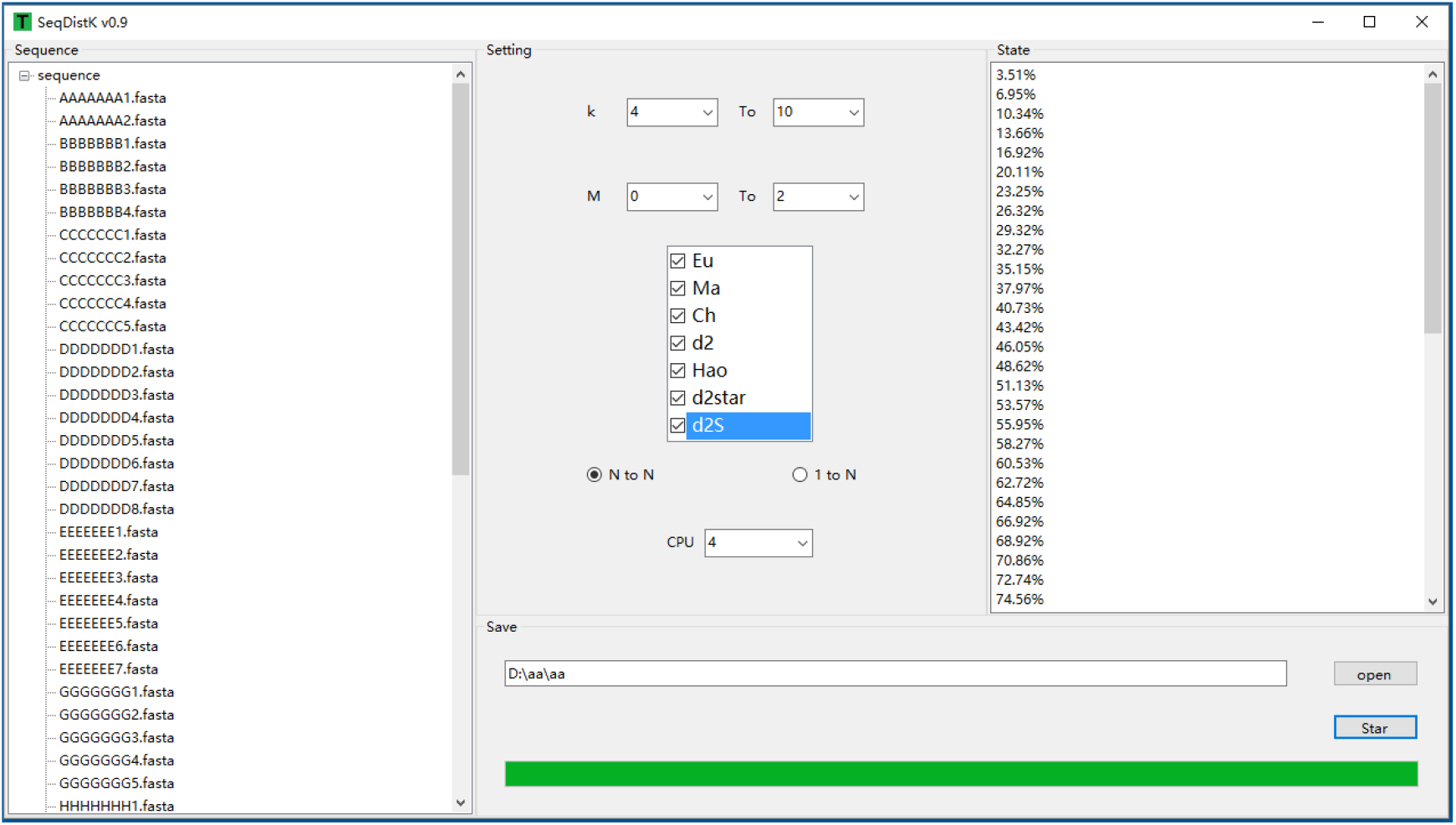
The working interface of SeqDistK

SeqDistK is a tool to calculate the distance among sequences and the window interface as Fig.1. There are 7 dissimilarity measures Eu, Ma, Ch, d2, d2star, d2S and Hao in SeqDistK. The software supports Windows. The advantage of the software is convenience. The whole process can be operated by mouse. It supports multiple directory-unit and multilevel directory structure and each directory will calculate independently. The results will be saved in directory structures independently.

### (2) Supplementary Tables

**Table 1.**
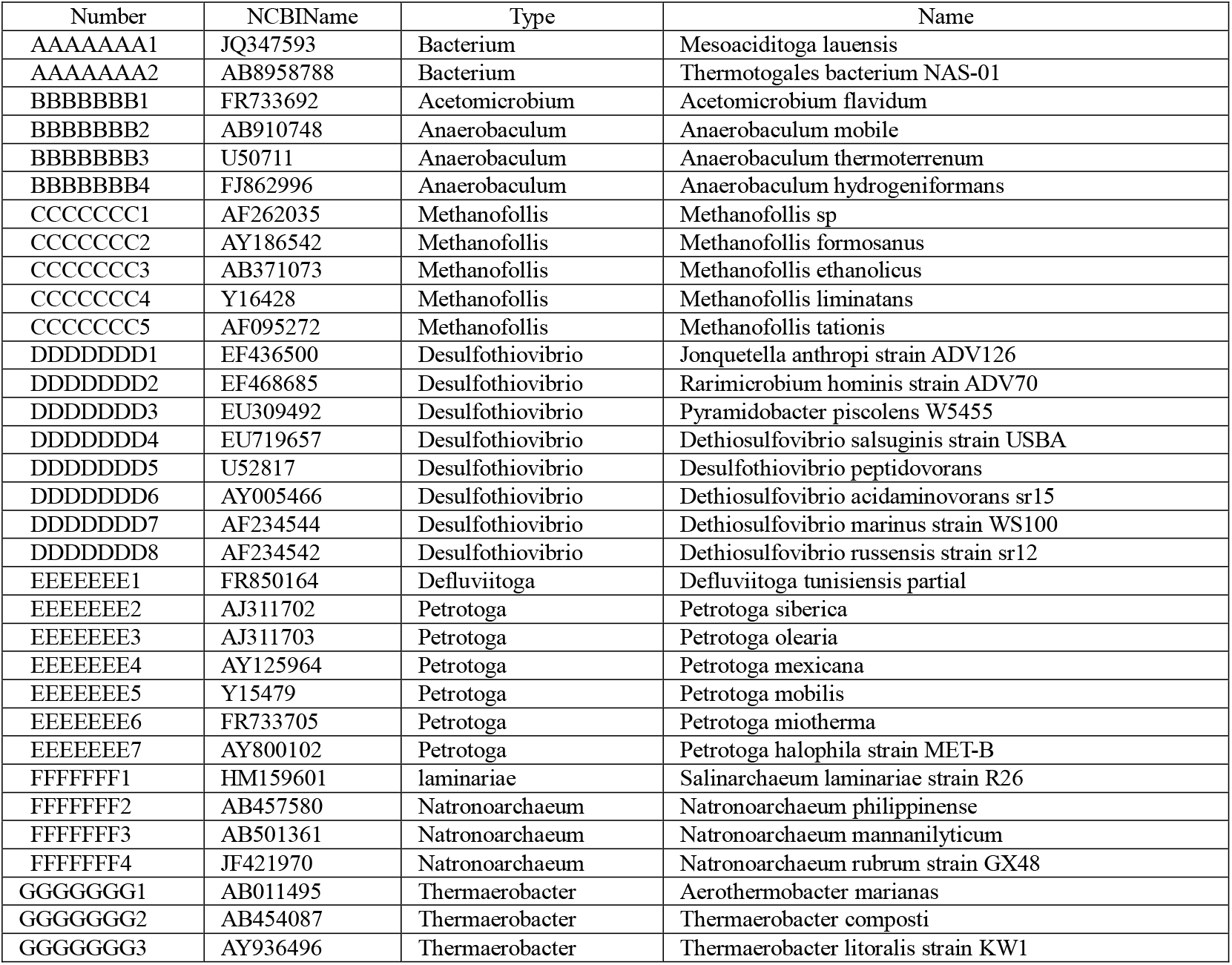
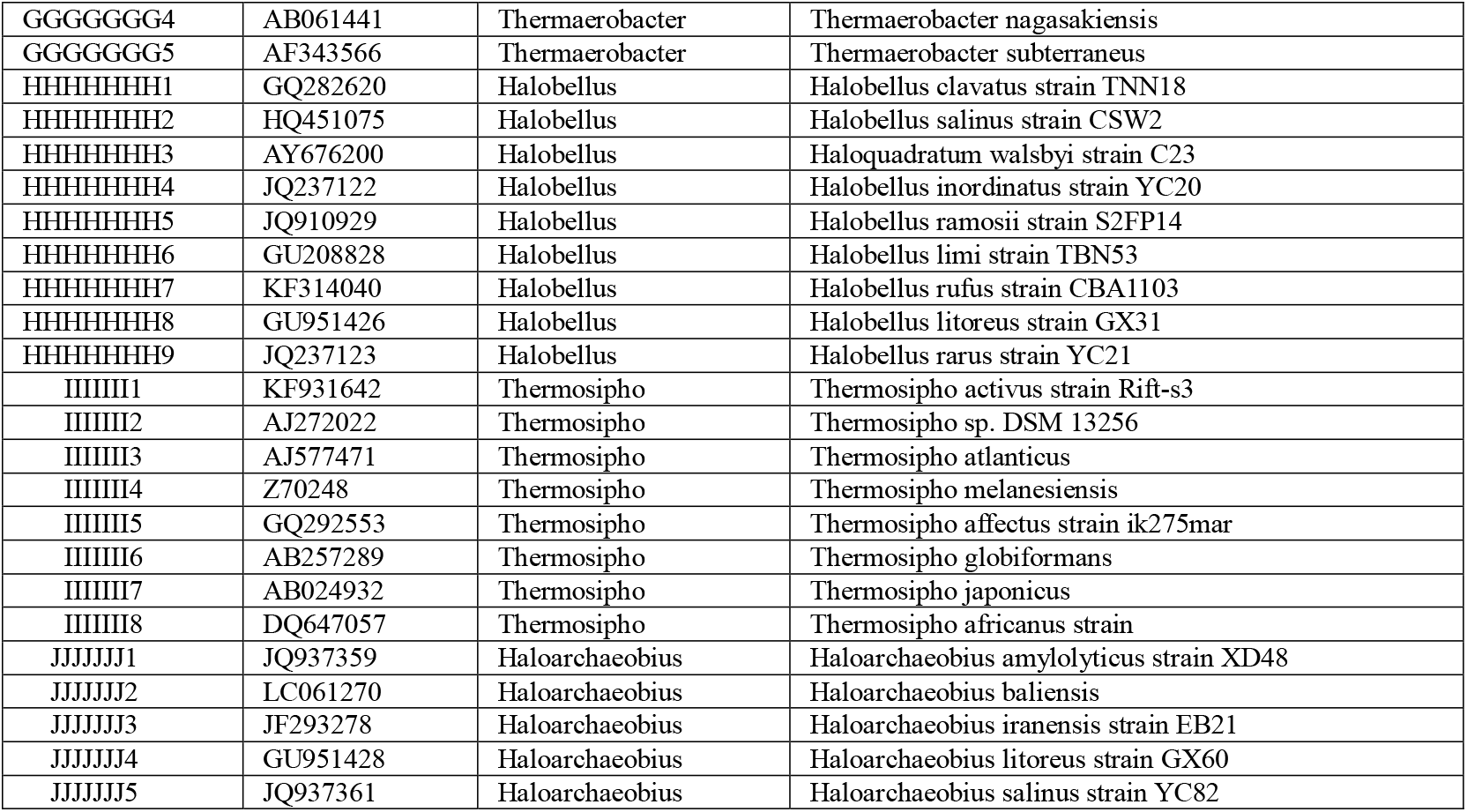
Data sets of 16S rRNA sequences(57 sequences)for standard tree

**Table 2.**
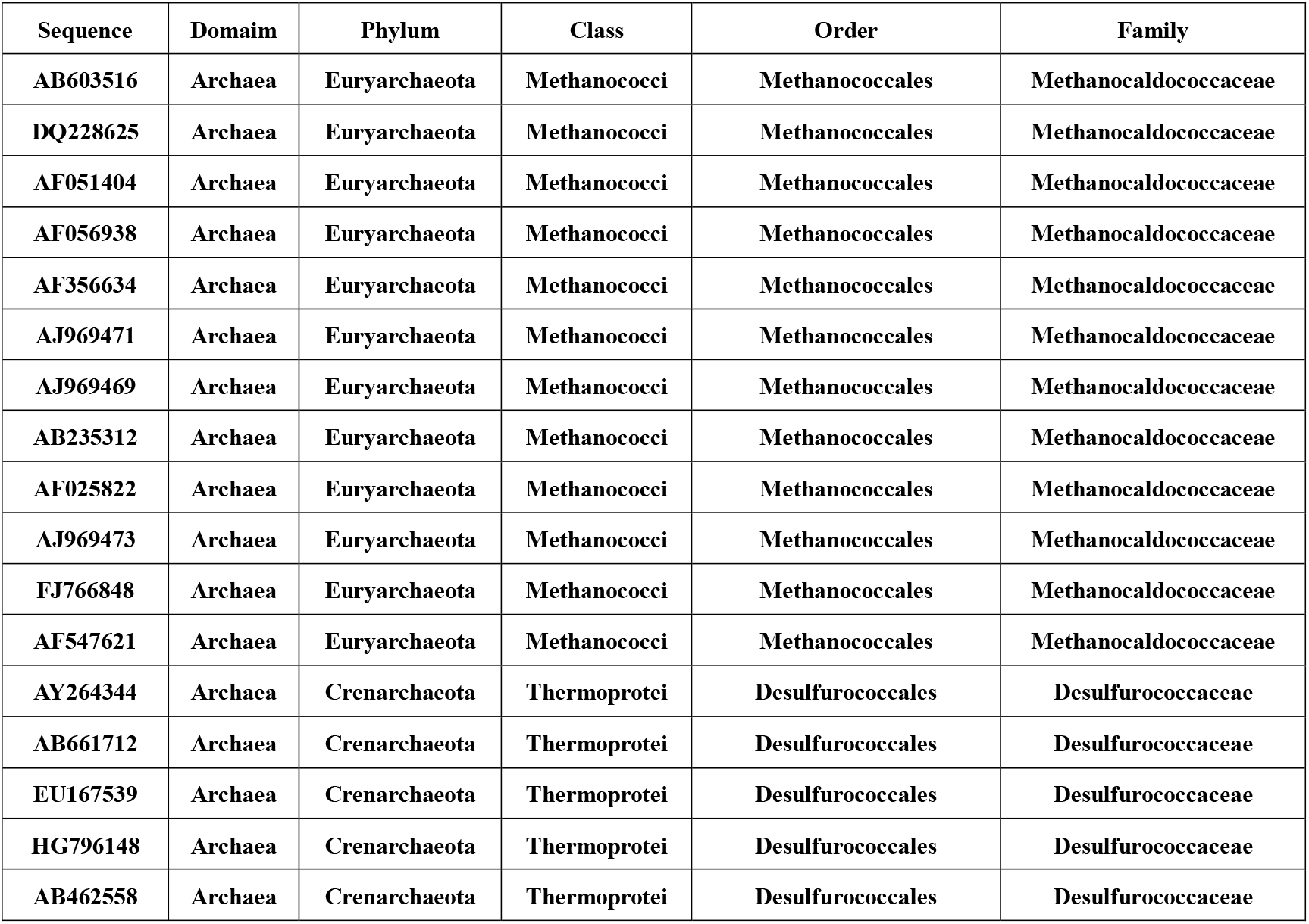
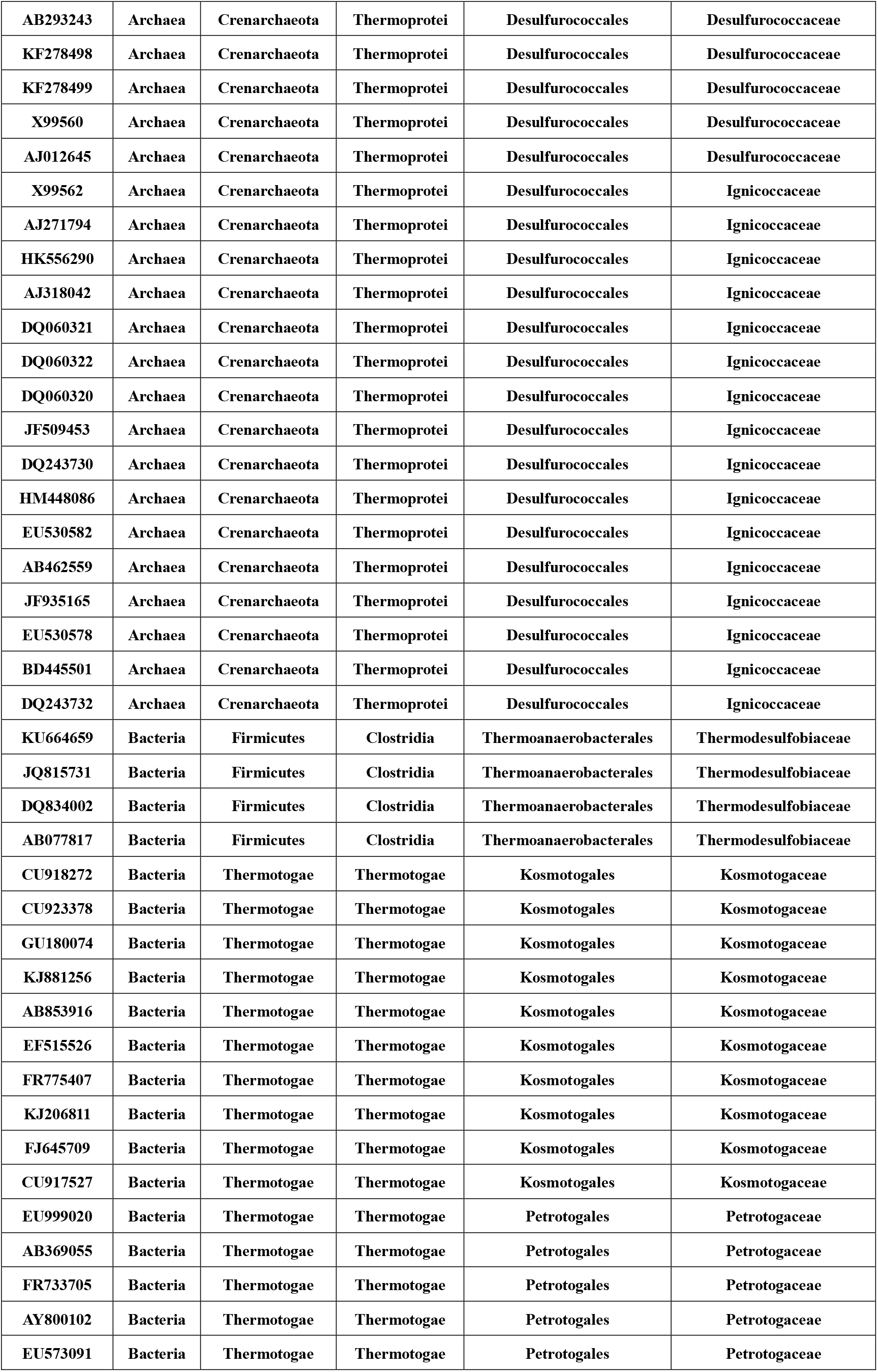
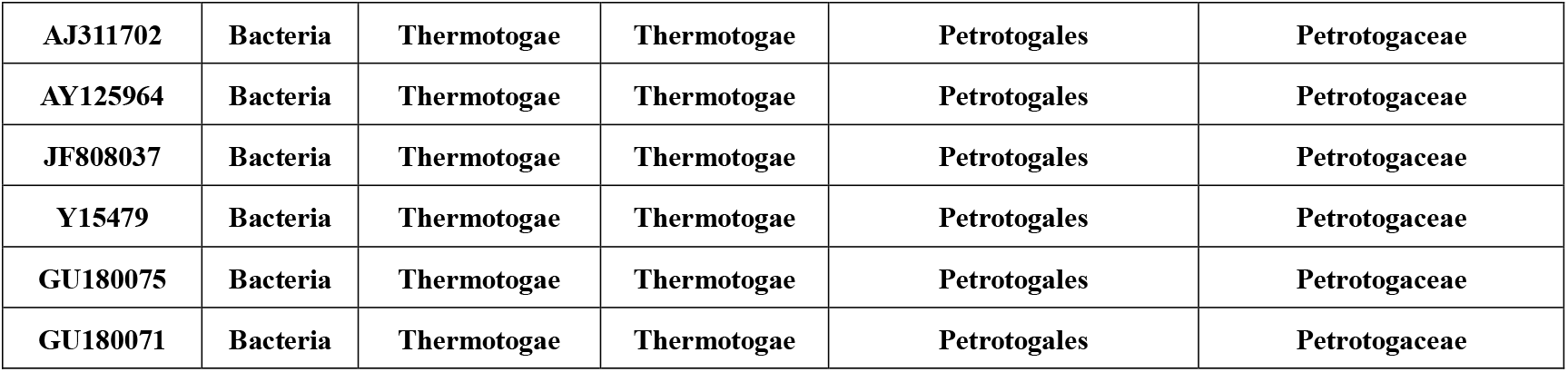
Data sets of 16S rRNA sequences(63sequences)for clustering

### (3) Supplementary Figures

**Fig.2.**
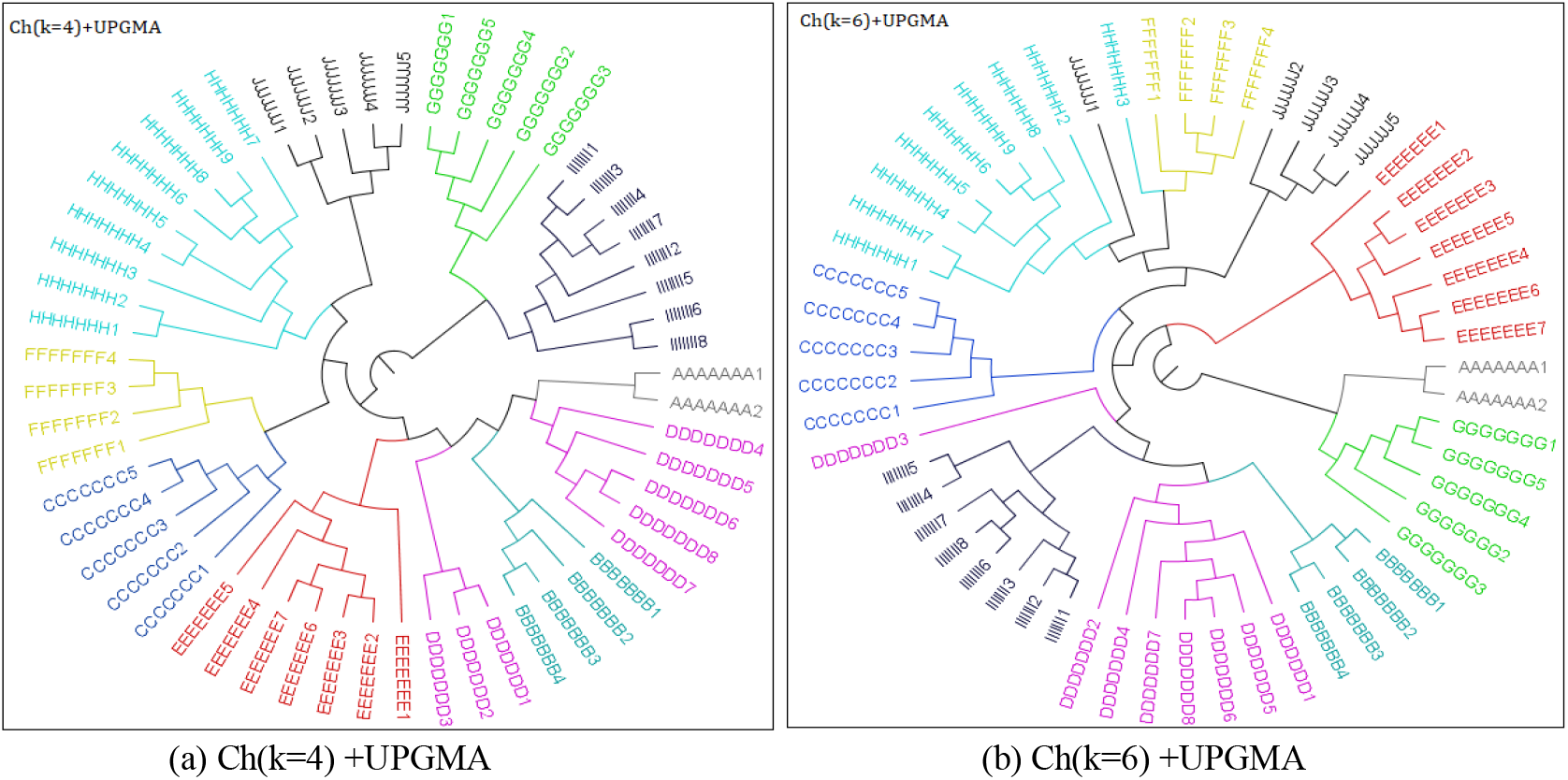
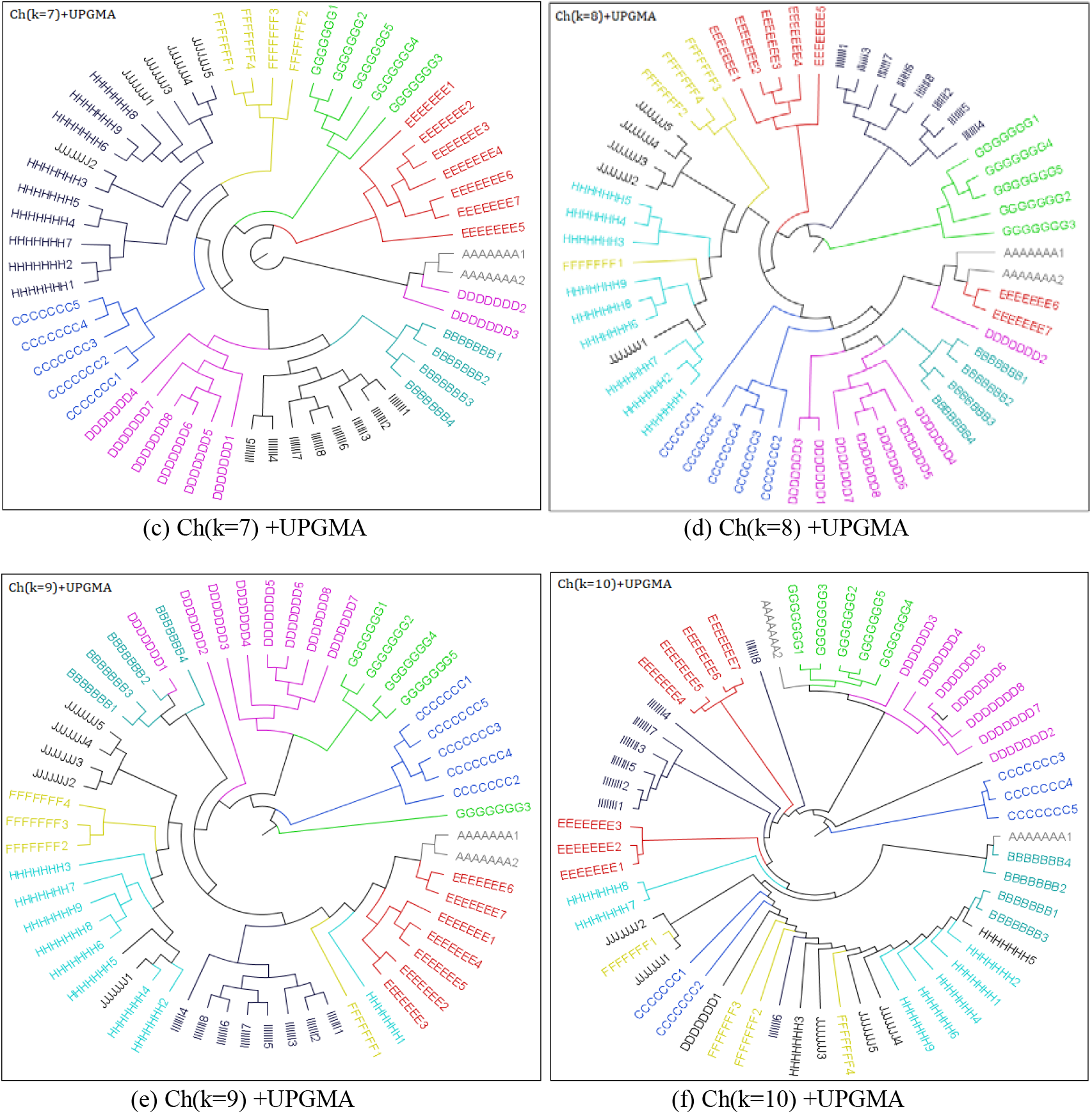
Phylogenetic trees for 16S rRNA sequences(57 sequences)via Ch (k=4, 6, 7, 8, 9, 10)

**Fig.3.**
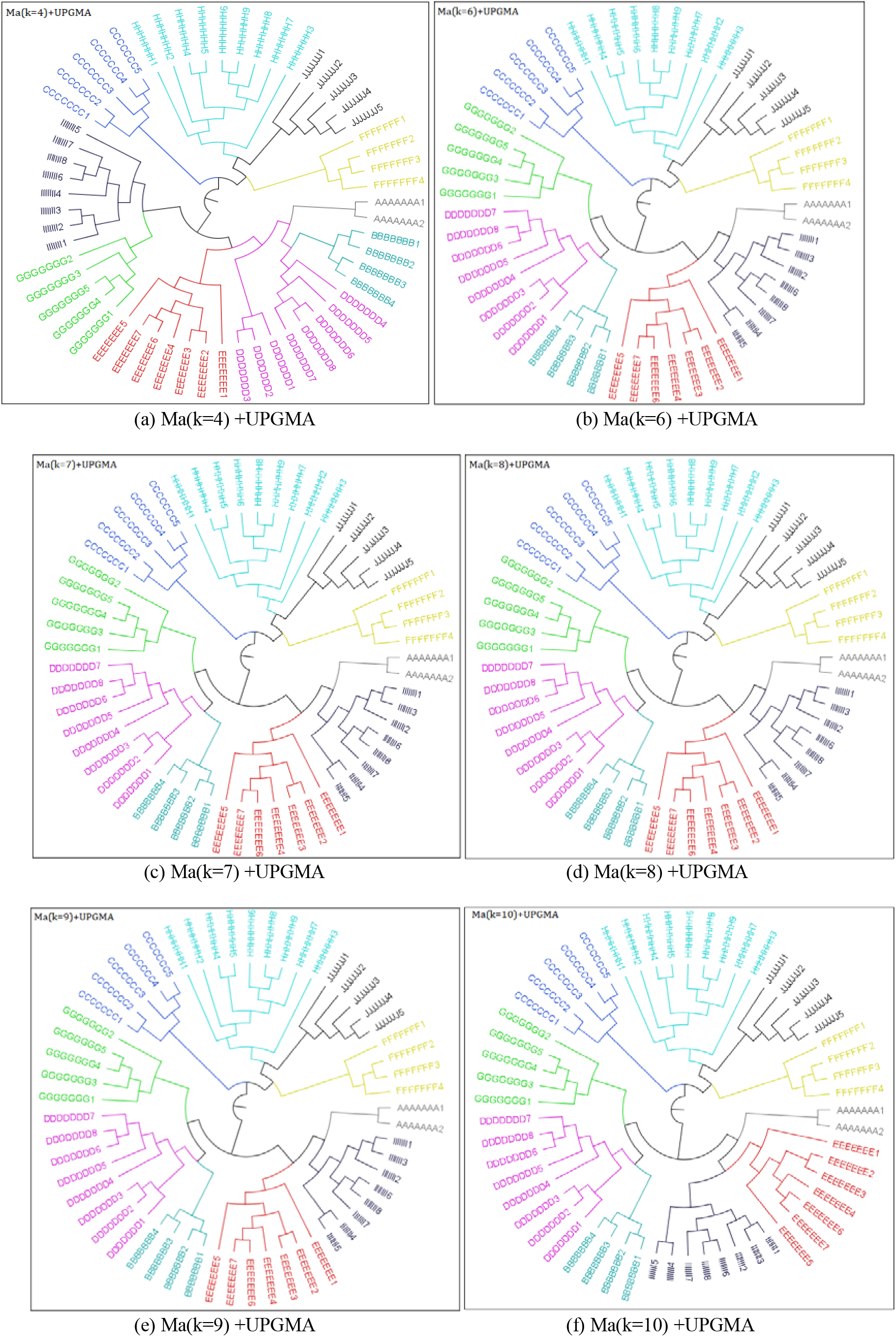
Phylogenetic trees for 16S rRNA sequences(57 sequences)via Ma (k=4, 6, 7, 8, 9, 10)

**Fig.4.**
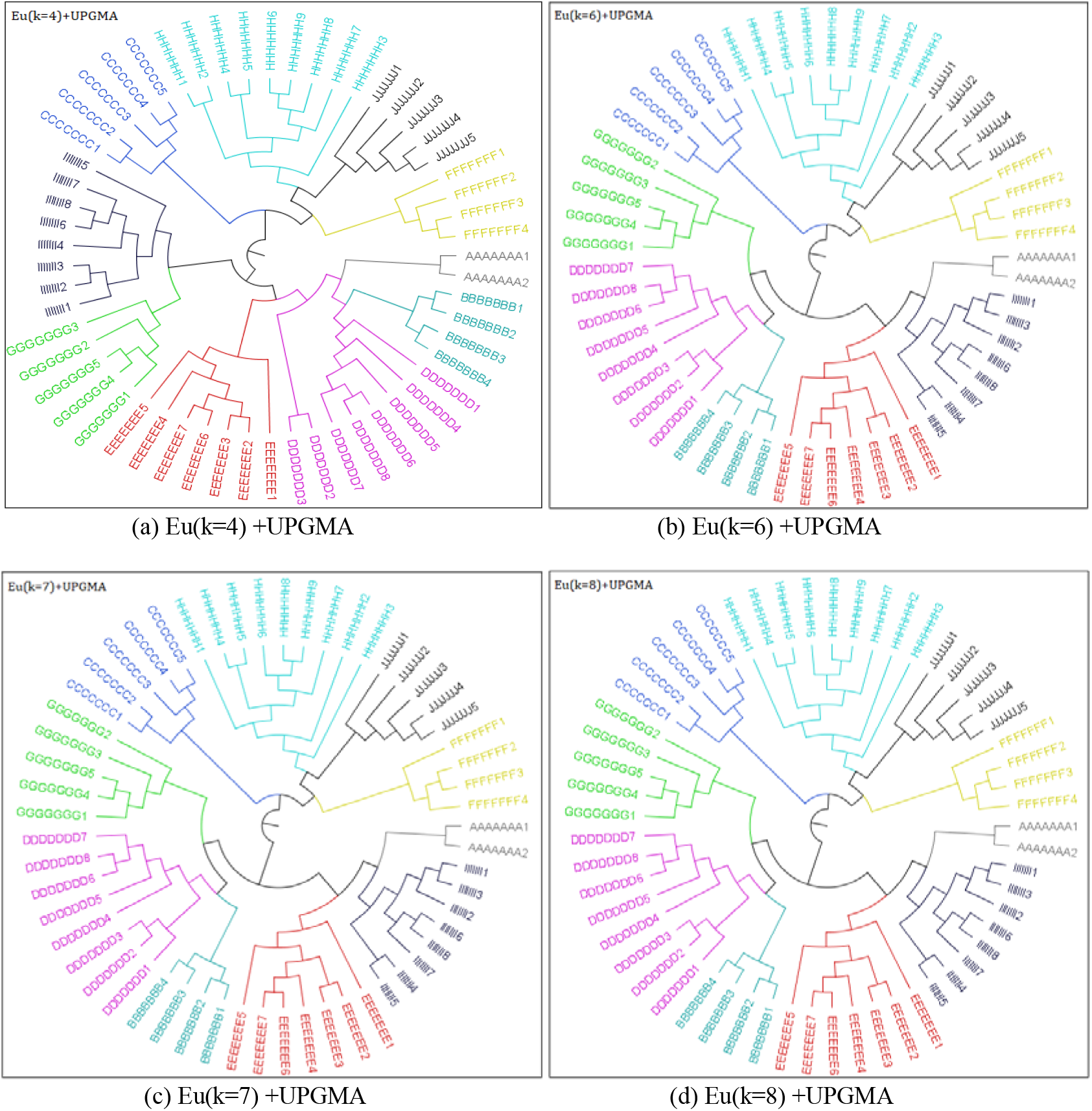
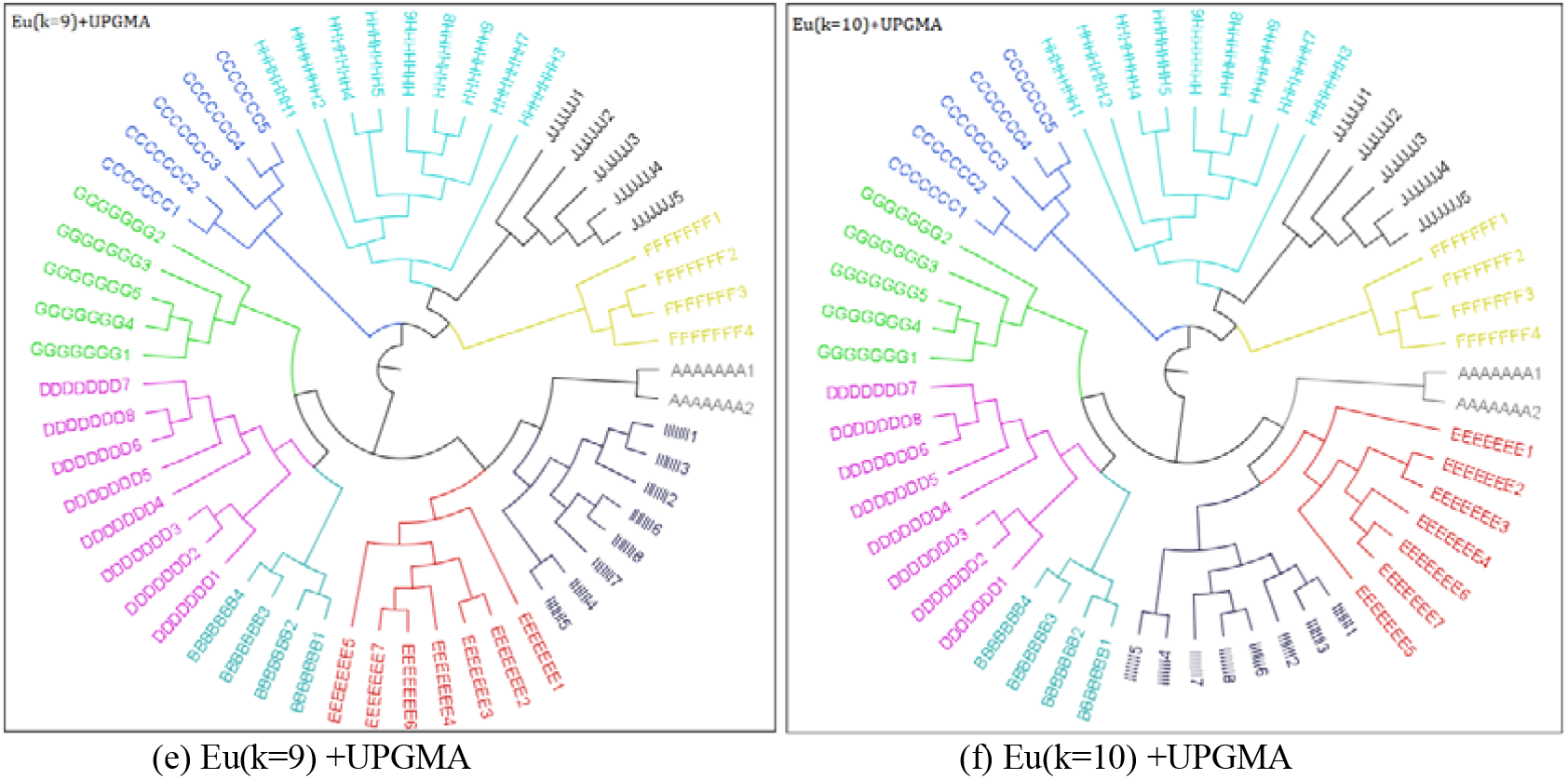
Phylogenetic trees for 16S rRNA sequences(57 sequences)via Eu (k=4, 6, 7, 8, 9, 10)

**Fig.5.**
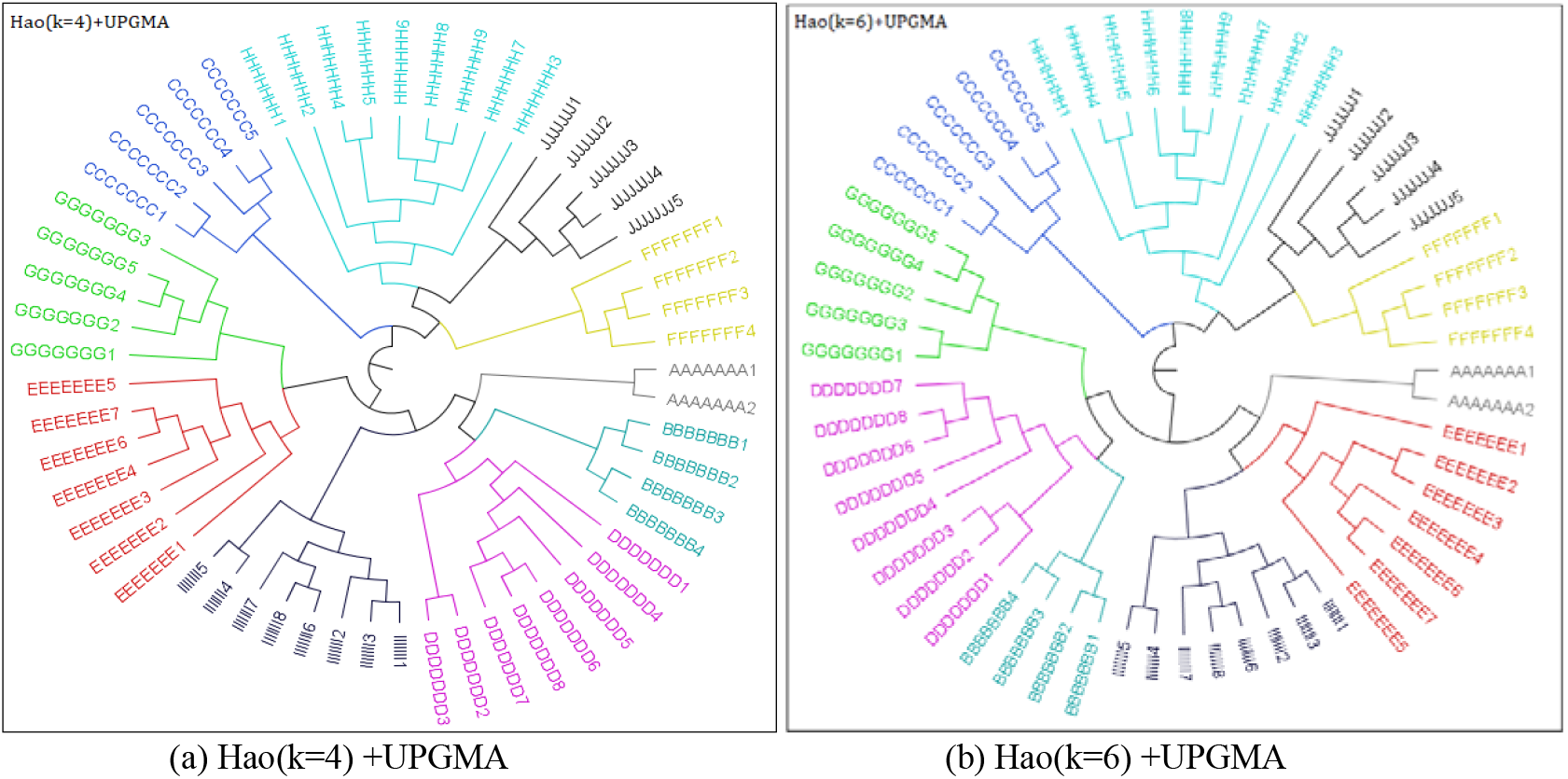
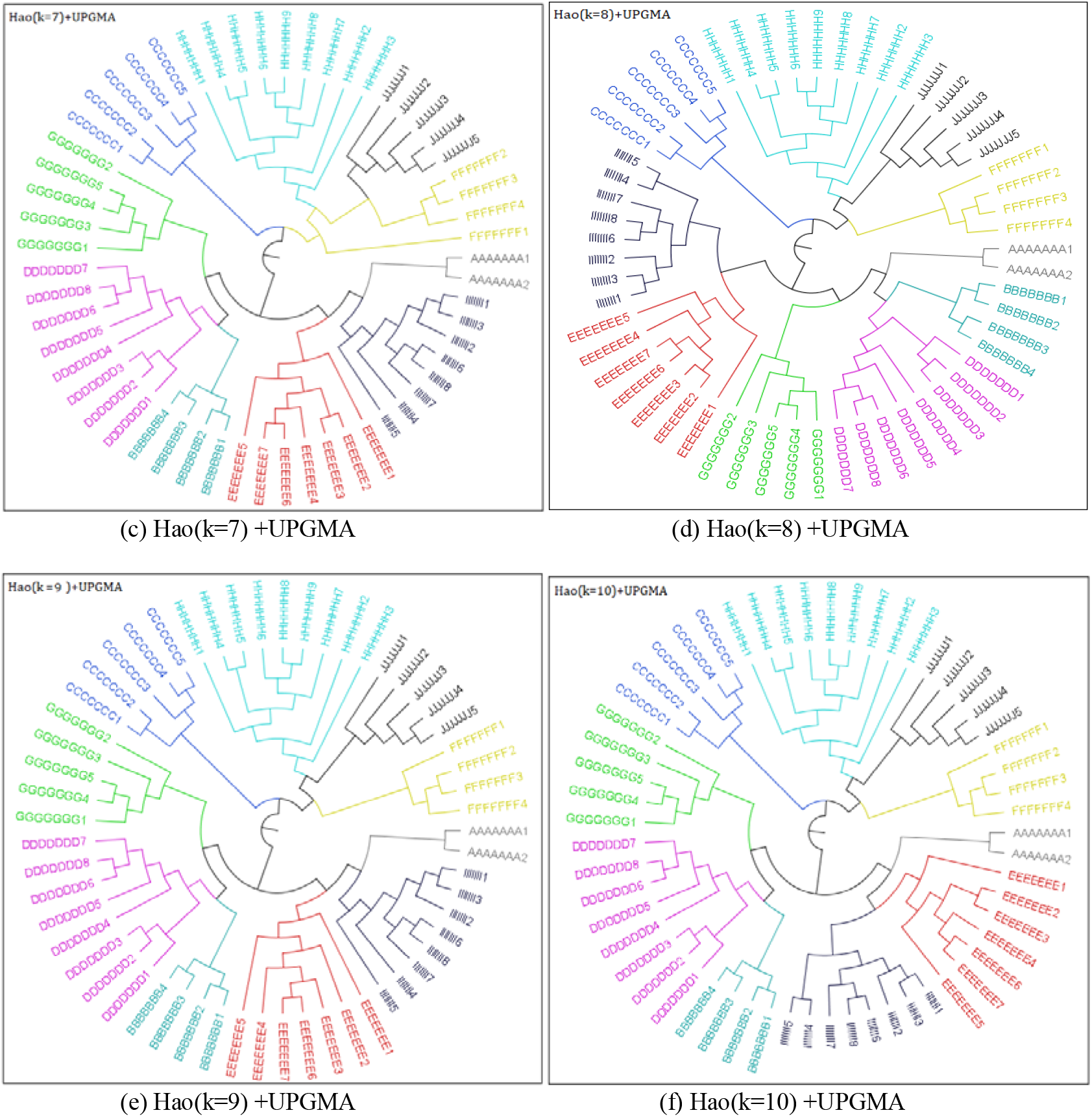
Phylogenetic trees for 16S rRNA sequences(57 sequences)via Hao (k=4, 6, 7, 8, 9, 10)

**Fig.6.**
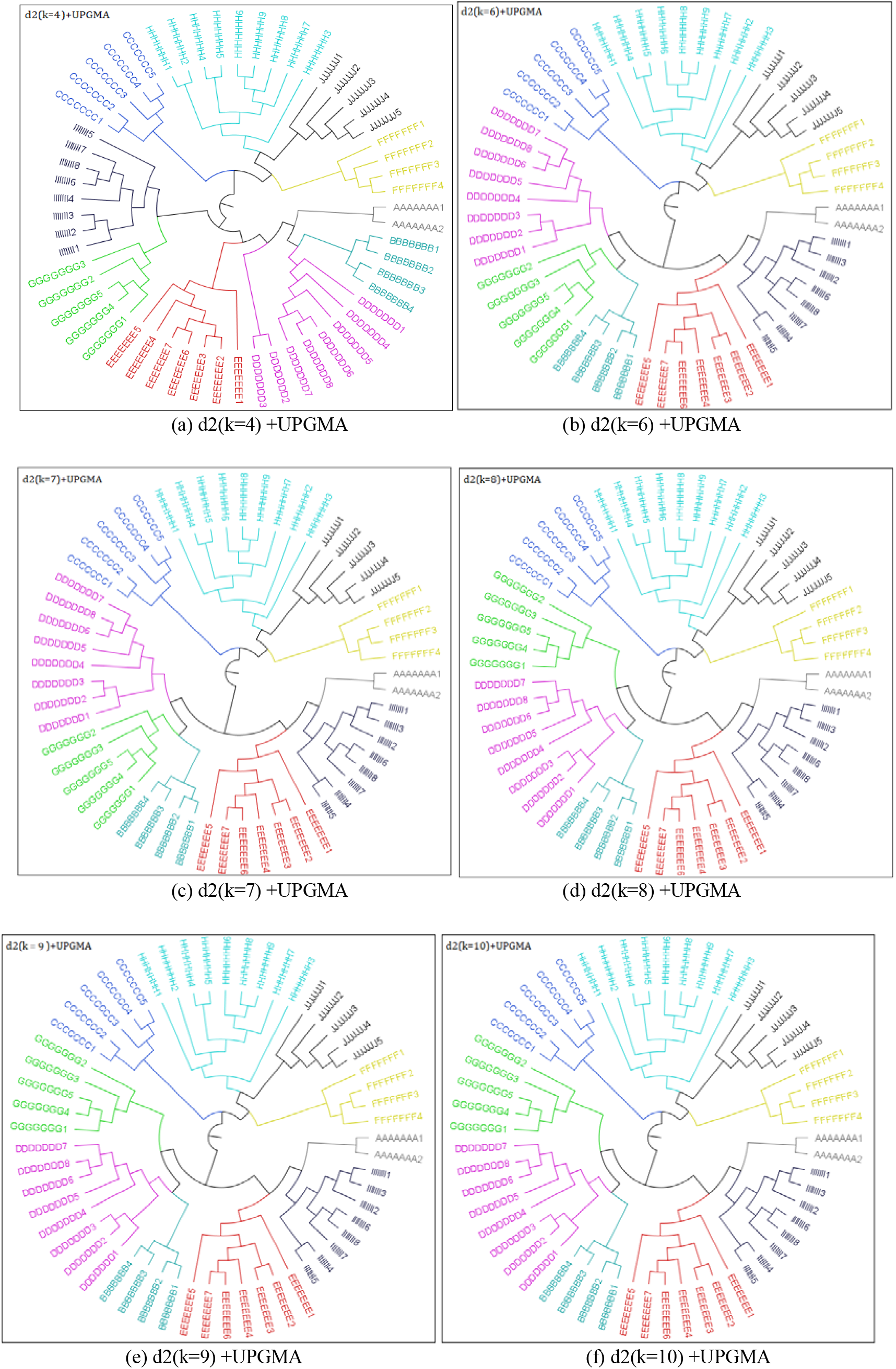
Phylogenetic trees for 16S rRNA sequences(57 sequences)via d2 (k=4, 6, 7, 8, 9, 10)

**Fig.7.**
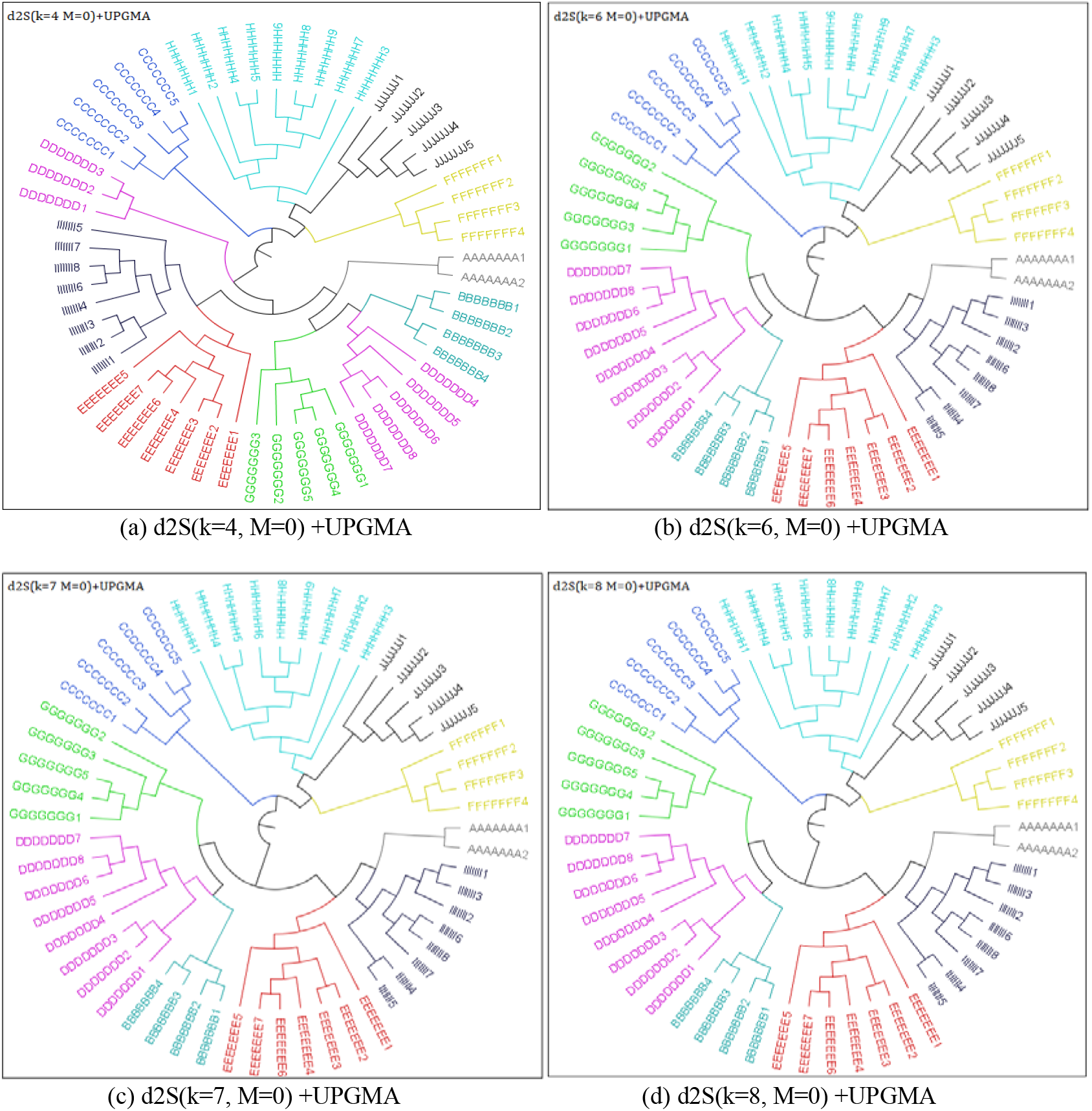
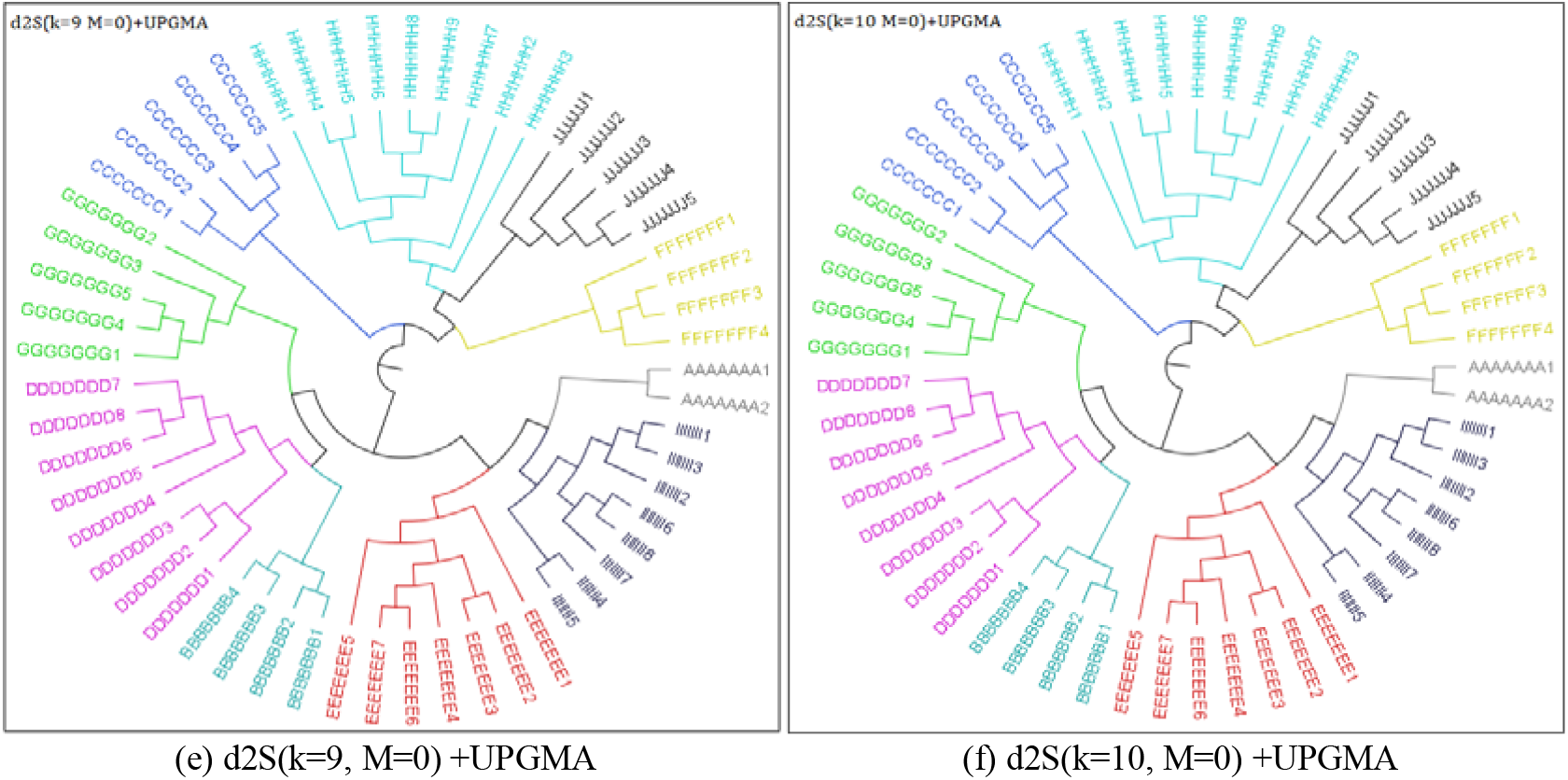
Phylogenetic trees for 16S rRNA sequences(57 sequences)via d2S (k=4, 6, 7, 8, 9, 10, M=0)

**Fig.8.**
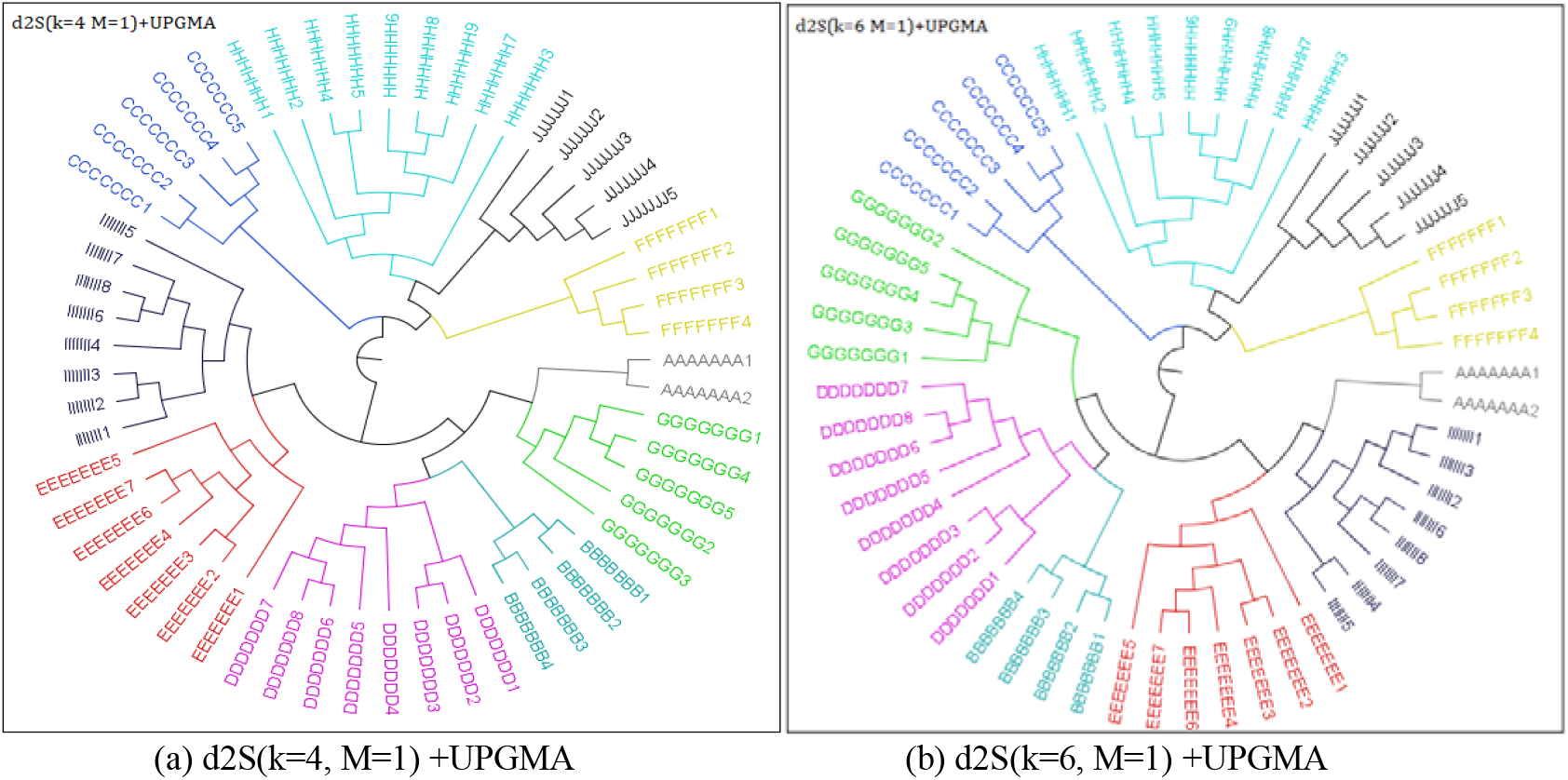
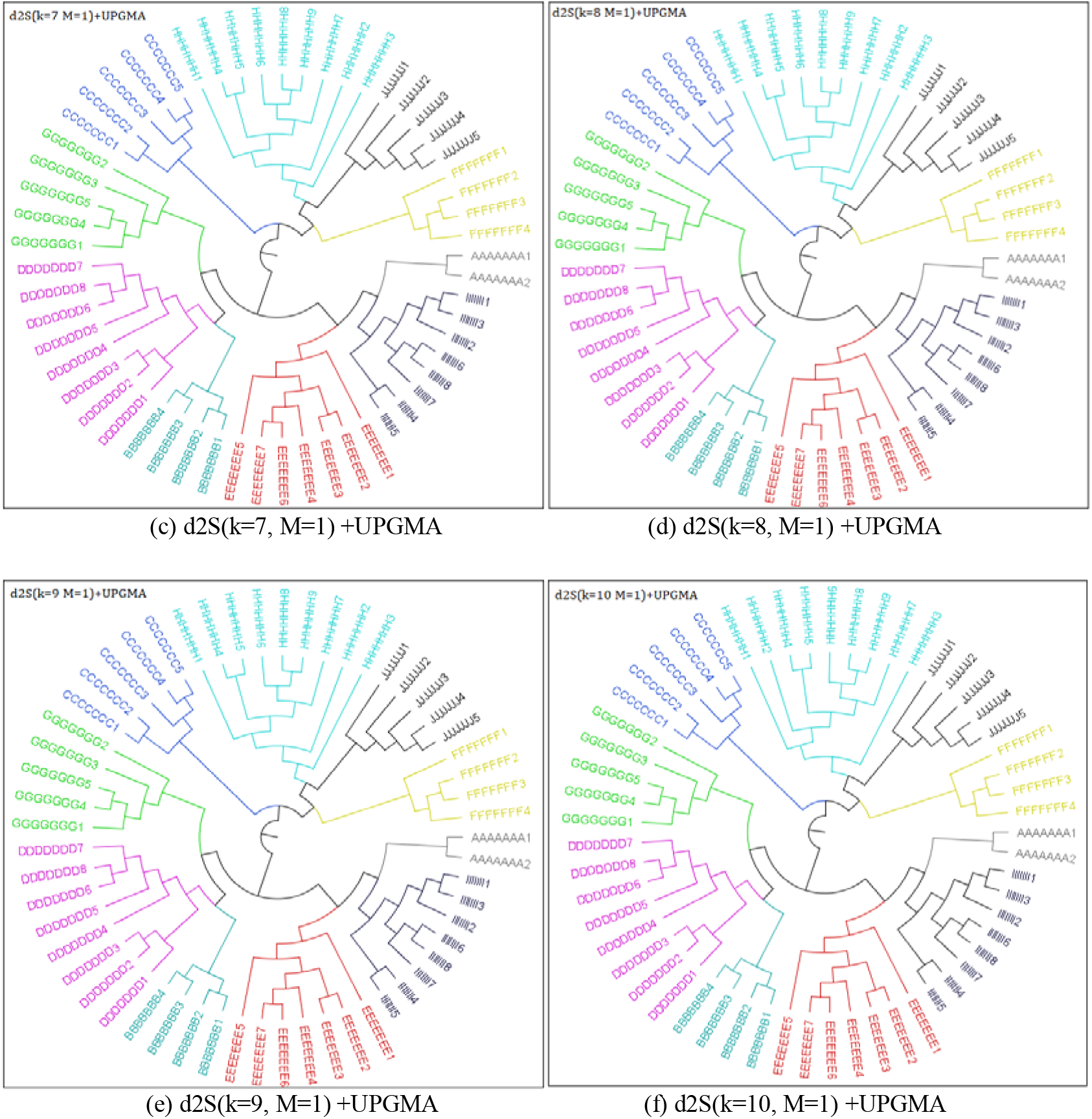
Phylogenetic trees for 16S rRNA sequences(57 sequences)via d2S (k=4, 6, 7, 8, 9, 10, M=1)

**Fig.9.**
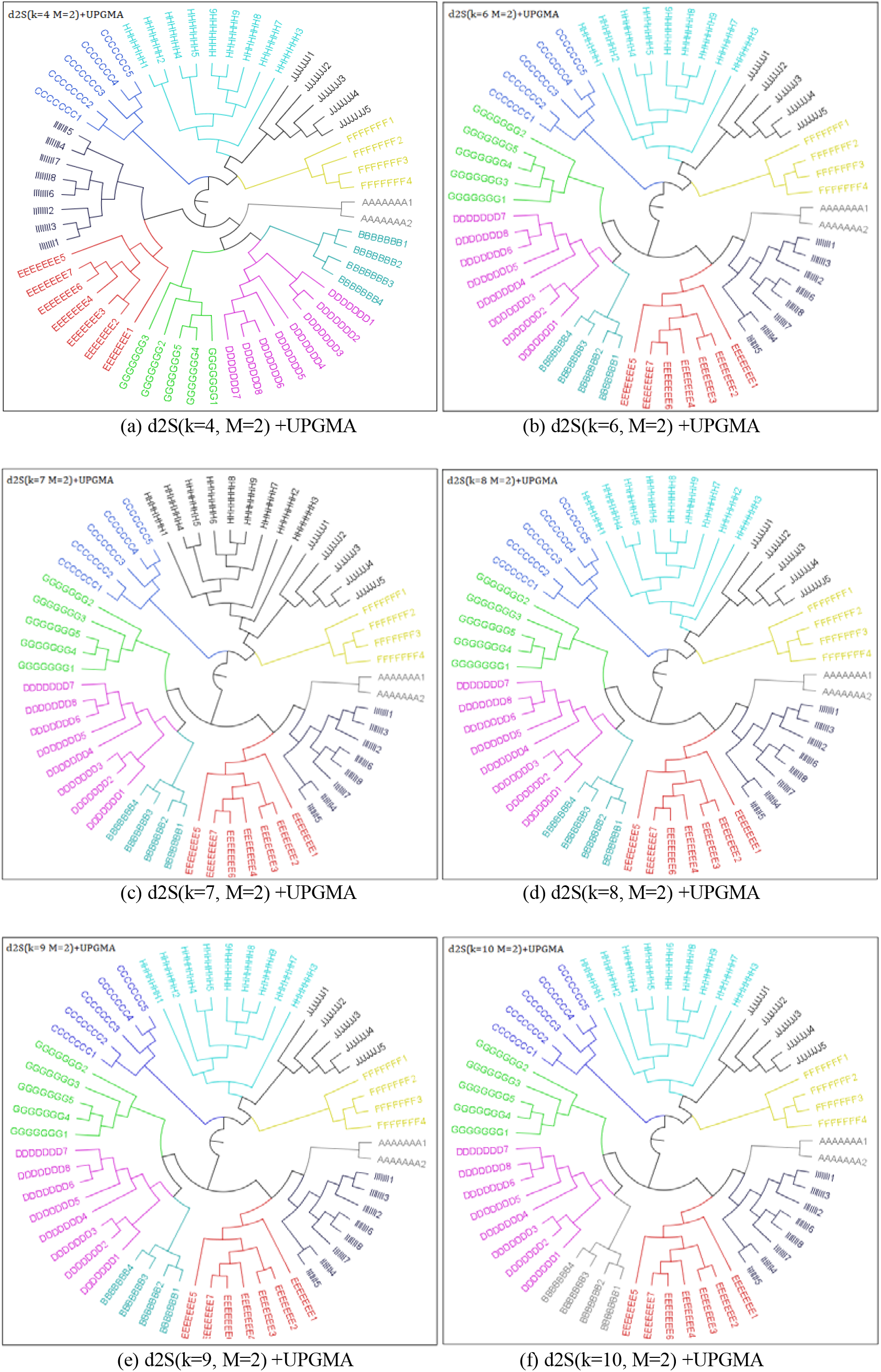
Phylogenetic trees for 16S rRNA sequences(57 sequences)via d2S (k=4, 6, 7, 8, 9, 10, M=2)

**Fig.10.**
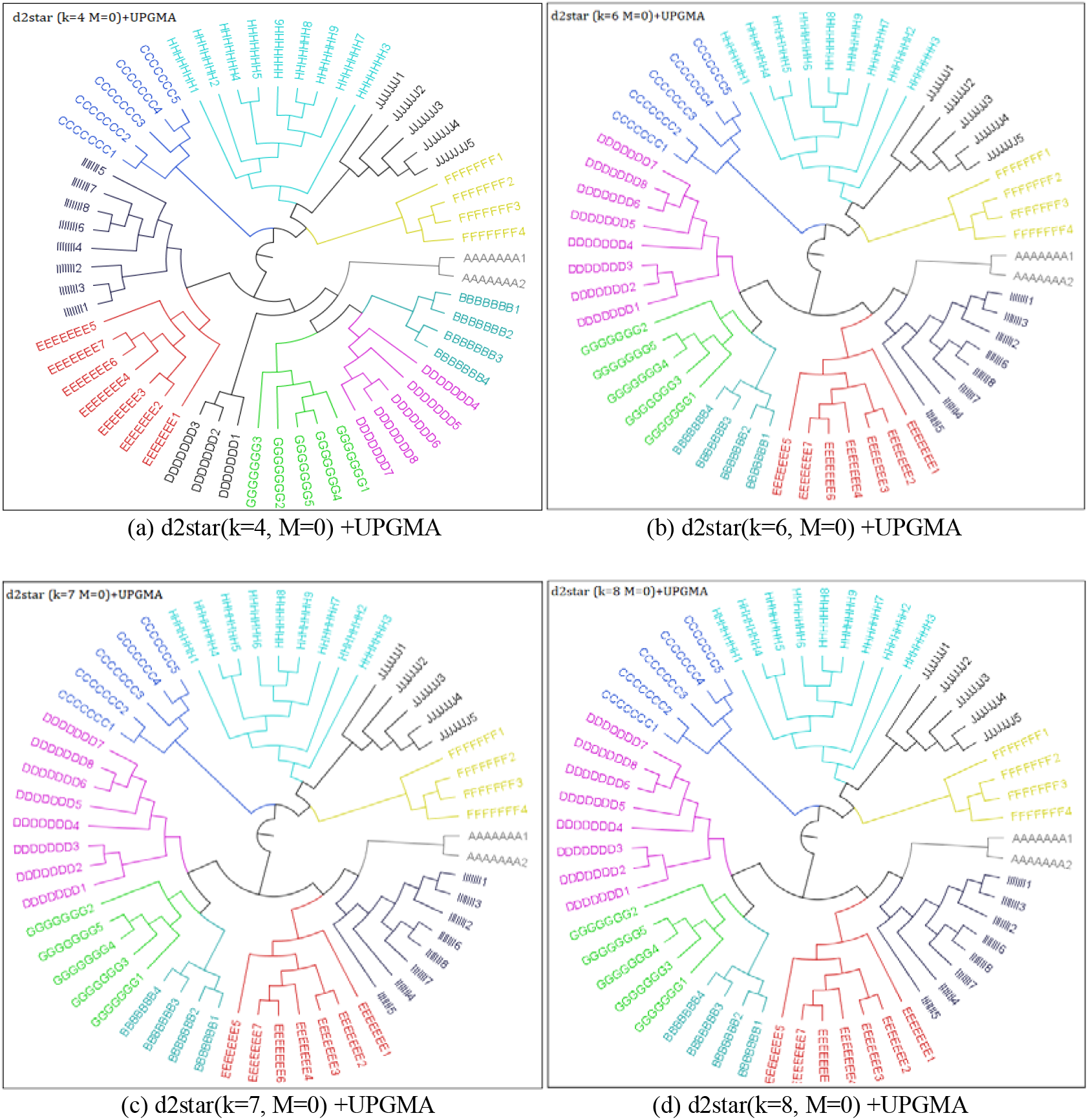
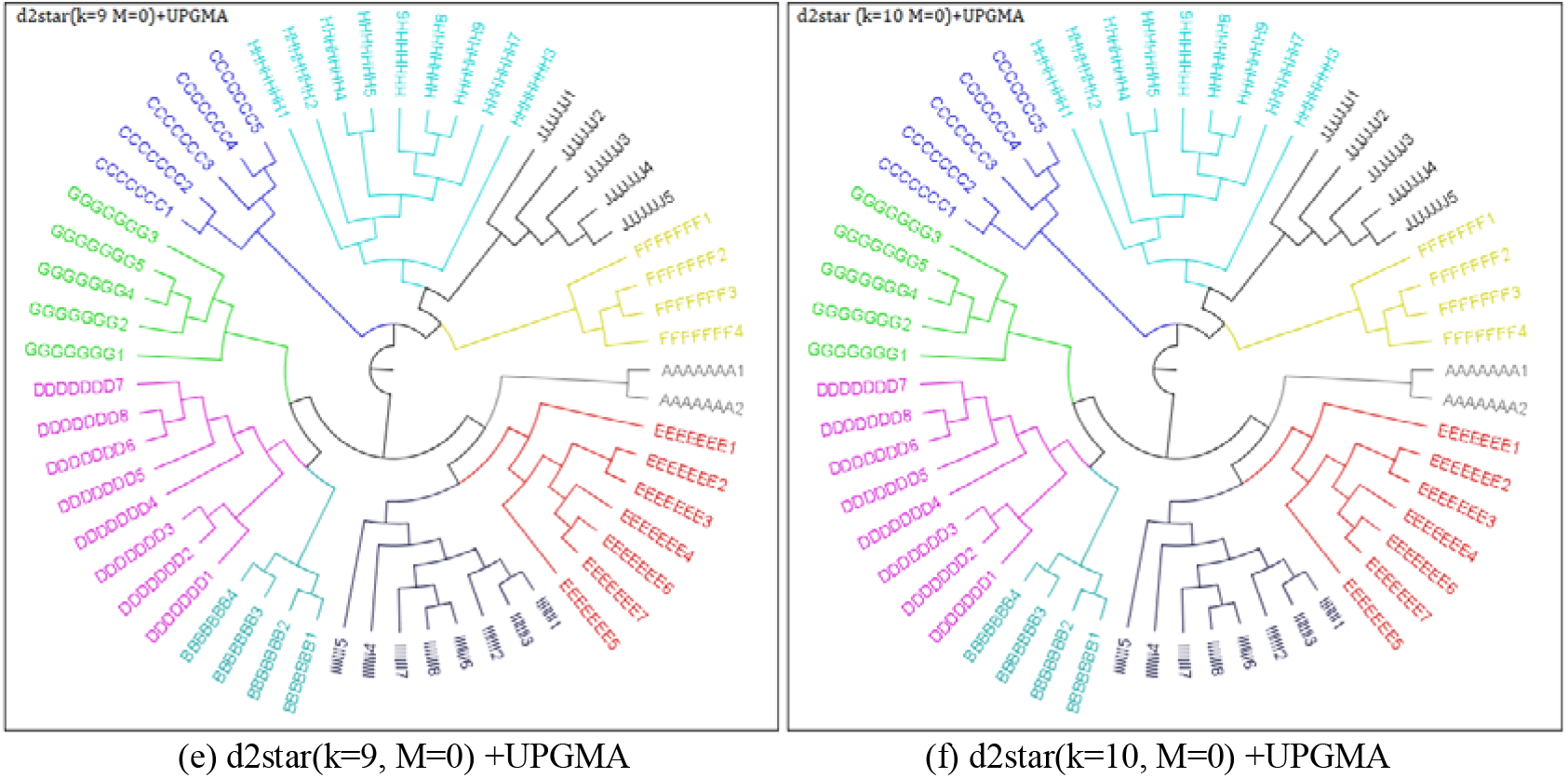
Phylogenetic trees for 16S rRNA sequences(57 sequences)via d2star (k=4, 6, 7, 8, 9, 10, M=0)

**Fig.11.**
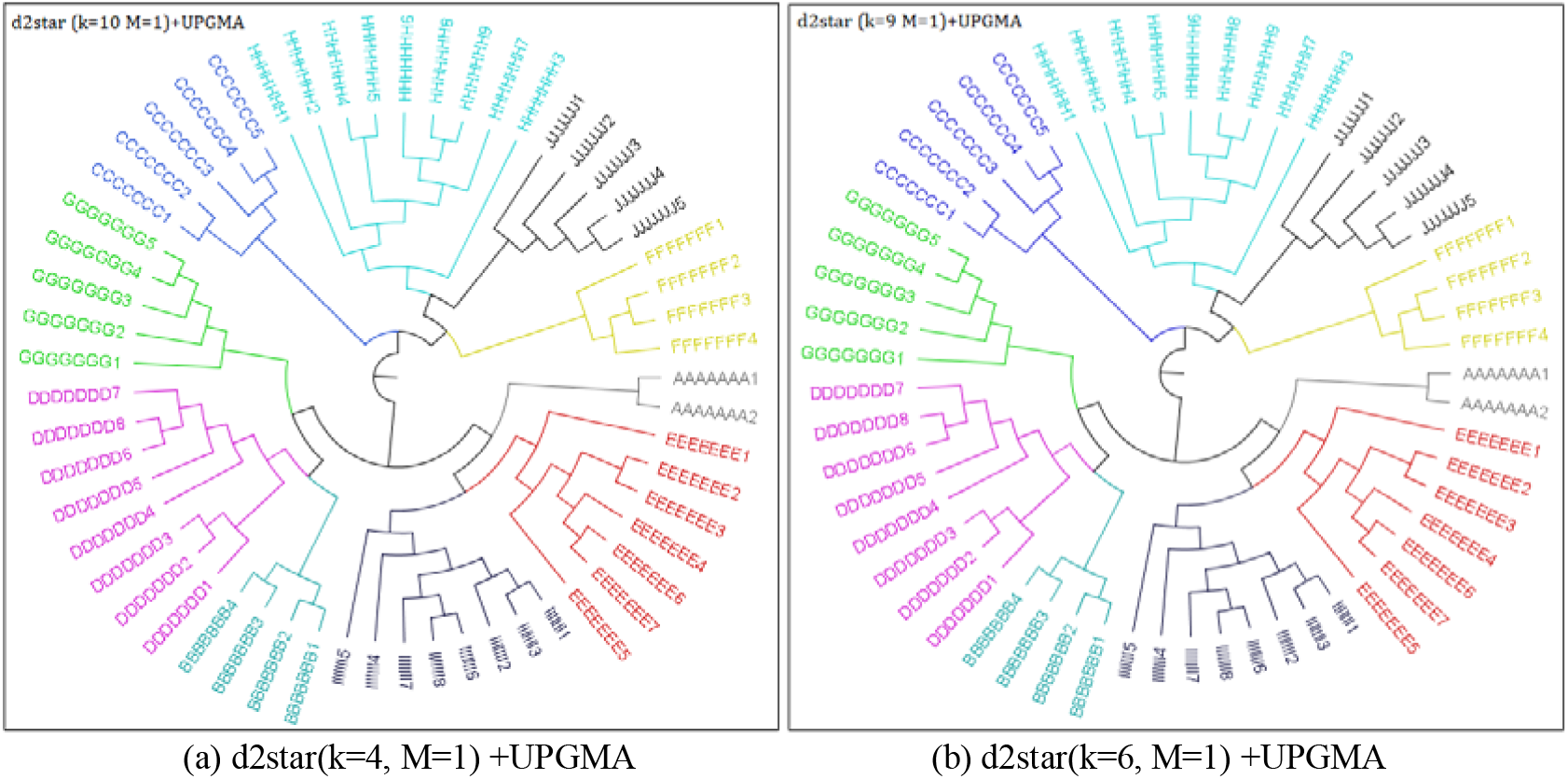
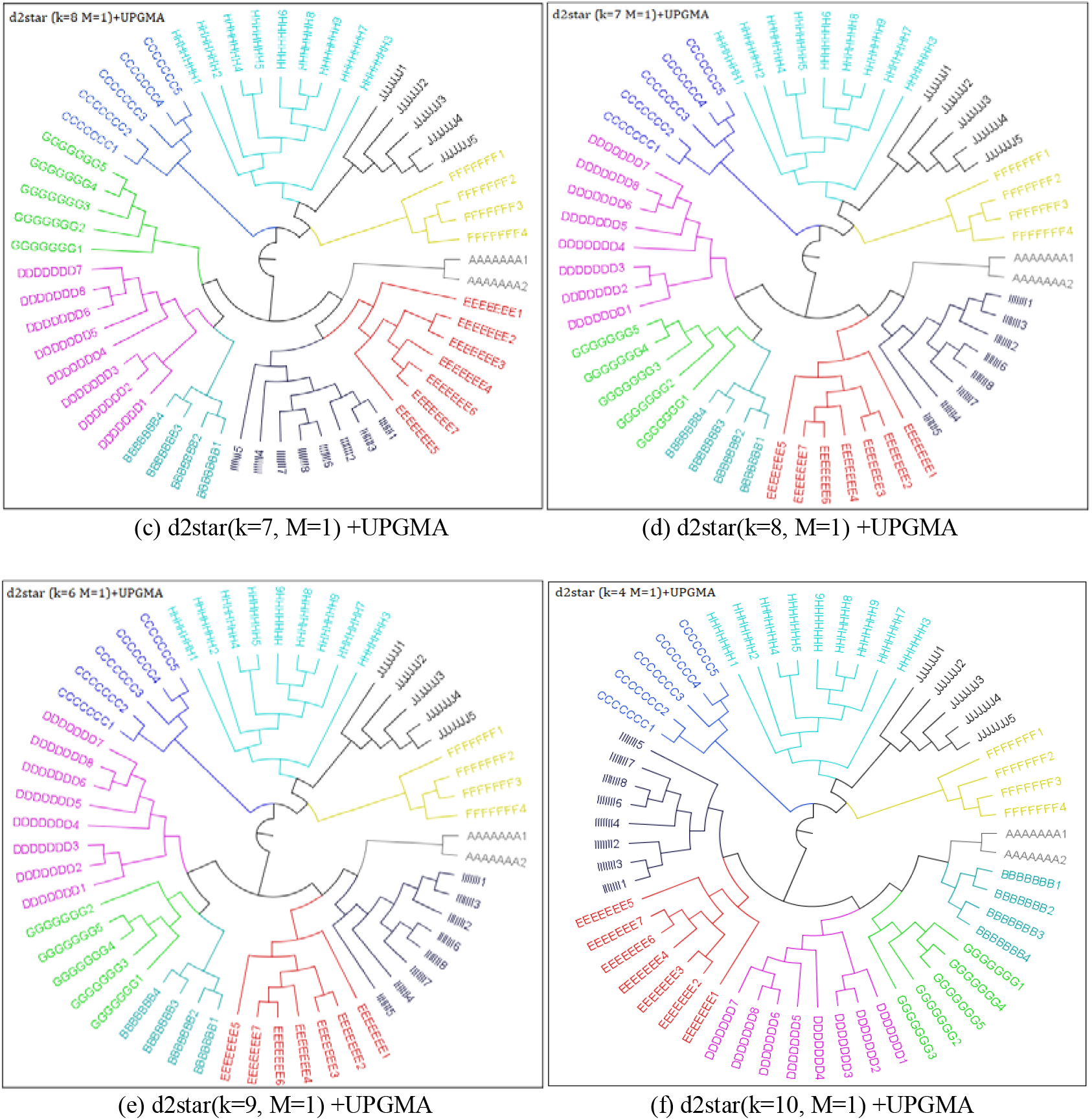
Phylogenetic trees for 16S rRNA sequences(57 sequences)via d2star (k=4, 6, 7, 8, 9, 10, M=1)

**Fig.12.**
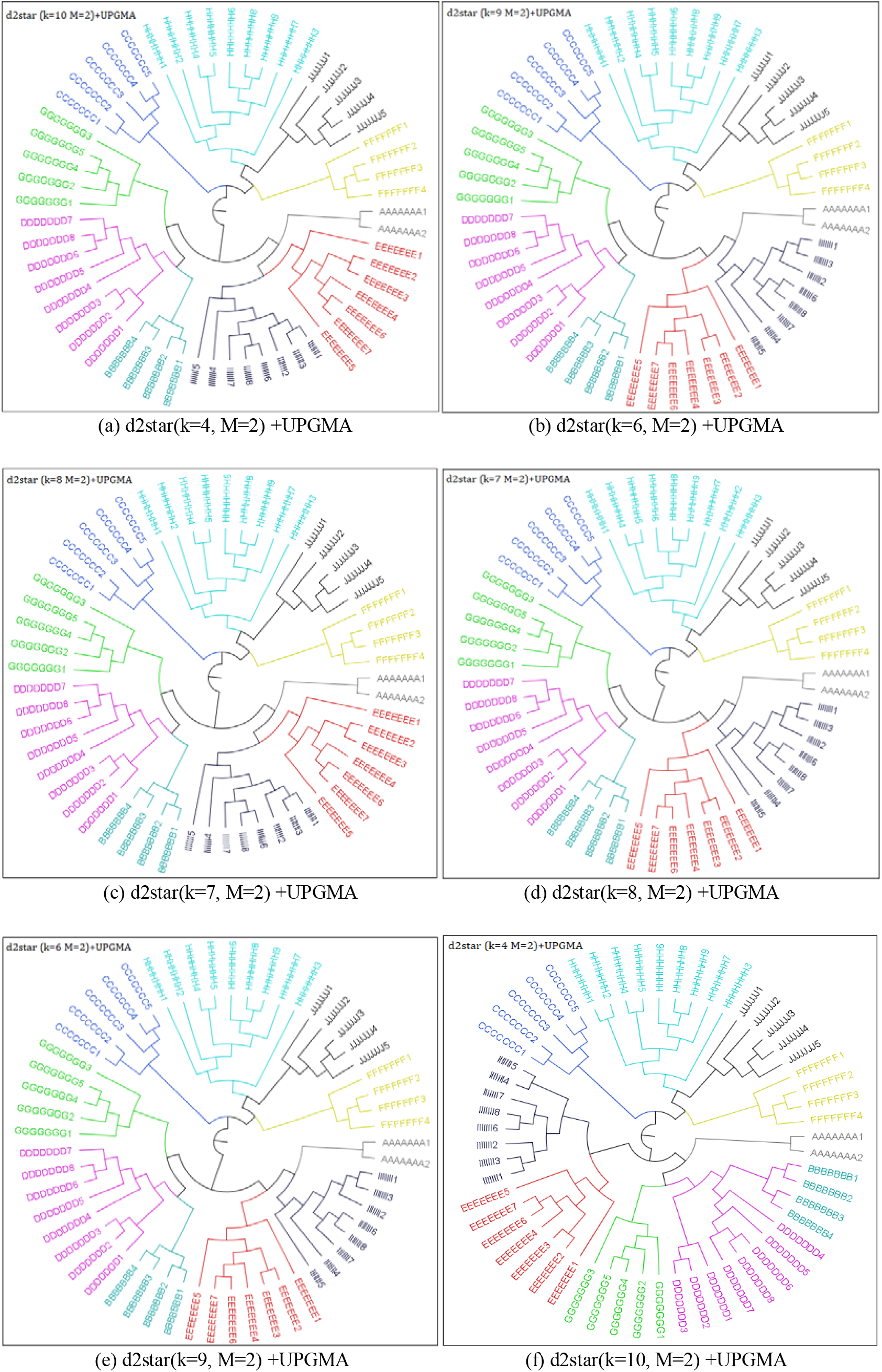
Phylogenetic trees for 16S rRNA sequences(57 sequences)via d2star (k=4, 6, 7, 8, 9, 10, M=2)

## References

[1] F. Delsuc, B. Henner and P. Herve. Phylogenomics and the reconstruction of the tree of life. Nature Reviews Genetics, 6 (5): 361–375, 2005.

[2] J. A. Eisen and C.M. Fraser. Phylogenomics: intersection of evolution and genomics. Science, 300(5626): 1706–1707, 2003.

[3] H. Carrillo and D. Lipman. The multiple sequence alignment problem in biology. Siam Journal on Applied Mathematics, 48(5): 1073–1082, 1988.

[4] W. G. Weisburg, S. M. Barns, D. A. Pelletier and D. J. Lane. 16s Ribosomal DNA Amplification for Phylogenetic Study. Journal of Bacteriology, 173(2): 697–703, 1991.

[5] D. F. Feng, and R. F. Doolittle. Progressive sequence alignment as a prerequisite to correct phylogenetic trees. Journal of molecular evolution, 25(4): 351–360, 1987.

[6] M. Zhang, L. Yang, J. Ren, N.A. Ahlgren, J.A. Fuhrman and F.Z. Sun. Prediction of virus-host infectious associ-ation by supervised learning methods, BMC Bioinformatics, 18(3):60, 2017.

[7] Y. Y. Lu, K. Tang, J. Ren, J. A. Fuhrman, M. S. Waterman and F. Z. Sun. CAFE: aCcelerated Alignment-FrEe sequence analysis, Nucleic Acids Research, 45(W1), 2017.

[8] M. A. Ragan, G. Bernard and C.X. Chan. Molecular phylogenetics before sequences: oligonucleotide catalogs as k-mer spectra. RNA Biology, 11(3): 176–185, 2014.

[9] D. T. Pride, R. J. Meinersmann, T. M. Wassenaar and M. J. Blaser. Evolutionary implications of microbial genome tetra nucleotide frequency biase. Genome Research, 13(2): 145–158, 2003.

[10] R. T. Miller. A comprehensive approach to clustering of expressed human gene sequence: the sequence tag alignment and consensus knowledge base. Genome Research, 9(11): 1143–1155, 1999.

[11] F. Huan, A. R. Ives, Y. Surget-Groba and C. H. Cannon. An assembly and alignment-free method of phylogeny reconstruction from next-generation sequencing data. BMC Genomics, 16(1): 522(1-18), 2015.

[12] C. X. Chan, G. Bernard, O. Poirion, J. M. Hogan and M. A. Ragan. Inferring phylogenies of evolving sequences without multiple sequence alignment. Scientific Reports, 4(1): 6504, 2014.

[13] G. Cattaneo, U. F. Petrillo, R. Giancarlo and G. Roscigno. An effective extension of the applicability of alignment-free biological sequence comparison algorithms with hadoop. The Journal of Supercomputing, 73(4): 1467–1483, 2017.

[14] Z. Xu and B. L. Hao. CVTree update: a newly designed phylogenetic study platform using composition vectors and whole genomes. Nucleic Acids Research, 37: W174–178, 2009.

[15] J. Qi, H. Luo and B. L. Hao. CVTree: a phylogenetic tree reconstruction tool based on whole genomes. Nucleic Acids Research, 32: W45–W47, 2004.

[16] J. Qi, B. Wang and B. I Hao. Whole proteome prokaryote phylogeny without sequence alignment: a k-string composition approach. Journal of Molecular Evolution, 58(1):1–11, 2004.

[17] H. H. Huang. An ensemble distance measure of k-mer and Natural Vector for the phylogenetic analysis of multiple-segmented viruses, Journal of Theoretical Biology 398:136–144, 2016.

[18] J. Wen., R. H. Chan., R. L. He and S. S. Yau. The k-mer natural vector and its application to the phylogenetic analysis of genetic sequences, Gene, 546(1):25–34, 2014.

[19] R. A. Lippert, H. Huang and M. S. Waterman. Distributional regimes for the number of kword matches between two random sequences. Proceedings of the National Academy of Sciences of the Unite States of America, 99(22): 13980–13989, 2002.

[20] K. Song, J. Ren, G. Reinert, M. H. Deng, M. S. Waterman and F. Z. Sun. New developments of alignment-free sequence comparison: measures, statistics and next-generation sequencing. Briefings in Bioinformatics 15(3): 343–353, 2013.

[21] N. A. Ahlgren, J. Ren, Y. Y. Lu, J. A. Fuhrman and F. Z. Sun. Alignment-free d2* oligonucleotide frequency dissimilarity measure improves prediction of hosts from metagenomically-derived viral sequences. Nucleic Acids Research, 45(1): 39–53, 2017.

[22] R. Sokal and C. Michener. “A statistical method for evaluating systematic relationships”. University of Kansas Science Bulletin. 38: 1409–1438, 1958.

[23] R. C. Edgar. MUSCLE: a multiple sequence alignment method with reduced time and space complexity, BMC Bioinformatics, 5:113, 2004.

[24] R. Kumar, G. Stecher, M. Li, C. Knyaz and K. Tamura. MEGA X: Molecular Evolutionary Genetics Analysis across computing platforms, Molecular Biology and Evolution, 35:1547–1549, 2018.

[25] S. Foret, S. R. Wilson and C. J. Burden. Characterizing the D2 statistic: word matches in biological sequences. Statistical Applications in Genetics and Molecular Biology, 8(1): 1–21, 2009.

[26] C. R. Woese and G. E. Fox. Phylogenetic structure of the prokaryotic domain: the primary kingdoms. Proceedings of the National Academy of Sciences of the Unite States of America, 74(11): 5088–5090, 1977.

[27] C. R Woese. Bacterial evolution. Microbiological Reviews, 51:221, 1987.

[28] C. Quast, E. Pruesse, P. Yilmaz, J. Gerken, T. Schweer, P. Yarza, J. Peplies and F. O. Glöckner. The SILVA ribosomal RNA gene database project: improved data processing and web-based tools. Nucleic Acids Research, 41 (D1): D590–D596, 2012.

[29] P. Yilmaz, L. W. Parfrey, P. Yarza, J. Gerken, E. Pruesse, C. Quast, T. Schweer, J. Peplies, W. Ludwig and F. O. Glöckner. The SILVA and “All-species Living Tree Project (LTP)” taxonomic frameworks. Nucleic Acids Research, 42(D1): D643–D648, 2014.

[30] F. O. Glöckner, P. Yilmaz, C. Quast, J. Gerken, A. Beccati, A. Ciuprina, G. Bruns, P. Yarza, J. Peplies, R. Westram and W. Ludwig. 25 years of serving the community with ribosomal RNA gene reference databases and tools. Journal of Biotechnology, 261:169–176, 2017.

[31] A. Beccati, J. Gerken, C. Quast, P. Yilmaz and F. O. Glöckner. SILVA tree viewer: interactive web browsing of the SILVA phylogenetic guide trees. BMC Bioinformatics, 18(1): 433–437, 2017.

[32] D. R. Robinson and L. R. Foulds. Comparison of phylogenetic trees. Mathematical Bioscience, 53:131–147, 1981.

