## Supplementary Information for "SeqDistK: a Novel Tool for Alignment-free Phylogenetic Analysis"

(1) **Working interface**

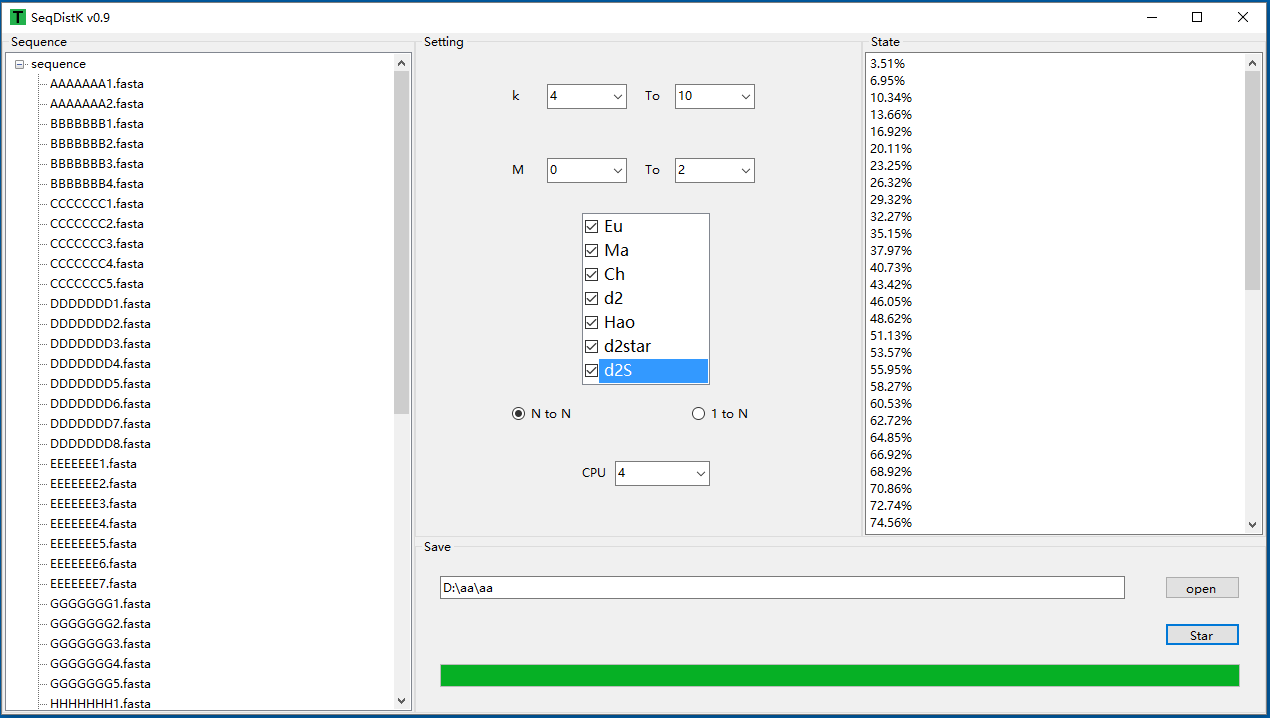

Fig.1 The working interface of SeqDistK

SeqDistK is a tool to calculate the distance among sequences and the window interface as Fig.1. There are 7 dissimilarity measures Eu, Ma, Ch, d2, d2star, d2S and Hao in SeqDistK. The software supports Windows. The advantage of the software is convenience. The whole process can be operated by mouse. It supports multiple directory-unit and multilevel directory structure and each directory will calculate independently. The results will be saved in directory structures independently.

(2) **Supplementary Tables**

Table 1. Data sets of 16S rRNA sequences（57 sequences）for standard tree

| Number | NCBIName | Type | Name |
| --- | --- | --- | --- |
| AAAAAAA1 | JQ347593 | Bacterium | Mesoaciditoga lauensis |
| AAAAAAA2 | AB8958788 | Bacterium | Thermotogales bacterium NAS-01 |
| BBBBBBB1 | FR733692 | Acetomicrobium | Acetomicrobium flavidum |
| BBBBBBB2 | AB910748 | Anaerobaculum | Anaerobaculum mobile |
| BBBBBBB3 | U50711 | Anaerobaculum | Anaerobaculum thermoterrenum |
| BBBBBBB4 | FJ862996 | Anaerobaculum | Anaerobaculum hydrogeniformans |
| CCCCCCC1 | AF262035 | Methanofollis | Methanofollis sp |
| CCCCCCC2 | AY186542 | Methanofollis | Methanofollis formosanus |
| CCCCCCC3 | AB371073 | Methanofollis | Methanofollis ethanolicus |
| CCCCCCC4 | Y16428 | Methanofollis | Methanofollis liminatans |
| CCCCCCC5 | AF095272 | Methanofollis | Methanofollis tationis |
| DDDDDDD1 | EF436500 | Desulfothiovibrio | Jonquetella anthropi strain ADV126 |
| DDDDDDD2 | EF468685 | Desulfothiovibrio | Rarimicrobium hominis strain ADV70 |
| DDDDDDD3 | EU309492 | Desulfothiovibrio | Pyramidobacter piscolens W5455 |
| DDDDDDD4 | EU719657 | Desulfothiovibrio | Dethiosulfovibrio salsuginis strain USBA |
| DDDDDDD5 | U52817 | Desulfothiovibrio | Desulfothiovibrio peptidovorans |
| DDDDDDD6 | AY005466 | Desulfothiovibrio | Dethiosulfovibrio acidaminovorans sr15 |
| DDDDDDD7 | AF234544 | Desulfothiovibrio | Dethiosulfovibrio marinus strain WS100 |
| DDDDDDD8 | AF234542 | Desulfothiovibrio | Dethiosulfovibrio russensis strain sr12 |
| EEEEEEE1 | FR850164 | Defluviitoga | Defluviitoga tunisiensis partial |
| EEEEEEE2 | AJ311702 | Petrotoga | Petrotoga siberica |
| EEEEEEE3 | AJ311703 | Petrotoga | Petrotoga olearia |
| EEEEEEE4 | AY125964 | Petrotoga | Petrotoga mexicana |
| EEEEEEE5 | Y15479 | Petrotoga | Petrotoga mobilis |
| EEEEEEE6 | FR733705 | Petrotoga | Petrotoga miotherma |
| EEEEEEE7 | AY800102 | Petrotoga | Petrotoga halophila strain MET-B |
| FFFFFFF1 | HM159601 | laminariae | Salinarchaeum laminariae strain R26 |
| FFFFFFF2 | AB457580 | Natronoarchaeum | Natronoarchaeum philippinense |
| FFFFFFF3 | AB501361 | Natronoarchaeum | Natronoarchaeum mannanilyticum |
| FFFFFFF4 | JF421970 | Natronoarchaeum | Natronoarchaeum rubrum strain GX48 |
| GGGGGGG1 | AB011495 | Thermaerobacter | Aerothermobacter marianas |
| GGGGGGG2 | AB454087 | Thermaerobacter | Thermaerobacter composti |
| GGGGGGG3 | AY936496 | Thermaerobacter | Thermaerobacter litoralis strain KW1 |
| GGGGGGG4 | AB061441 | Thermaerobacter | Thermaerobacter nagasakiensis |
| GGGGGGG5 | AF343566 | Thermaerobacter | Thermaerobacter subterraneus |
| HHHHHHH1 | GQ282620 | Halobellus | Halobellus clavatus strain TNN18 |
| HHHHHHH2 | HQ451075 | Halobellus | Halobellus salinus strain CSW2 |
| HHHHHHH3 | AY676200 | Halobellus | Haloquadratum walsbyi strain C23 |
| HHHHHHH4 | JQ237122 | Halobellus | Halobellus inordinatus strain YC20 |
| HHHHHHH5 | JQ910929 | Halobellus | Halobellus ramosii strain S2FP14 |
| HHHHHHH6 | GU208828 | Halobellus | Halobellus limi strain TBN53 |
| HHHHHHH7 | KF314040 | Halobellus | Halobellus rufus strain CBA1103 |
| HHHHHHH8 | GU951426 | Halobellus | Halobellus litoreus strain GX31 |
| HHHHHHH9 | JQ237123 | Halobellus | Halobellus rarus strain YC21 |
| IIIIIII1 | KF931642 | Thermosipho | Thermosipho activus strain Rift-s3 |
| IIIIIII2 | AJ272022 | Thermosipho | Thermosipho sp. DSM 13256 |
| IIIIIII3 | AJ577471 | Thermosipho | Thermosipho atlanticus |
| IIIIIII4 | Z70248 | Thermosipho | Thermosipho melanesiensis |
| IIIIIII5 | GQ292553 | Thermosipho | Thermosipho affectus strain ik275mar |
| IIIIIII6 | AB257289 | Thermosipho | Thermosipho globiformans |
| IIIIIII7 | AB024932 | Thermosipho | Thermosipho japonicus |
| IIIIIII8 | DQ647057 | Thermosipho | Thermosipho africanus strain |
| JJJJJJJ1 | JQ937359 | Haloarchaeobius | Haloarchaeobius amylolyticus strain XD48 |
| JJJJJJJ2 | LC061270 | Haloarchaeobius | Haloarchaeobius baliensis |
| JJJJJJJ3 | JF293278 | Haloarchaeobius | Haloarchaeobius iranensis strain EB21 |
| JJJJJJJ4 | GU951428 | Haloarchaeobius | Haloarchaeobius litoreus strain GX60 |
| JJJJJJJ5 | JQ937361 | Haloarchaeobius | Haloarchaeobius salinus strain YC82 |

Table 2. Data sets of 16S rRNA sequences（63sequences）for clustering

| **Sequence** | **Domaim** | **Phylum** | **Class** | **Order** | **Family** |
| --- | --- | --- | --- | --- | --- |
| **AB603516** | **Archaea** | **Euryarchaeota** | **Methanococci** | **Methanococcales** | **Methanocaldococcaceae** |
| **DQ228625** | **Archaea** | **Euryarchaeota** | **Methanococci** | **Methanococcales** | **Methanocaldococcaceae** |
| **AF051404** | **Archaea** | **Euryarchaeota** | **Methanococci** | **Methanococcales** | **Methanocaldococcaceae** |
| **AF056938** | **Archaea** | **Euryarchaeota** | **Methanococci** | **Methanococcales** | **Methanocaldococcaceae** |
| **AF356634** | **Archaea** | **Euryarchaeota** | **Methanococci** | **Methanococcales** | **Methanocaldococcaceae** |
| **AJ969471** | **Archaea** | **Euryarchaeota** | **Methanococci** | **Methanococcales** | **Methanocaldococcaceae** |
| **AJ969469** | **Archaea** | **Euryarchaeota** | **Methanococci** | **Methanococcales** | **Methanocaldococcaceae** |
| **AB235312** | **Archaea** | **Euryarchaeota** | **Methanococci** | **Methanococcales** | **Methanocaldococcaceae** |
| **AF025822** | **Archaea** | **Euryarchaeota** | **Methanococci** | **Methanococcales** | **Methanocaldococcaceae** |
| **AJ969473** | **Archaea** | **Euryarchaeota** | **Methanococci** | **Methanococcales** | **Methanocaldococcaceae** |
| **FJ766848** | **Archaea** | **Euryarchaeota** | **Methanococci** | **Methanococcales** | **Methanocaldococcaceae** |
| **AF547621** | **Archaea** | **Euryarchaeota** | **Methanococci** | **Methanococcales** | **Methanocaldococcaceae** |
| **AY264344** | **Archaea** | **Crenarchaeota** | **Thermoprotei** | **Desulfurococcales** | **Desulfurococcaceae** |
| **AB661712** | **Archaea** | **Crenarchaeota** | **Thermoprotei** | **Desulfurococcales** | **Desulfurococcaceae** |
| **EU167539** | **Archaea** | **Crenarchaeota** | **Thermoprotei** | **Desulfurococcales** | **Desulfurococcaceae** |
| **HG796148** | **Archaea** | **Crenarchaeota** | **Thermoprotei** | **Desulfurococcales** | **Desulfurococcaceae** |
| **AB462558** | **Archaea** | **Crenarchaeota** | **Thermoprotei** | **Desulfurococcales** | **Desulfurococcaceae** |
| **AB293243** | **Archaea** | **Crenarchaeota** | **Thermoprotei** | **Desulfurococcales** | **Desulfurococcaceae** |
| **KF278498** | **Archaea** | **Crenarchaeota** | **Thermoprotei** | **Desulfurococcales** | **Desulfurococcaceae** |
| **KF278499** | **Archaea** | **Crenarchaeota** | **Thermoprotei** | **Desulfurococcales** | **Desulfurococcaceae** |
| **X99560** | **Archaea** | **Crenarchaeota** | **Thermoprotei** | **Desulfurococcales** | **Desulfurococcaceae** |
| **AJ012645** | **Archaea** | **Crenarchaeota** | **Thermoprotei** | **Desulfurococcales** | **Desulfurococcaceae** |
| **X99562** | **Archaea** | **Crenarchaeota** | **Thermoprotei** | **Desulfurococcales** | **Ignicoccaceae** |
| **AJ271794** | **Archaea** | **Crenarchaeota** | **Thermoprotei** | **Desulfurococcales** | **Ignicoccaceae** |
| **HK556290** | **Archaea** | **Crenarchaeota** | **Thermoprotei** | **Desulfurococcales** | **Ignicoccaceae** |
| **AJ318042** | **Archaea** | **Crenarchaeota** | **Thermoprotei** | **Desulfurococcales** | **Ignicoccaceae** |
| **DQ060321** | **Archaea** | **Crenarchaeota** | **Thermoprotei** | **Desulfurococcales** | **Ignicoccaceae** |
| **DQ060322** | **Archaea** | **Crenarchaeota** | **Thermoprotei** | **Desulfurococcales** | **Ignicoccaceae** |
| **DQ060320** | **Archaea** | **Crenarchaeota** | **Thermoprotei** | **Desulfurococcales** | **Ignicoccaceae** |
| **JF509453** | **Archaea** | **Crenarchaeota** | **Thermoprotei** | **Desulfurococcales** | **Ignicoccaceae** |
| **DQ243730** | **Archaea** | **Crenarchaeota** | **Thermoprotei** | **Desulfurococcales** | **Ignicoccaceae** |
| **HM448086** | **Archaea** | **Crenarchaeota** | **Thermoprotei** | **Desulfurococcales** | **Ignicoccaceae** |
| **EU530582** | **Archaea** | **Crenarchaeota** | **Thermoprotei** | **Desulfurococcales** | **Ignicoccaceae** |
| **AB462559** | **Archaea** | **Crenarchaeota** | **Thermoprotei** | **Desulfurococcales** | **Ignicoccaceae** |
| **JF935165** | **Archaea** | **Crenarchaeota** | **Thermoprotei** | **Desulfurococcales** | **Ignicoccaceae** |
| **EU530578** | **Archaea** | **Crenarchaeota** | **Thermoprotei** | **Desulfurococcales** | **Ignicoccaceae** |
| **BD445501** | **Archaea** | **Crenarchaeota** | **Thermoprotei** | **Desulfurococcales** | **Ignicoccaceae** |
| **DQ243732** | **Archaea** | **Crenarchaeota** | **Thermoprotei** | **Desulfurococcales** | **Ignicoccaceae** |
| **KU664659** | **Bacteria** | **Firmicutes** | **Clostridia** | **Thermoanaerobacterales** | **Thermodesulfobiaceae** |
| **JQ815731** | **Bacteria** | **Firmicutes** | **Clostridia** | **Thermoanaerobacterales** | **Thermodesulfobiaceae** |
| **DQ834002** | **Bacteria** | **Firmicutes** | **Clostridia** | **Thermoanaerobacterales** | **Thermodesulfobiaceae** |
| **AB077817** | **Bacteria** | **Firmicutes** | **Clostridia** | **Thermoanaerobacterales** | **Thermodesulfobiaceae** |
| **CU918272** | **Bacteria** | **Thermotogae** | **Thermotogae** | **Kosmotogales** | **Kosmotogaceae** |
| **CU923378** | **Bacteria** | **Thermotogae** | **Thermotogae** | **Kosmotogales** | **Kosmotogaceae** |
| **GU180074** | **Bacteria** | **Thermotogae** | **Thermotogae** | **Kosmotogales** | **Kosmotogaceae** |
| **KJ881256** | **Bacteria** | **Thermotogae** | **Thermotogae** | **Kosmotogales** | **Kosmotogaceae** |
| **AB853916** | **Bacteria** | **Thermotogae** | **Thermotogae** | **Kosmotogales** | **Kosmotogaceae** |
| **EF515526** | **Bacteria** | **Thermotogae** | **Thermotogae** | **Kosmotogales** | **Kosmotogaceae** |
| **FR775407** | **Bacteria** | **Thermotogae** | **Thermotogae** | **Kosmotogales** | **Kosmotogaceae** |
| **KJ206811** | **Bacteria** | **Thermotogae** | **Thermotogae** | **Kosmotogales** | **Kosmotogaceae** |
| **FJ645709** | **Bacteria** | **Thermotogae** | **Thermotogae** | **Kosmotogales** | **Kosmotogaceae** |
| **CU917527** | **Bacteria** | **Thermotogae** | **Thermotogae** | **Kosmotogales** | **Kosmotogaceae** |
| **EU999020** | **Bacteria** | **Thermotogae** | **Thermotogae** | **Petrotogales** | **Petrotogaceae** |
| **AB369055** | **Bacteria** | **Thermotogae** | **Thermotogae** | **Petrotogales** | **Petrotogaceae** |
| **FR733705** | **Bacteria** | **Thermotogae** | **Thermotogae** | **Petrotogales** | **Petrotogaceae** |
| **AY800102** | **Bacteria** | **Thermotogae** | **Thermotogae** | **Petrotogales** | **Petrotogaceae** |
| **EU573091** | **Bacteria** | **Thermotogae** | **Thermotogae** | **Petrotogales** | **Petrotogaceae** |
| **AJ311702** | **Bacteria** | **Thermotogae** | **Thermotogae** | **Petrotogales** | **Petrotogaceae** |
| **AY125964** | **Bacteria** | **Thermotogae** | **Thermotogae** | **Petrotogales** | **Petrotogaceae** |
| **JF808037** | **Bacteria** | **Thermotogae** | **Thermotogae** | **Petrotogales** | **Petrotogaceae** |
| **Y15479** | **Bacteria** | **Thermotogae** | **Thermotogae** | **Petrotogales** | **Petrotogaceae** |
| **GU180075** | **Bacteria** | **Thermotogae** | **Thermotogae** | **Petrotogales** | **Petrotogaceae** |
| **GU180071** | **Bacteria** | **Thermotogae** | **Thermotogae** | **Petrotogales** | **Petrotogaceae** |

(3) **Supplementary Figures**

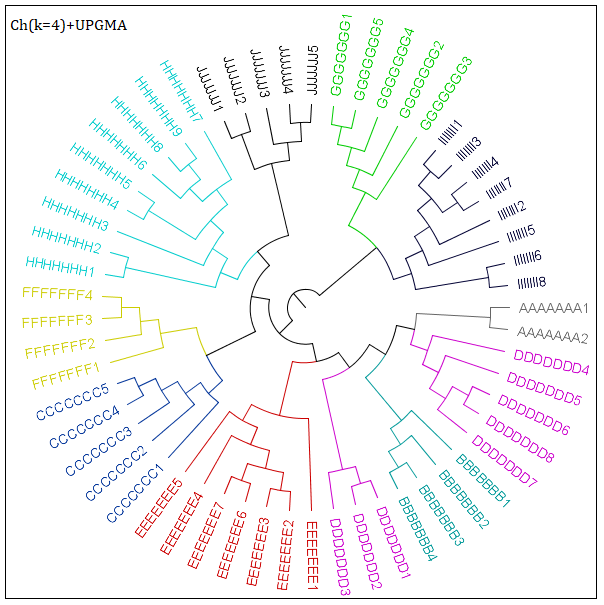

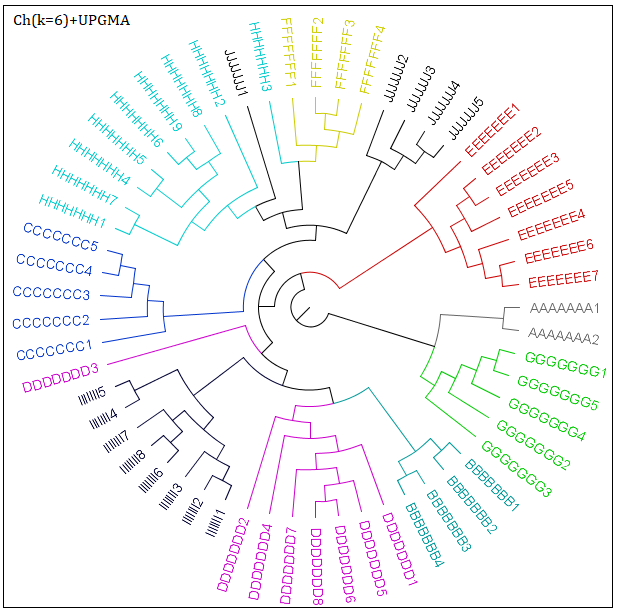

(a) Ch(k=4) +UPGMA	(b) Ch(k=6) +UPGMA

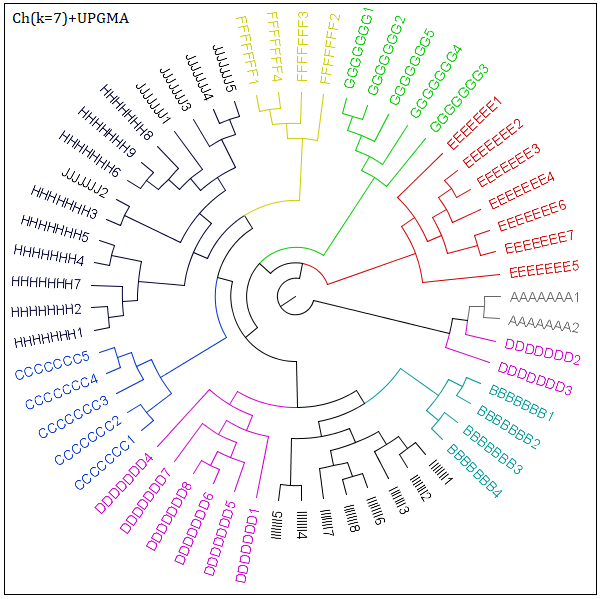

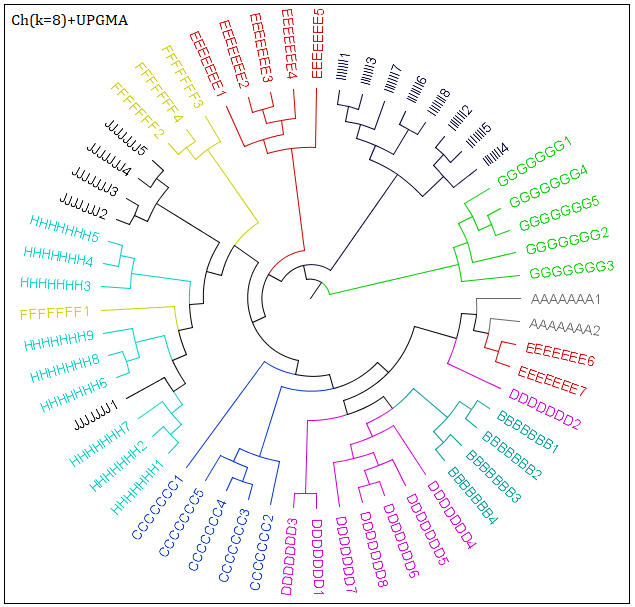

(c) Ch(k=7) +UPGMA	(d) Ch(k=8) +UPGMA

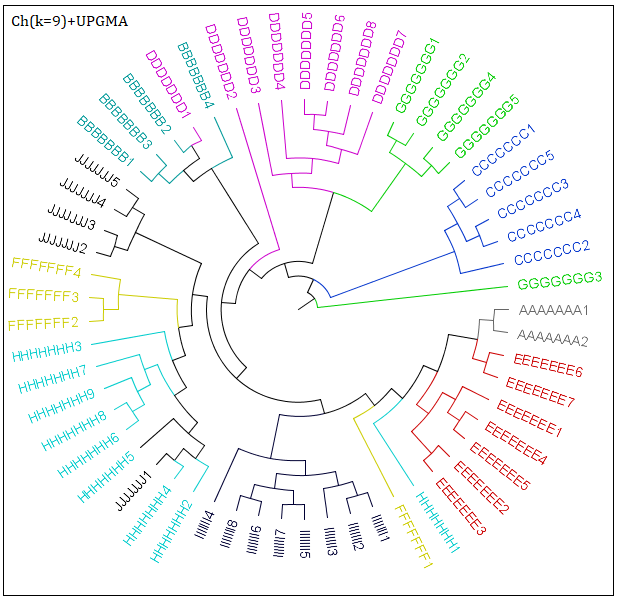

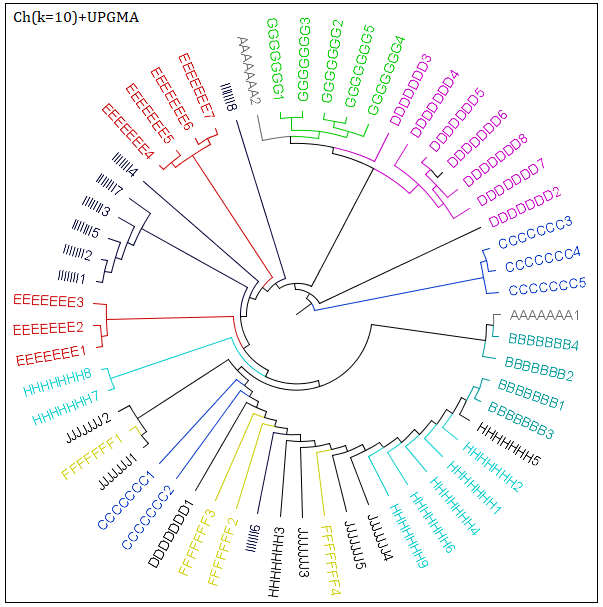

(e) Ch(k=9) +UPGMA	(f) Ch(k=10) +UPGMA

Fig.2. Phylogenetic trees for 16S rRNA sequences（57 sequences）via Ch (k=4, 6, 7, 8, 9, 10)

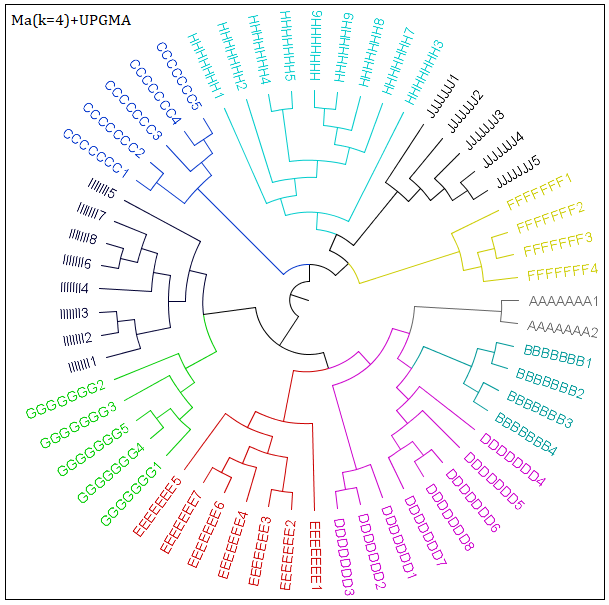

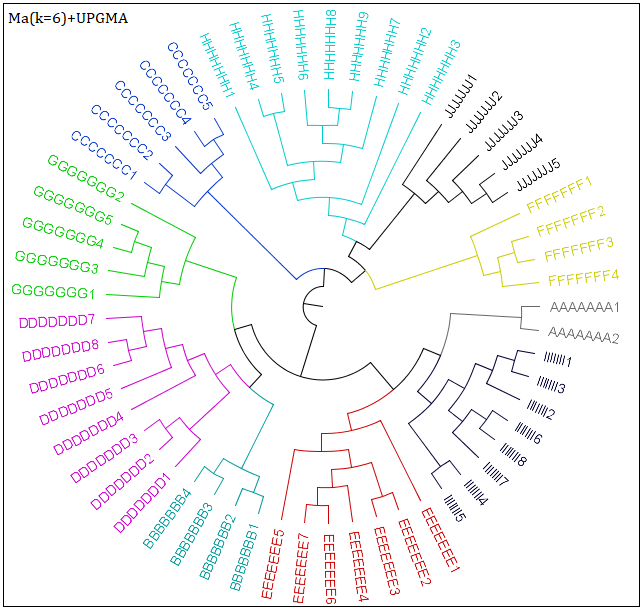

(a) Ma(k=4) +UPGMA	(b) Ma(k=6) +UPGMA

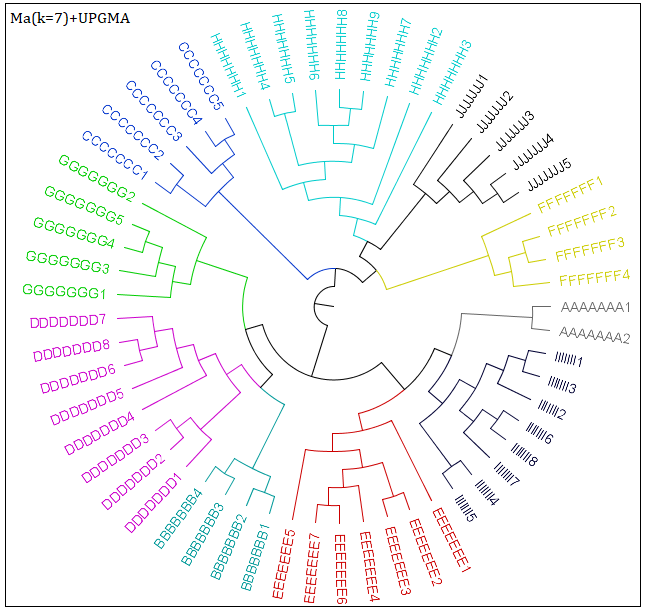

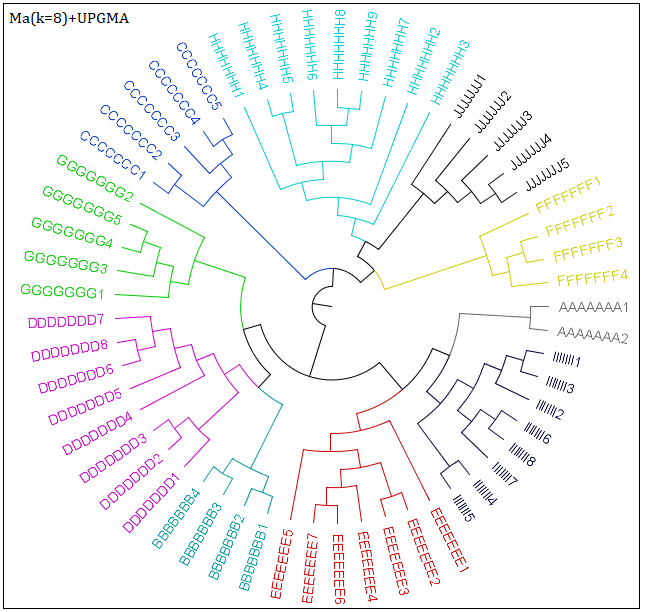

(c) Ma(k=7) +UPGMA	(d) Ma(k=8) +UPGMA

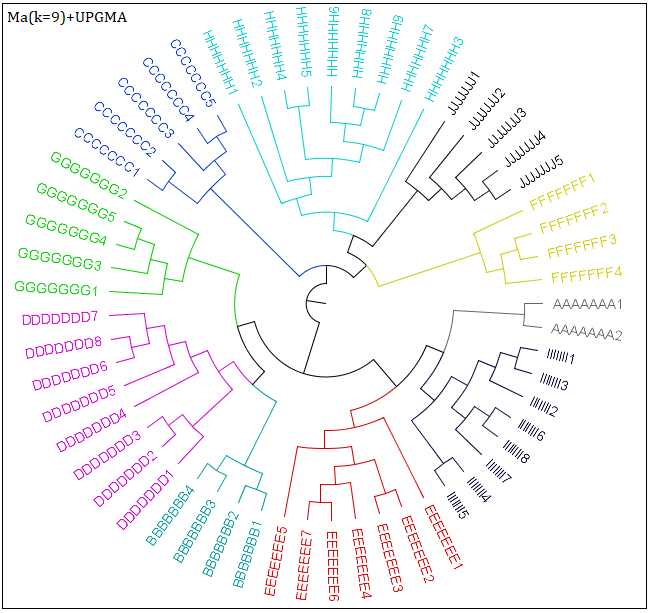

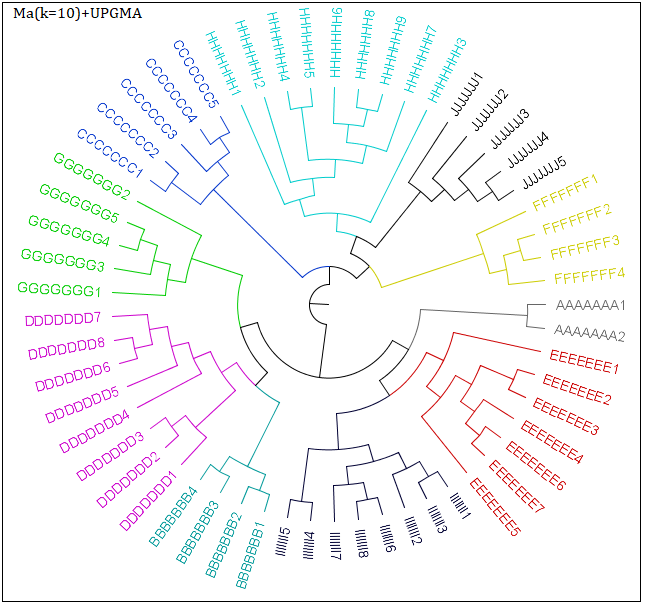

(e) Ma(k=9) +UPGMA	(f) Ma(k=10) +UPGMA

Fig.3. Phylogenetic trees for 16S rRNA sequences（57 sequences）via Ma (k=4, 6, 7, 8, 9, 10)

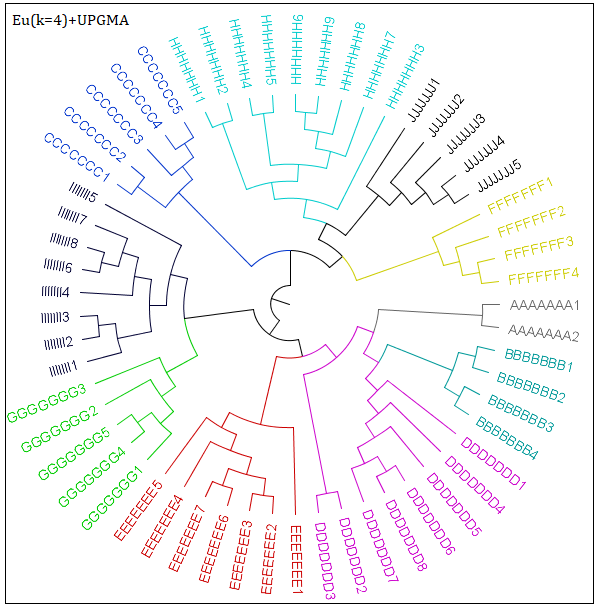

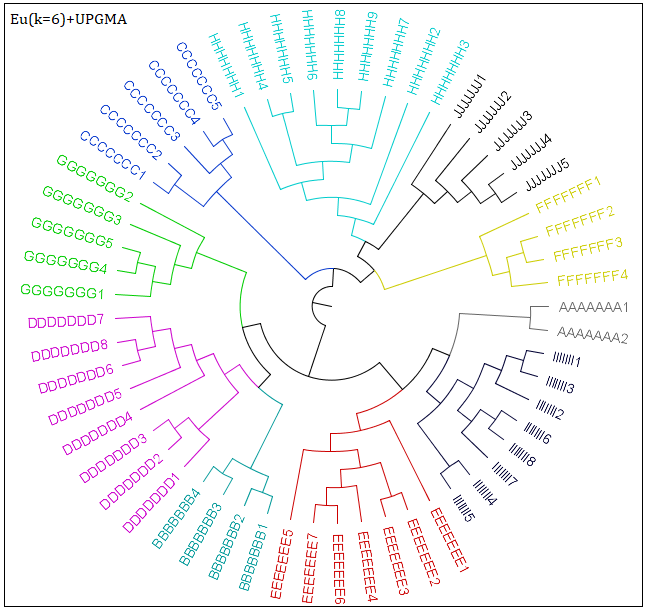

(a) Eu(k=4) +UPGMA	(b) Eu(k=6) +UPGMA

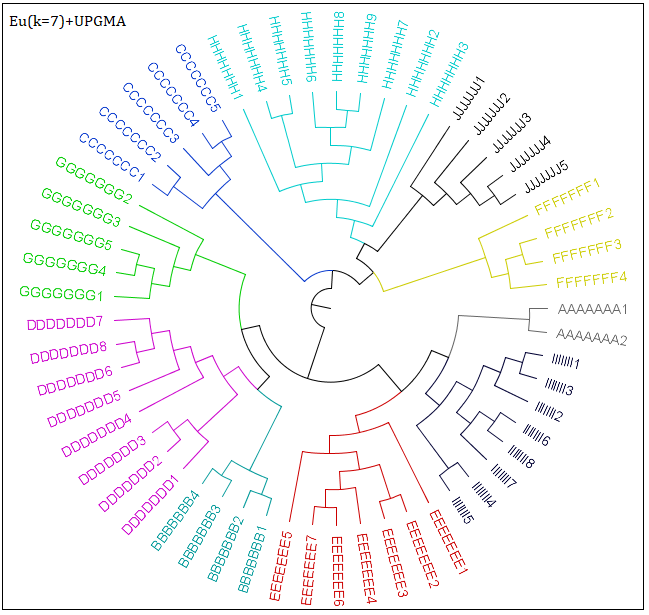

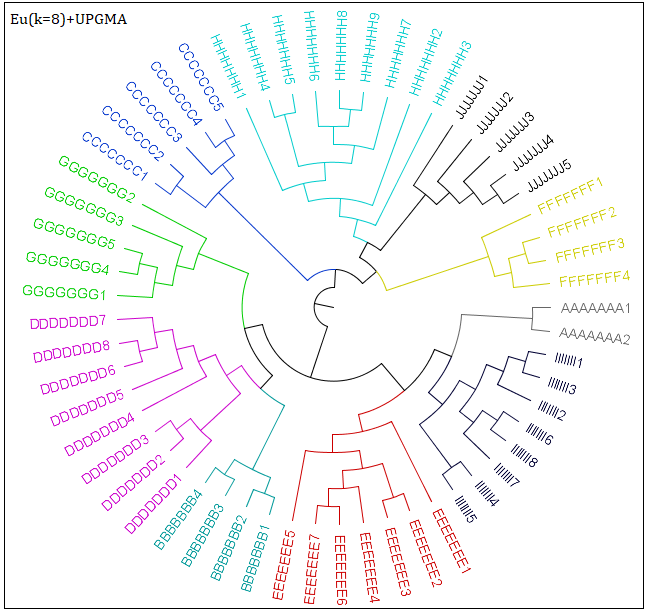

(c) Eu(k=7) +UPGMA	(d) Eu(k=8) +UPGMA

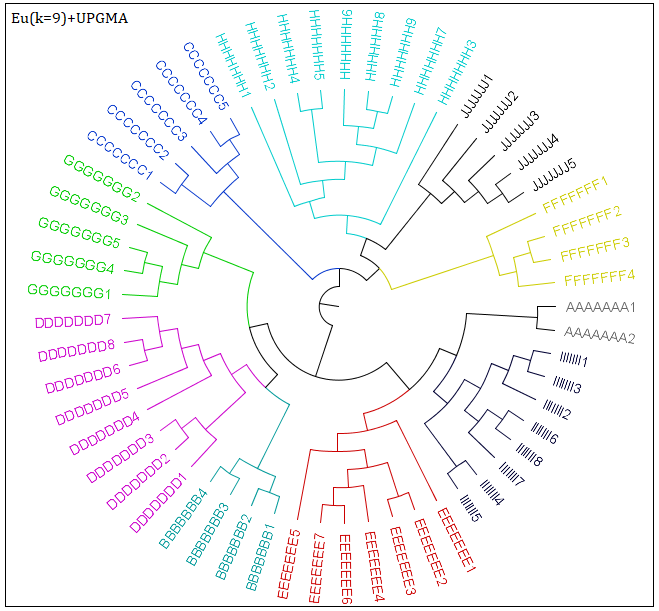

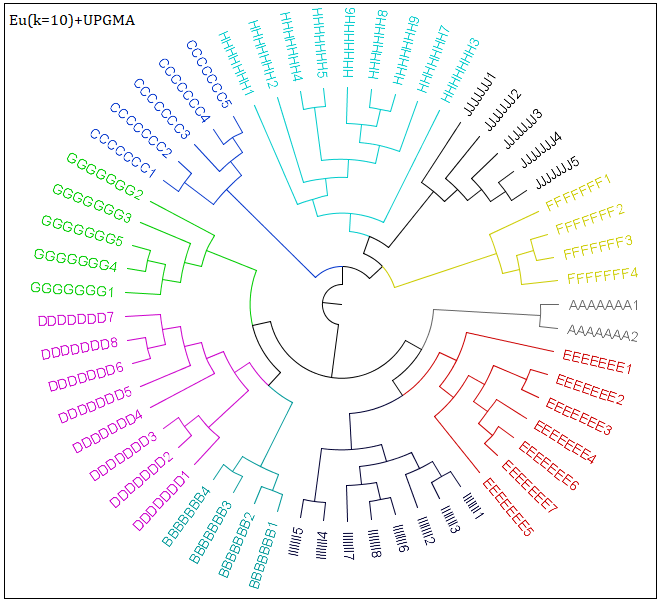

(e) Eu(k=9) +UPGMA	(f) Eu(k=10) +UPGMA

Fig.4. Phylogenetic trees for 16S rRNA sequences（57 sequences）via Eu (k=4, 6, 7, 8, 9, 10)

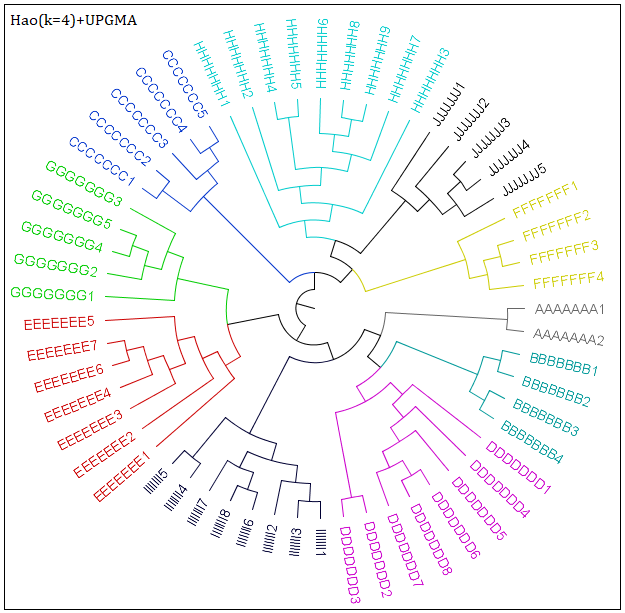

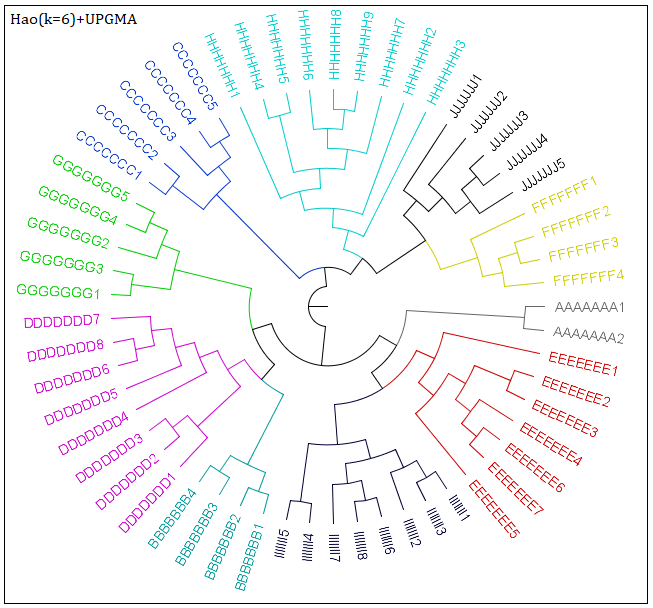

(a) Hao(k=4) +UPGMA	(b) Hao(k=6) +UPGMA

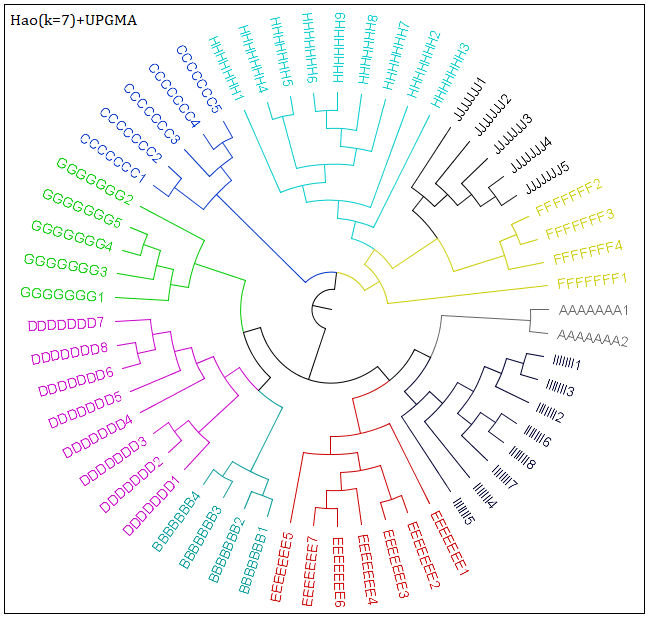

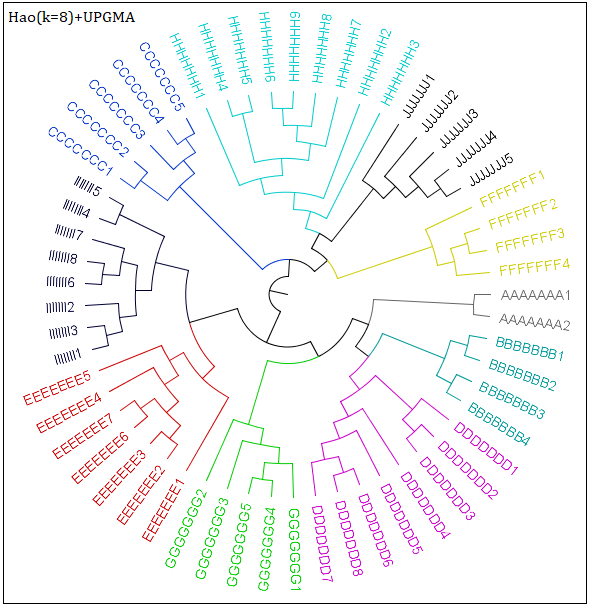

(c) Hao(k=7) +UPGMA	(d) Hao(k=8) +UPGMA

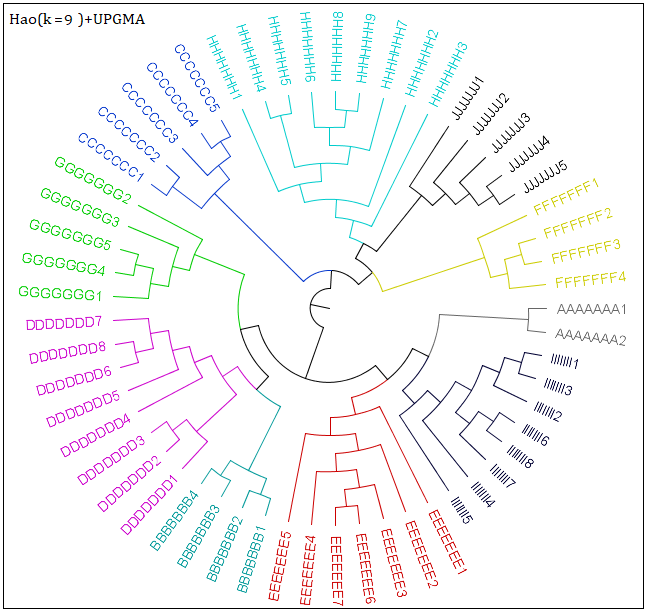

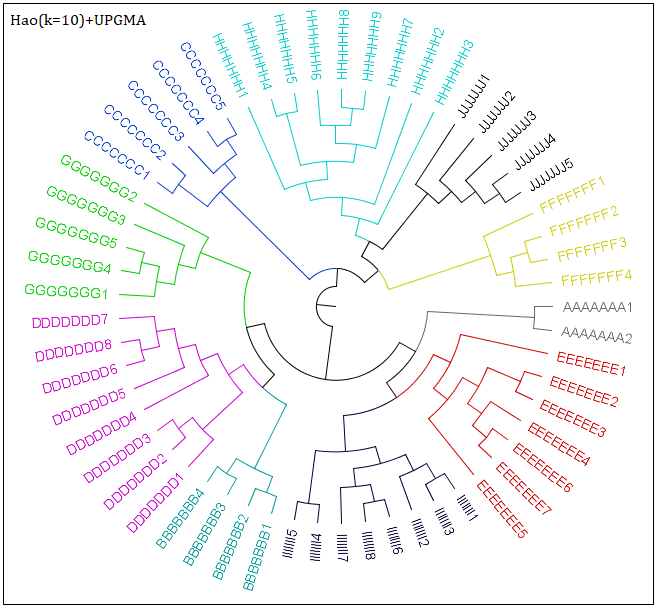

(e) Hao(k=9) +UPGMA	(f) Hao(k=10) +UPGMA

Fig.5. Phylogenetic trees for 16S rRNA sequences（57 sequences）via Hao (k=4, 6, 7, 8, 9, 10)

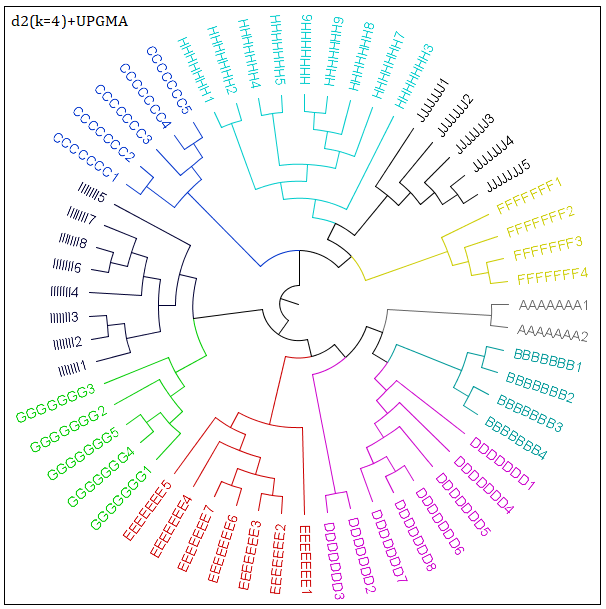

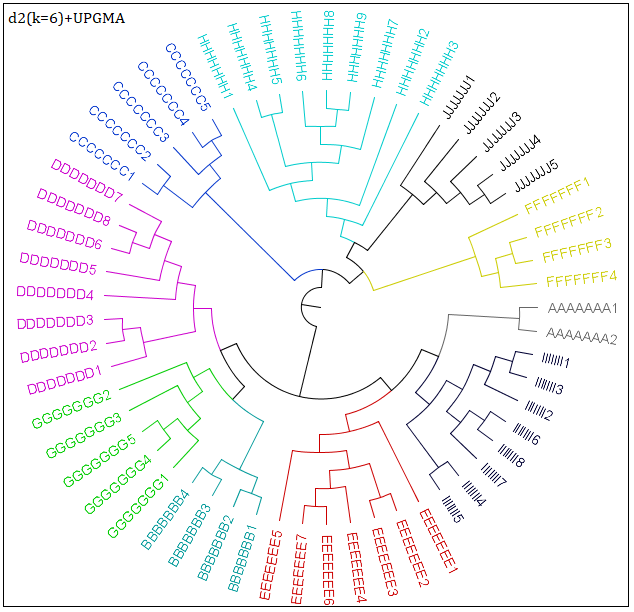

(a) d2(k=4) +UPGMA	(b) d2(k=6) +UPGMA

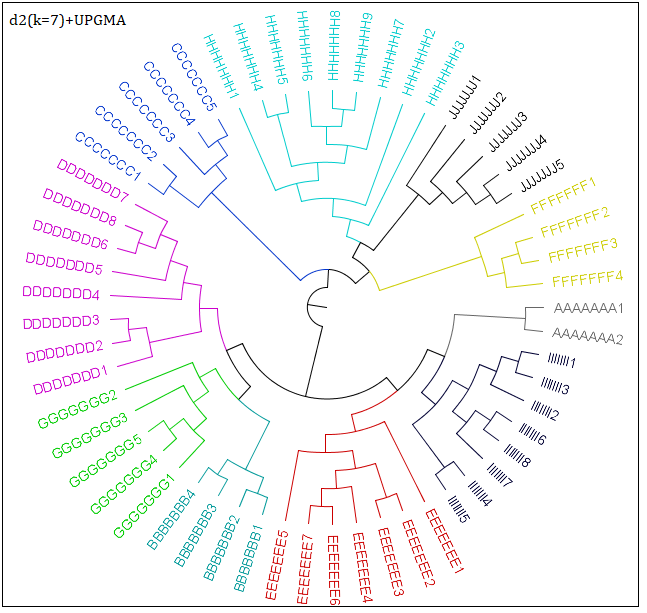

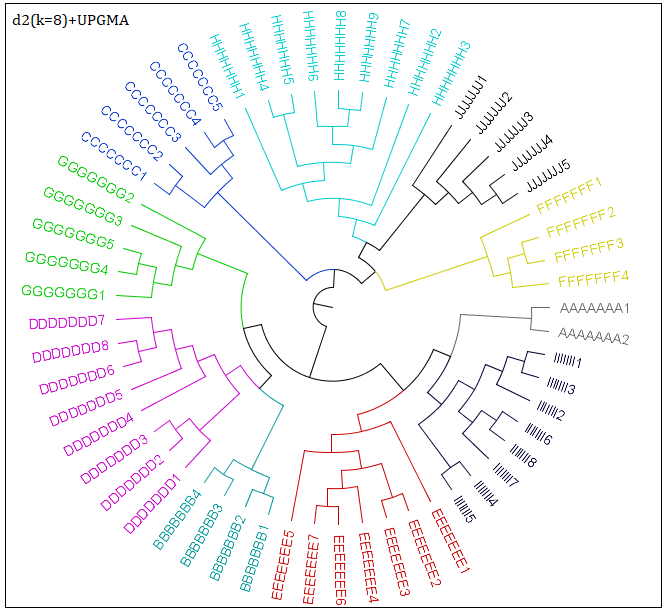

(c) d2(k=7) +UPGMA	(d) d2(k=8) +UPGMA

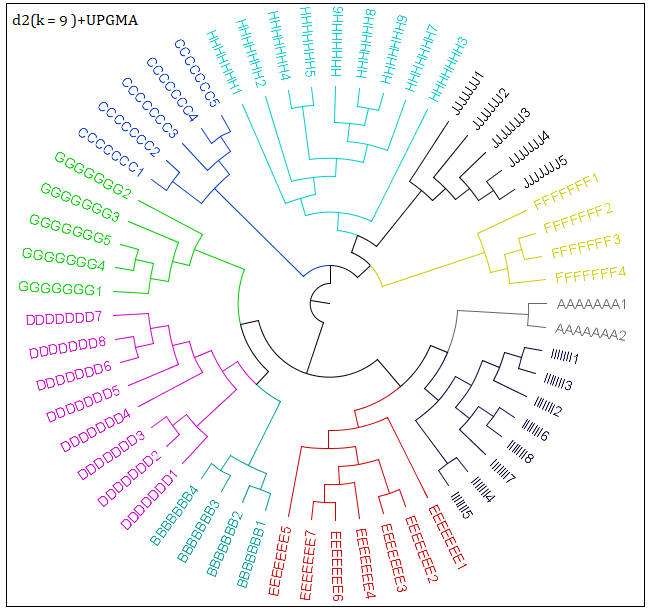

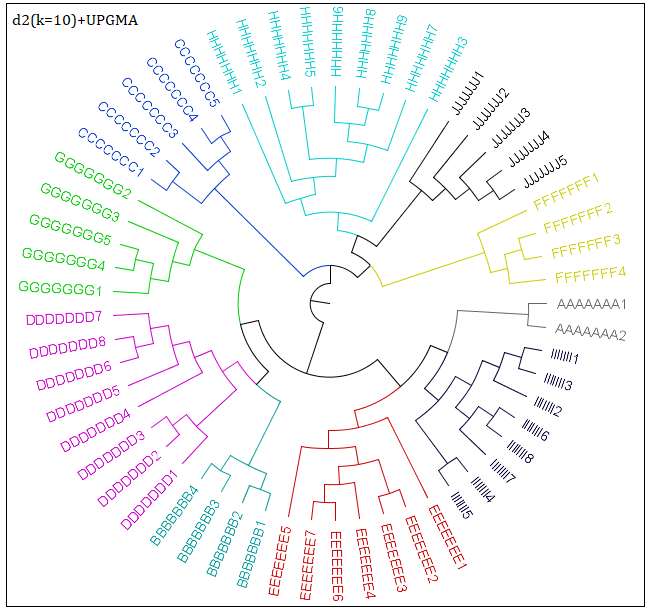

(e) d2(k=9) +UPGMA	(f) d2(k=10) +UPGMA

Fig.6. Phylogenetic trees for 16S rRNA sequences（57 sequences）via d2 (k=4, 6, 7, 8, 9, 10)

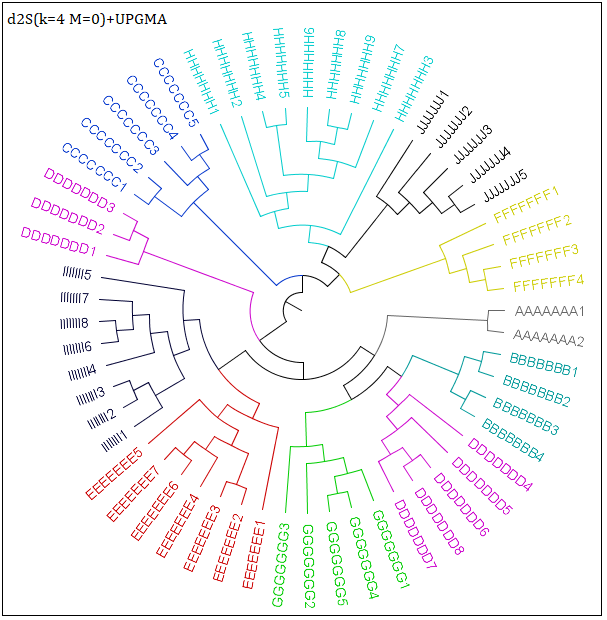

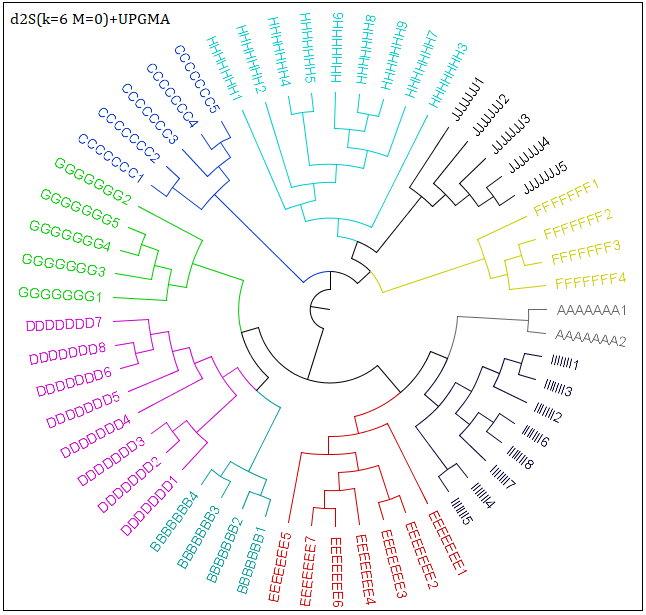

(a) d2S(k=4, M=0) +UPGMA	(b) d2S(k=6, M=0) +UPGMA

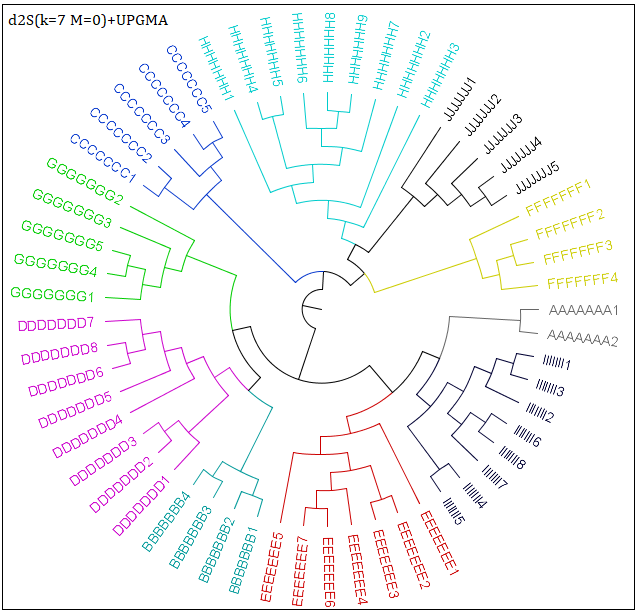

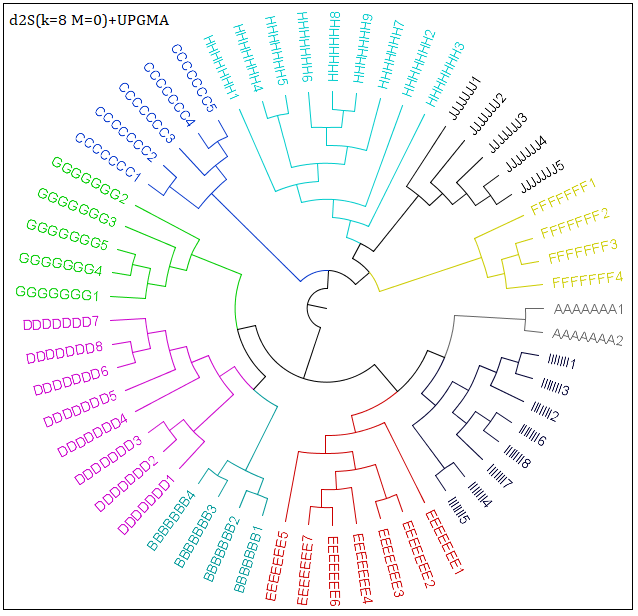

(c) d2S(k=7, M=0) +UPGMA	(d) d2S(k=8, M=0) +UPGMA

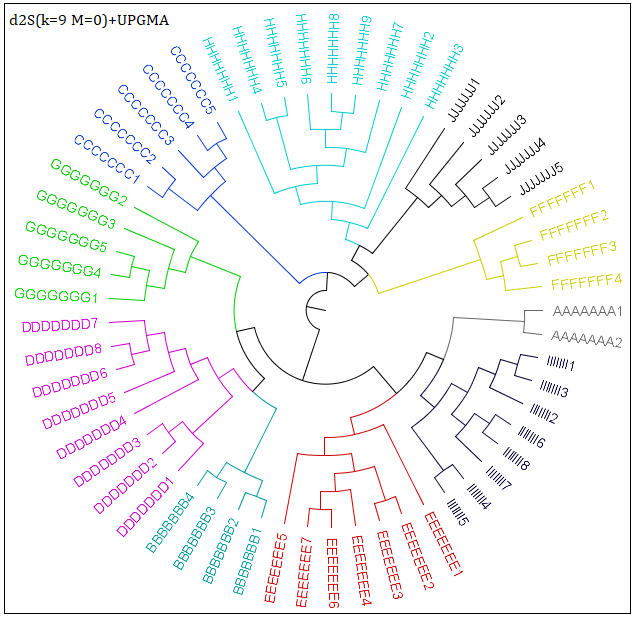

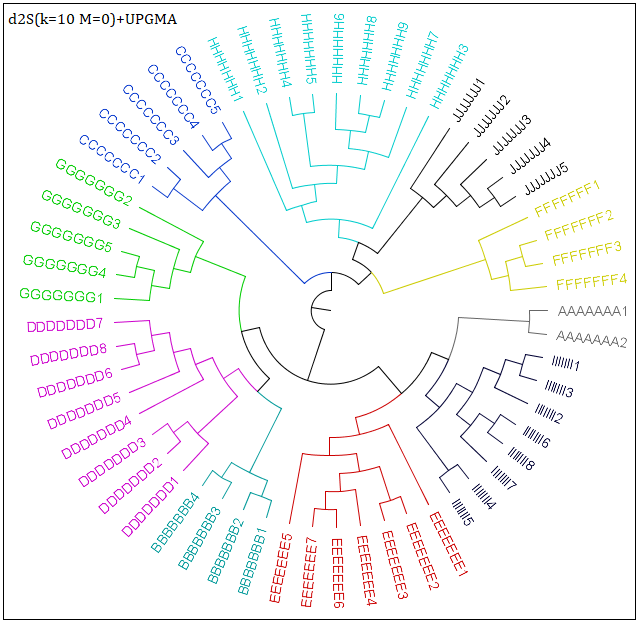

(e) d2S(k=9, M=0) +UPGMA	(f) d2S(k=10, M=0) +UPGMA

Fig.7. Phylogenetic trees for 16S rRNA sequences（57 sequences）via d2S (k=4, 6, 7, 8, 9, 10, M=0)

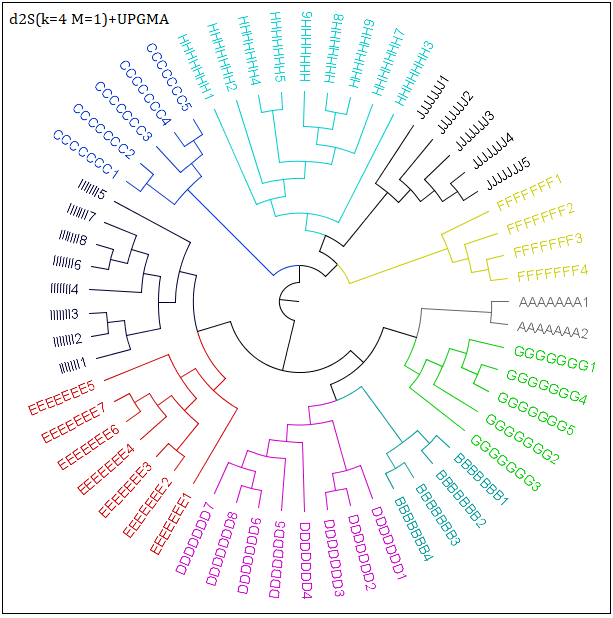

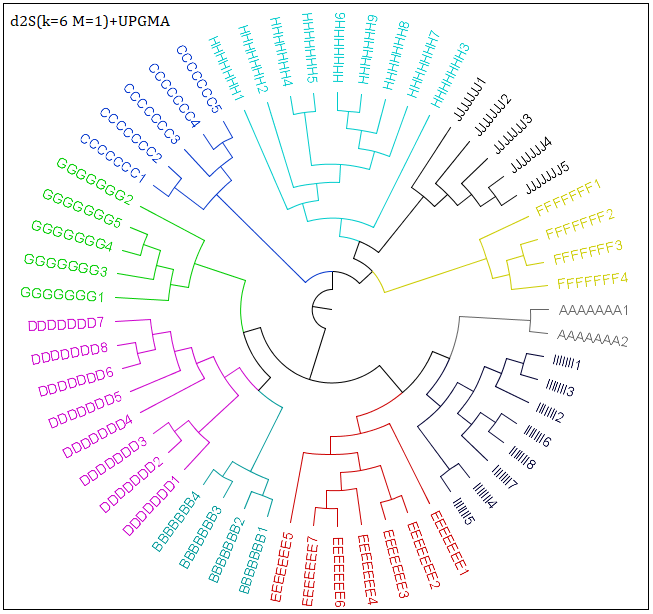

(a) d2S(k=4, M=1) +UPGMA	(b) d2S(k=6, M=1) +UPGMA

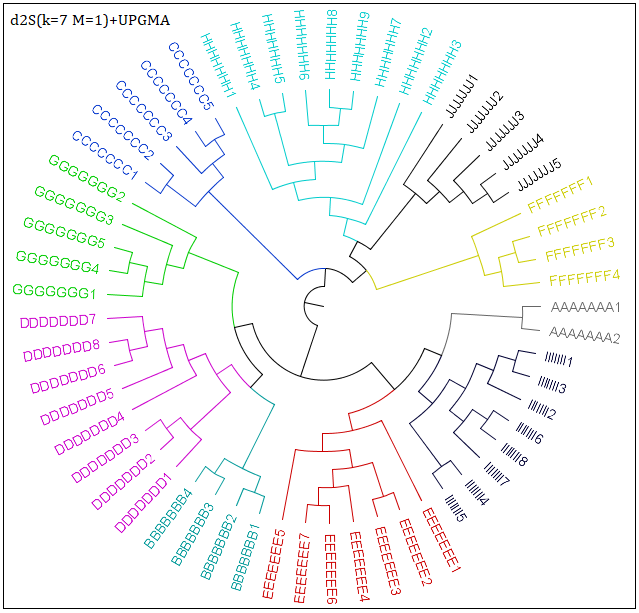

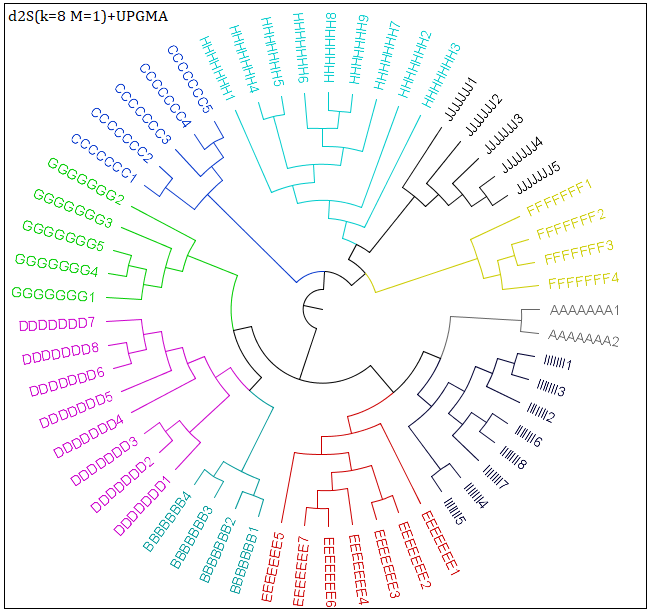

(c) d2S(k=7, M=1) +UPGMA	(d) d2S(k=8, M=1) +UPGMA

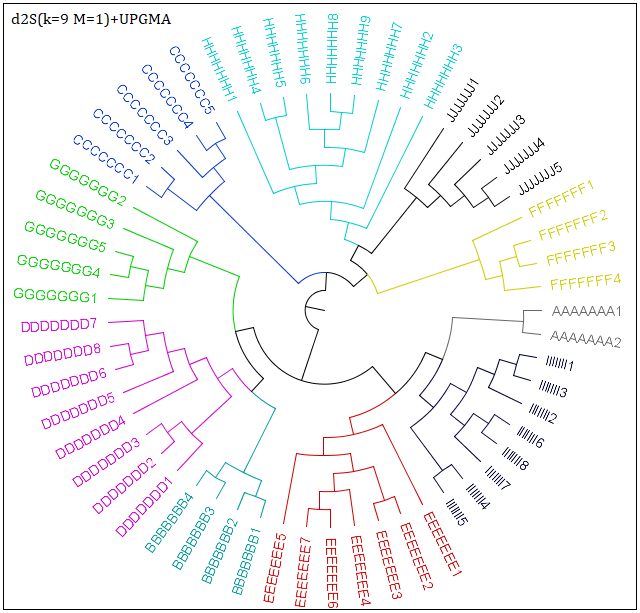

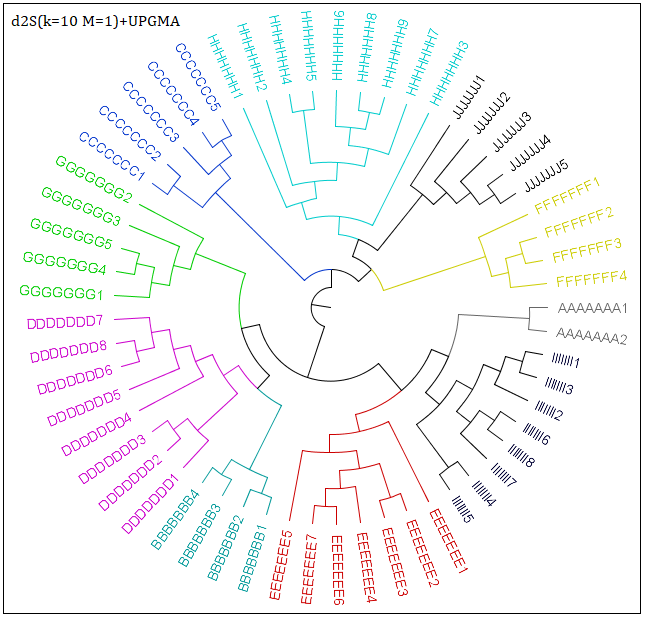

(e) d2S(k=9, M=1) +UPGMA	(f) d2S(k=10, M=1) +UPGMA

Fig.8. Phylogenetic trees for 16S rRNA sequences（57 sequences）via d2S (k=4, 6, 7, 8, 9, 10, M=1)

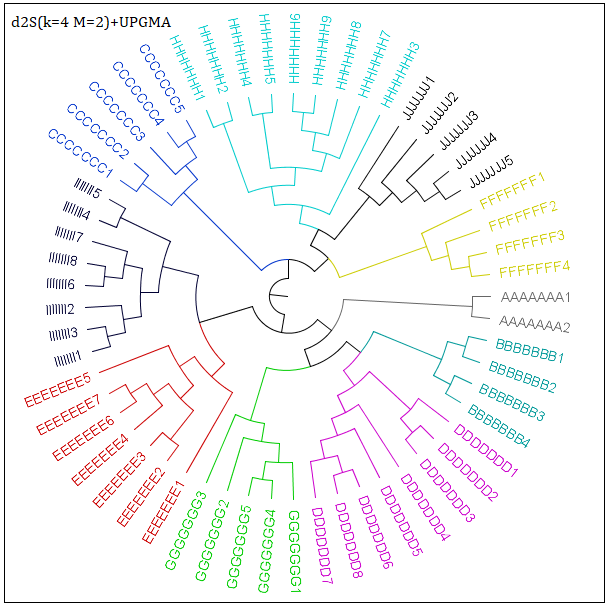

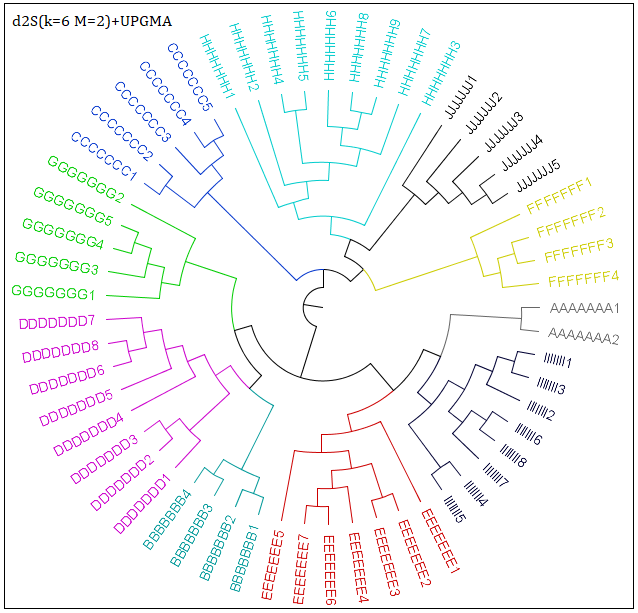

(a) d2S(k=4, M=2) +UPGMA	(b) d2S(k=6, M=2) +UPGMA

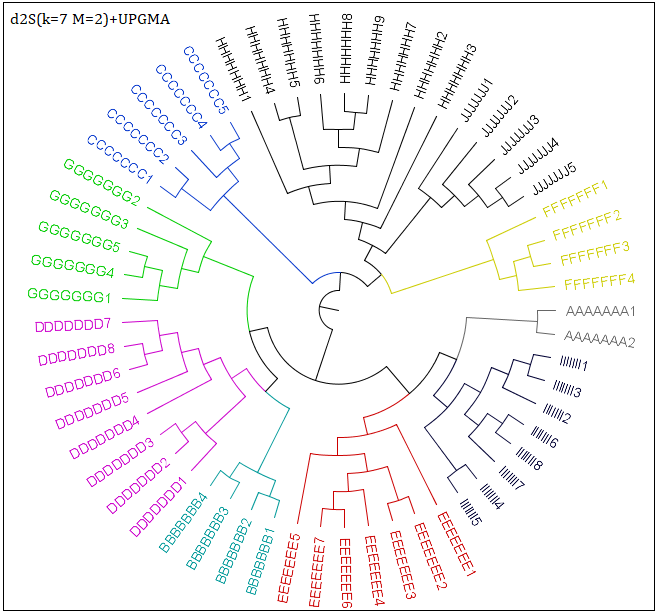

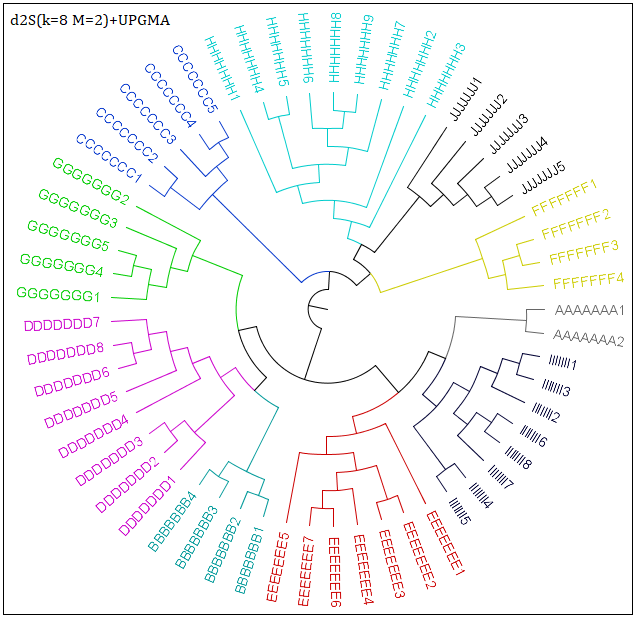

(c) d2S(k=7, M=2) +UPGMA	(d) d2S(k=8, M=2) +UPGMA

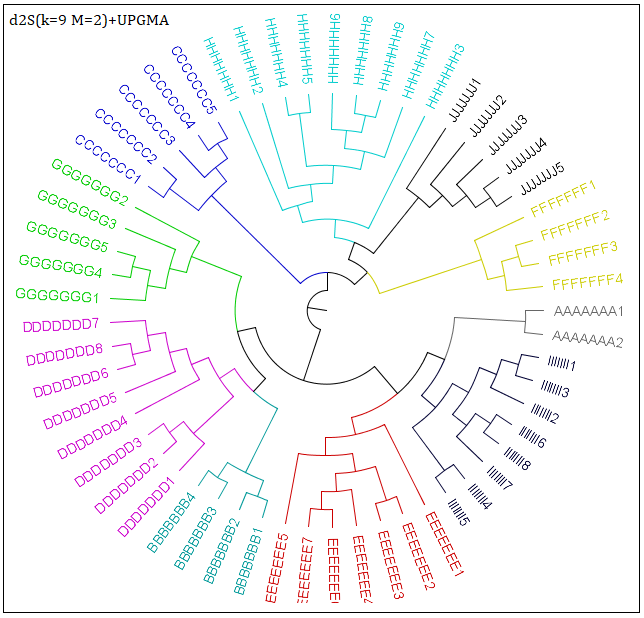

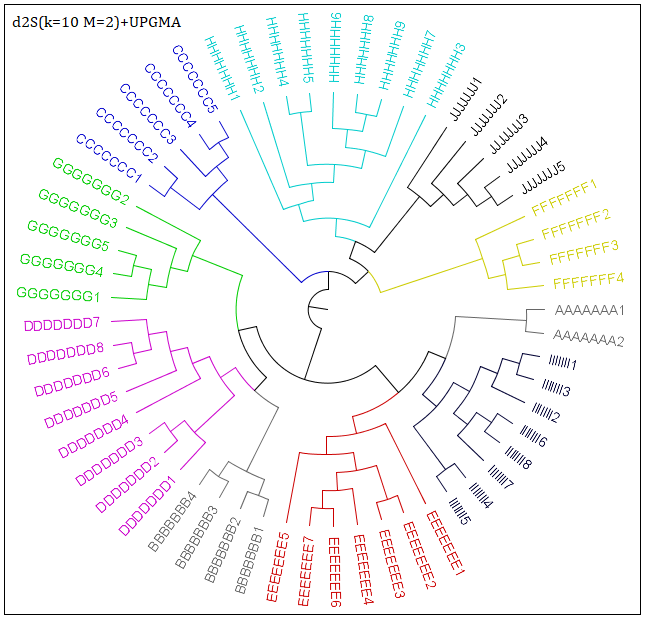

(e) d2S(k=9, M=2) +UPGMA	(f) d2S(k=10, M=2) +UPGMA

Fig.9. Phylogenetic trees for 16S rRNA sequences（57 sequences）via d2S (k=4, 6, 7, 8, 9, 10, M=2)

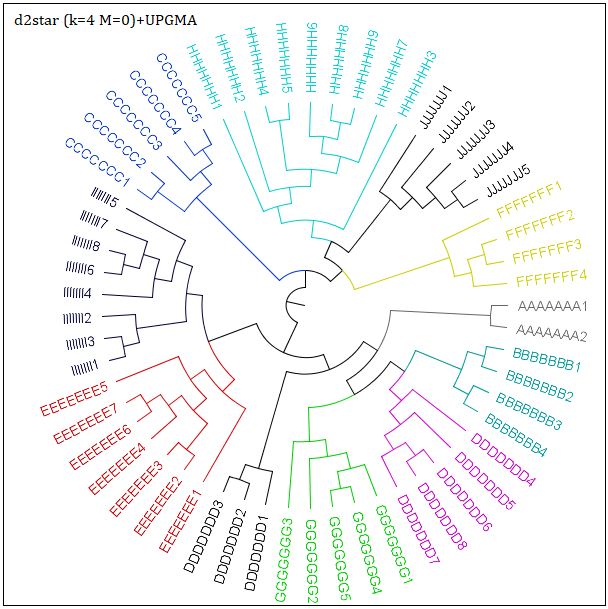

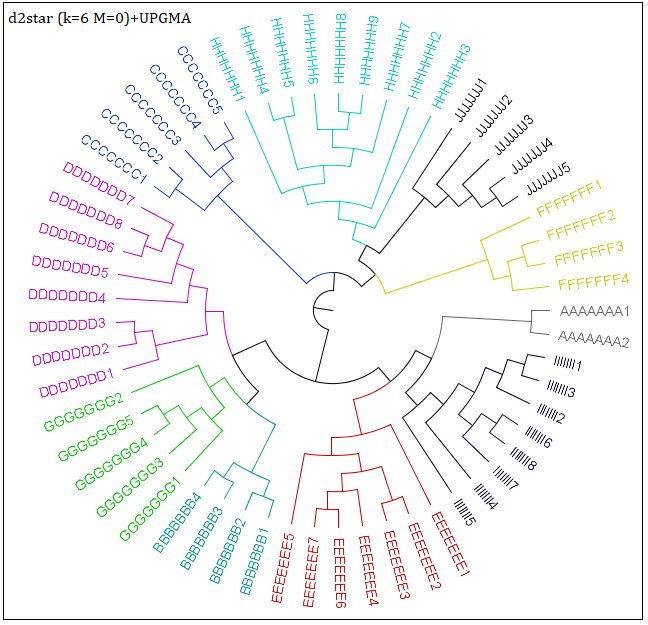

(a) d2star(k=4, M=0) +UPGMA	(b) d2star(k=6, M=0) +UPGMA

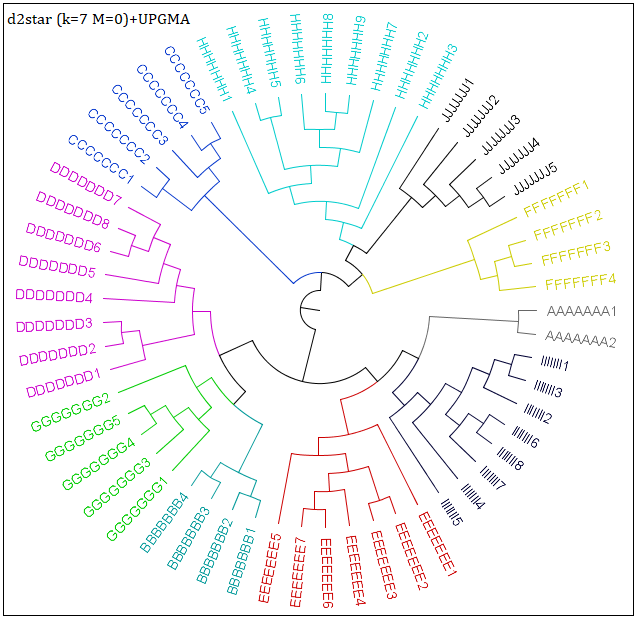

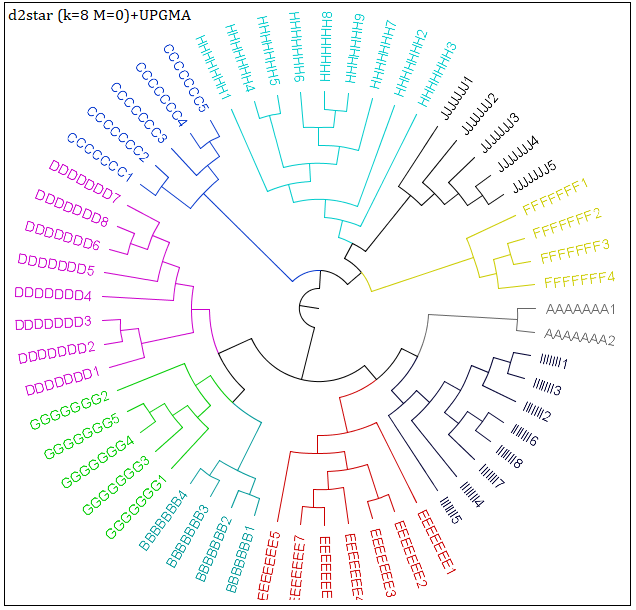

(c) d2star(k=7, M=0) +UPGMA	(d) d2star(k=8, M=0) +UPGMA

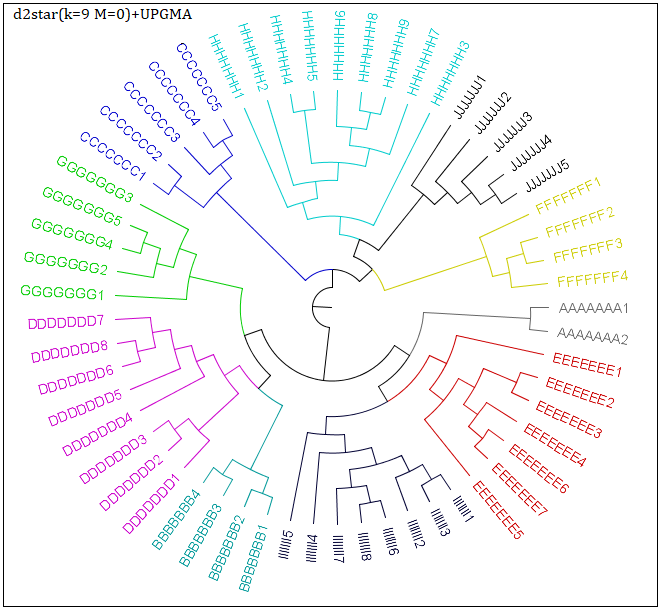

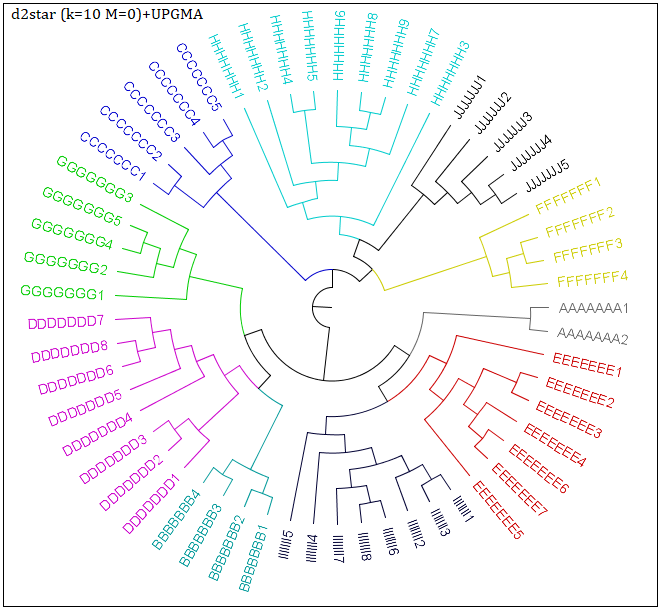

(e) d2star(k=9, M=0) +UPGMA	(f) d2star(k=10, M=0) +UPGMA

Fig.10. Phylogenetic trees for 16S rRNA sequences（57 sequences）via d2star (k=4, 6, 7, 8, 9, 10, M=0)

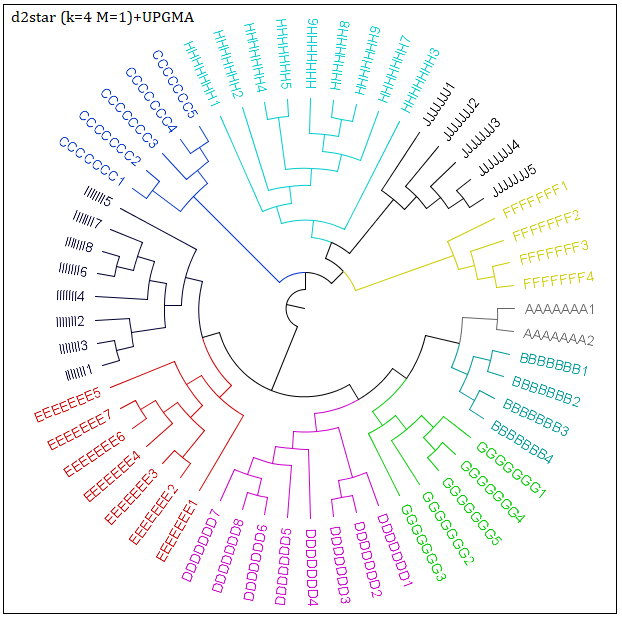

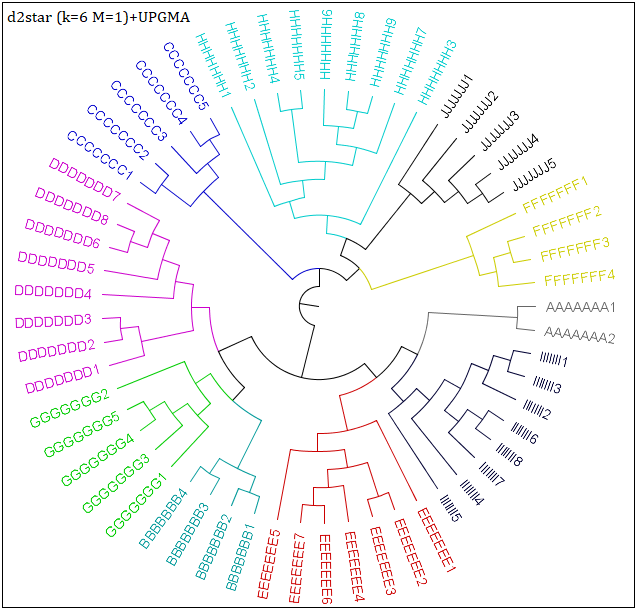

(a) d2star(k=4, M=1) +UPGMA	(b) d2star(k=6, M=1) +UPGMA

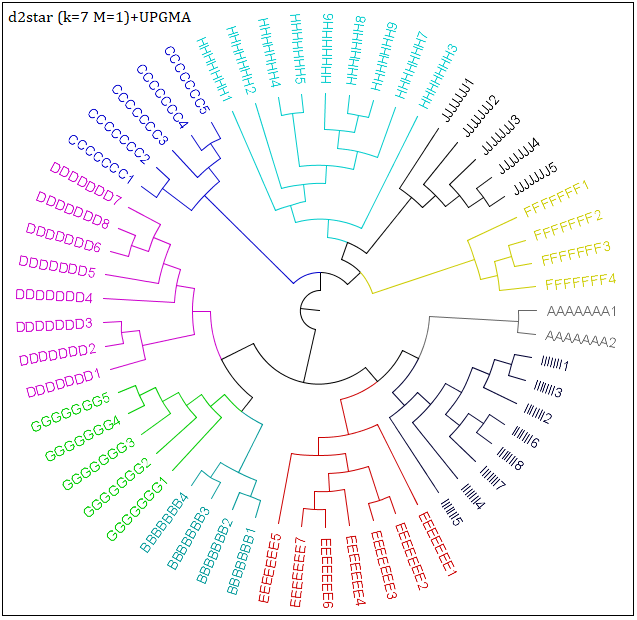

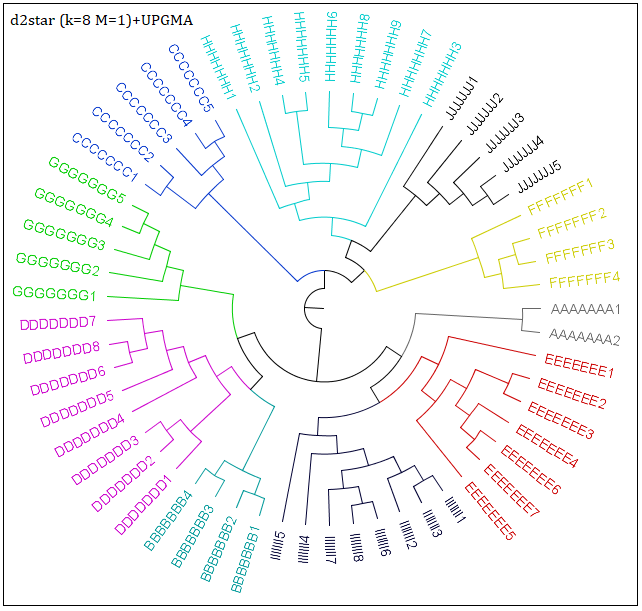

(c) d2star(k=7, M=1) +UPGMA	(d) d2star(k=8, M=1) +UPGMA

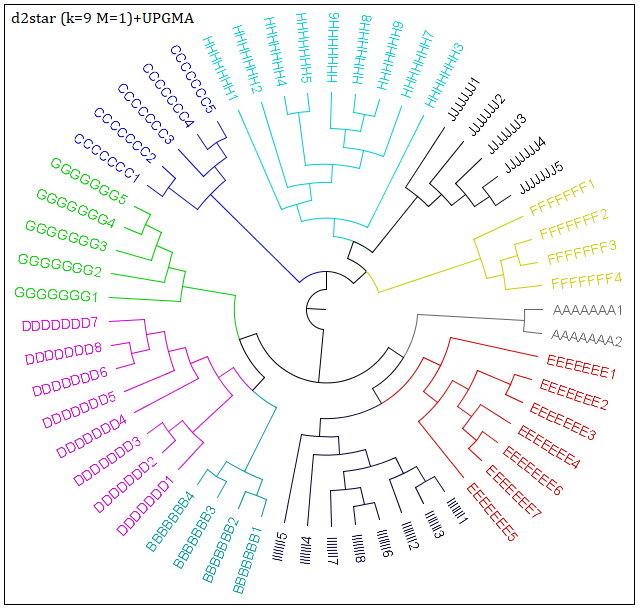

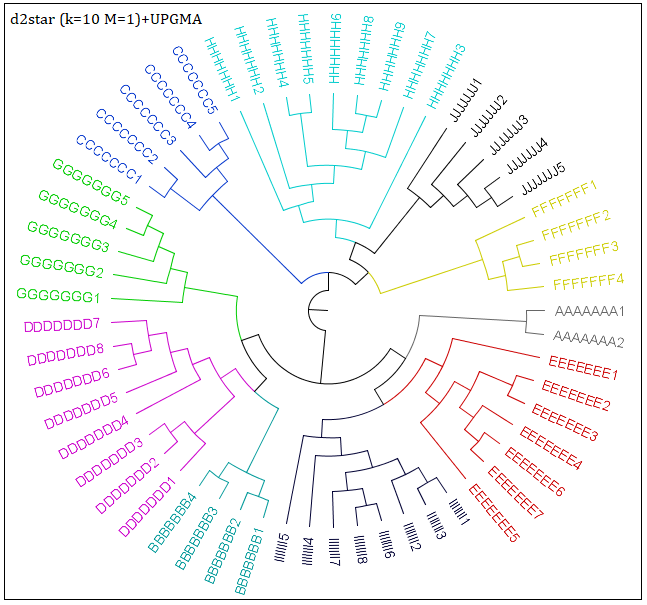

(e) d2star(k=9, M=1) +UPGMA	(f) d2star(k=10, M=1) +UPGMA

Fig.11. Phylogenetic trees for 16S rRNA sequences（57 sequences）via d2star (k=4, 6, 7, 8, 9, 10, M=1)

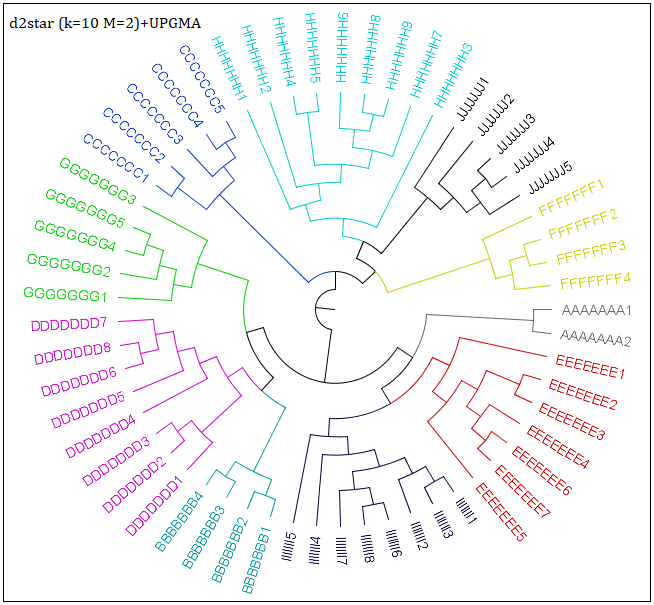

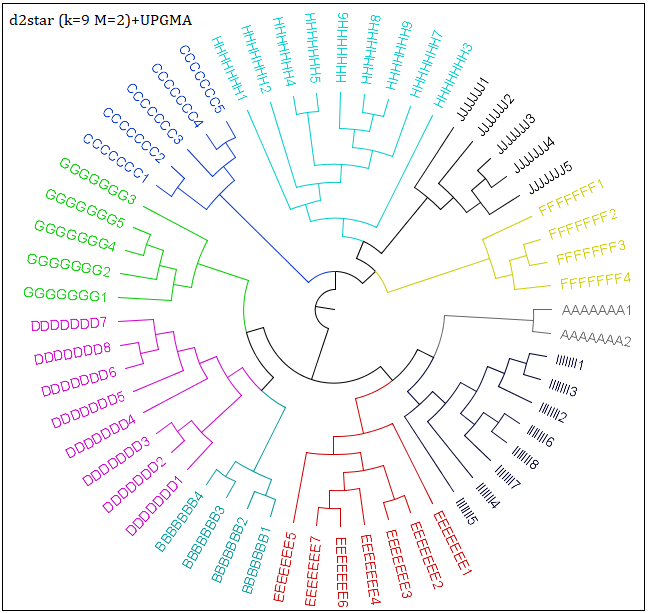

(a) d2star(k=4, M=2) +UPGMA	(b) d2star(k=6, M=2) +UPGMA

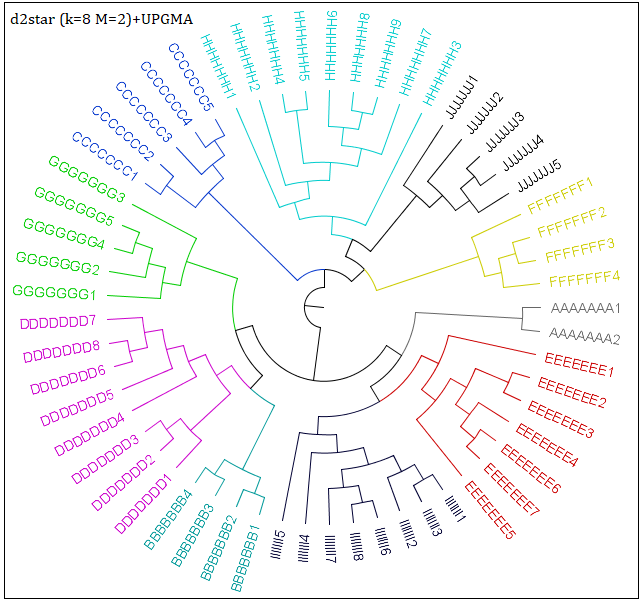

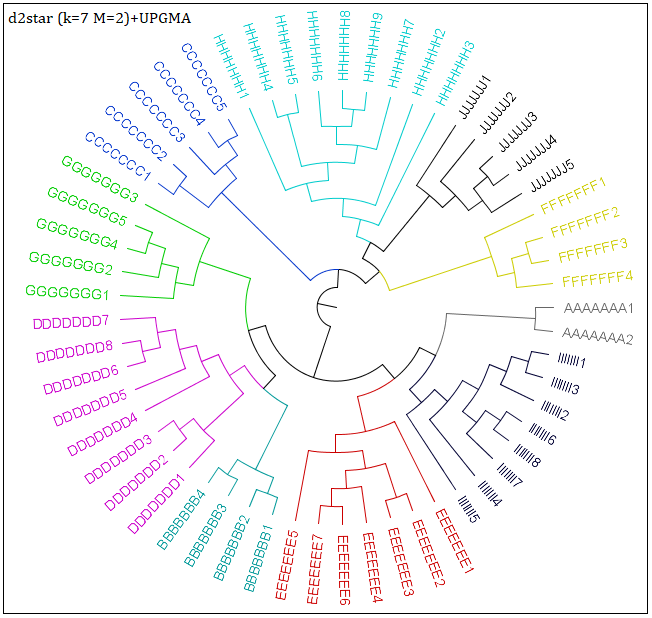

(c) d2star(k=7, M=2) +UPGMA	(d) d2star(k=8, M=2) +UPGMA

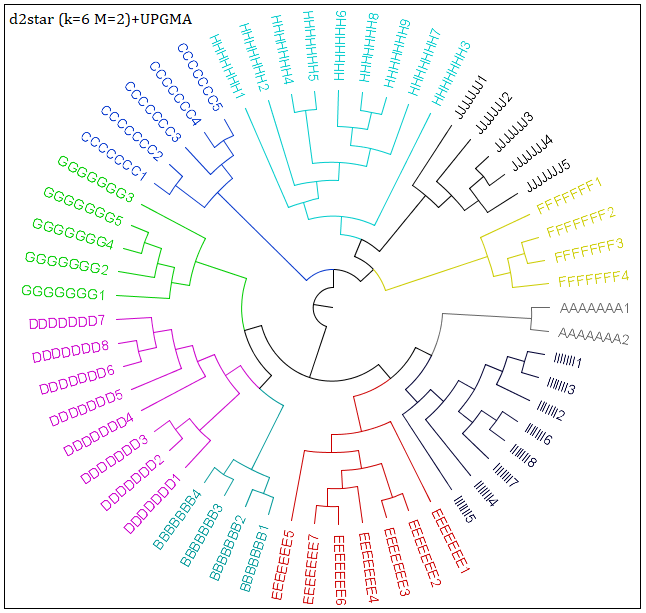

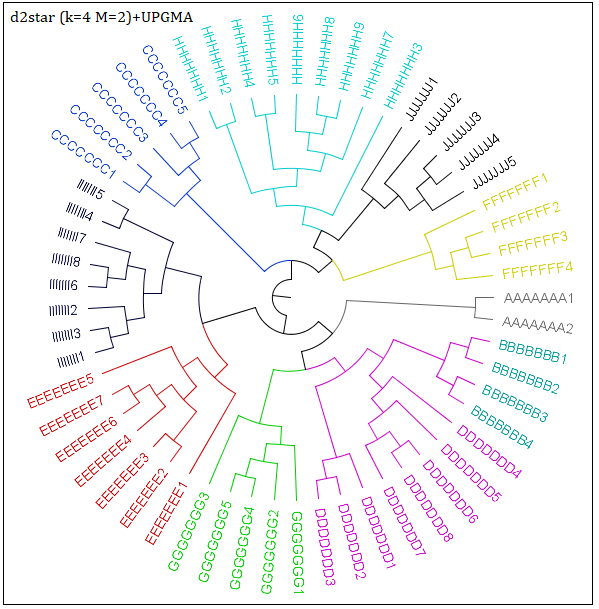

(e) d2star(k=9, M=2) +UPGMA	(f) d2star(k=10, M=2) +UPGMA

Fig.12. Phylogenetic trees for 16S rRNA sequences（57 sequences）via d2star (k=4, 6, 7, 8, 9, 10, M=2)
